# Revealing multi-scale population structure in large cohorts

**DOI:** 10.1101/423632

**Authors:** Alex Diaz-Papkovich, Luke Anderson-Trocmé, Simon Gravel

## Abstract

Genetic structure in large cohorts results from technical, sampling and demographic variation. Visualisation is therefore a first step in most genomic analyses. However, existing data exploration methods struggle with unbalanced sampling and the many scales of population structure. We investigate an approach to dimension reduction of genomic data that combines principal components analysis (PCA) with uniform manifold approximation and projection (UMAP) to succinctly illustrate population structure in large cohorts and capture their relationships on local and global scales. Using data from large-scale genomic datasets, we demonstrate that PCA-UMAP effectively clusters closely related individuals while placing them in a global continuum of genetic variation. This approach reveals previously overlooked subpopulations within the American Hispanic population and fine-scale relationships between geography, genotypes, and phenotypes in the UK population. This opens new lines of investigation for demographic research and statistical genetics. Given its small computational cost, PCA-UMAP also provides a general-purpose approach to exploratory analysis in population-scale datasets.

**Author summary:** Because of geographic isolation, individuals tend to be more genetically related to people living nearby than to people living far. This is an example of population structure, a situation where a large population contains subgroups that share more than the average amount of DNA. This structure can tell us about human history, and it can also have a large effect on medical studies. We use a newly developed method (UMAP) to visualize population structure from three genomic datasets. Using genotype data alone, we reveal numerous subgroups related to ancestry and correlated with traits such as white blood cell count, height, and FEV1, a measure used to detect airway obstruction. We demonstrate that UMAP reveals previously unobserved patterns and fine-scale structure. We show that visualizations work especially well in large datasets containing populations with diverse backgrounds, which are rapidly becoming more common, and that unlike other visualization methods, we can preserve intuitive connections between populations that reflect their shared ancestries. The combination of these results and the effectiveness of the strategy on large and diverse datasets make this an important approach for exploratory analysis for geneticists studying ancestral events and phenotype distributions.

## Introduction

Questions in medicine, anthropology, and related fields hinge on interpreting the deluge of genomic data provided by modern high-throughput sequencing technologies. Because genomic datasets are high-dimensional, their interpretation requires statistical methods that can comprehensively condense information in a manner that is understandable to researchers and minimizes the amount of data that is sacrificed. Both model-based and model-agnostic approaches to summarize data have played important roles in shaping our understanding of the evolution of our species (1).

Here we will focus on nonparametric approaches to visualize relatedness patterns among individuals within populations. If we consider unphased single nucleotide polymorphism (SNP) data, an individual genome can be represented as a sequence of integers corresponding to the number of derived alleles carried by the individual at each of the *L* SNPs for which genotypes are available, with *L* typically larger than 100, 000. Since each individual is represented as an *L*-dimensional vector, dimension reduction methods are needed to visualize the data.

Principal component analysis (PCA) is often the first dimensional reduction tool used for genomic data. It identifies and ranks directions in genotype space that explain most-to-least variance among individuals. Positions of individuals along directions of highest variance can then be used to summarize individual genotypes. PCA coordinates have natural genealogical interpretations in terms of times to a most recent common ancestor (TMRCA) (2), and are used empirically to reveal admixture (3), continuous isolation-by-distance (4, 5), as well as technical artefacts. PCA coordinates are particularly well-suited to correct for population structure in GWAS (6).

As sample sizes increase, the amount of information encoded in the lower-variance principal components increases, and researchers typically examine multiple two-dimensional projections to get a sense of the data. While many features of the data can be identified in this manner, other features may be hidden by the projections or hard to interpret.

To display more of the high-dimensional features of the data in a two dimensional figure, we can use non-linear transformations that seek to preserve the local structure of the data. A popular method for visualization is t-distributed stochastic neighbour embedding (t-SNE)(8). Whereas PC projection is designed to capture as much variance as possible for a linear transformation, t-SNE seeks a low-dimensional representations of the data that preserves pairwise distances, using a probabilistic heuristic to weight mismatches. t-SNE has been used before to visualize SNPs(9). Using data from the 1000 Genomes Project (1KGP)(7), it groups individuals corresponding roughly to their continent of origin, with smaller ethnic sub-groups visible within the larger continental clusters(10). However, t-SNE struggles with very large datasets, when a large number of locally optimal configurations make convergence to a globally satisfying solution difficult.

Uniform Manifold Approximation and Projection (UMAP) is a new dimension reduction technique grounded in Riemannian geometry, algebraic topology, and category theory, and designed to model and preserve the high-dimensional topology of data points in the low-dimensional space(11). The assumption behind UMAP is that data are uniformly distributed on local manifolds in high-dimensional space, which can be approximated as fuzzy sets that are patched together to form a topological representation. One can then construct a low-dimensional topological representation that minimizes the differences between the two representations. With genotype data, UMAP creates a neighbourhood around each individual’s genetic coordinates and identifies a pre-selected number of neighbours to build high-dimensional manifolds. The end result is a low-dimension representation that groups genetically similar individuals together on a local scale while preserving long-range topological connections to more distantly related individuals. The method has been successfully applied to single-cell RNA sequencing datasets(12).

A common practice in dimensional reduction is to first apply PCA to reduce the number of dimensions before performing nonlinear dimensional reduction. In addition to being computationally advantageous, this discards statistical noise that can confound nonlinear approaches: Population structure arising from *n* isolated randomly-mating demes can be described by the leading *n* − 1 PCs, with the following PCs describing stochastic variation in relatedness (6). Selecting the leading PCs therefore has potential to extract meaningful population structure while filtering out stochastic noise.

We explore different strategies to pre-process the data and investigate discrete and continuous population structure patterns present in large datasets of human genotypes: the 1KGP, the Health and Retirement Study (HRS)(13), and the UK BioBank (UKBB)(14).

## Results

### Fine-scale visualization of the 1KGP dataset

The 1KGP contains genotype data of 3,450 individuals from 26 relatively distinct labeled populations(7). Figure 1 shows visualizations using PCA, t-SNE, UMAP, and PCA-UMAP (that is, UMAP with PCA pre-processing). Using UMAP and t-SNE on the genotype data presents clusters that are roughly grouped by continent, with UMAP showing a clear hierarchy of population and continental clusters, whereas t-SNE fails to assign many individuals to population clusters. Using either on the top principal components leads to more distinct population clusters and less defined continental structure (see figure S2 for PCA-tSNE). Adding more components results in progressively finer clusters until approximately 20 populations appear using 15 components; further components gradually approach results similar to using the entire genotype data (see figures S1 and S2).

**Fig. 1.**
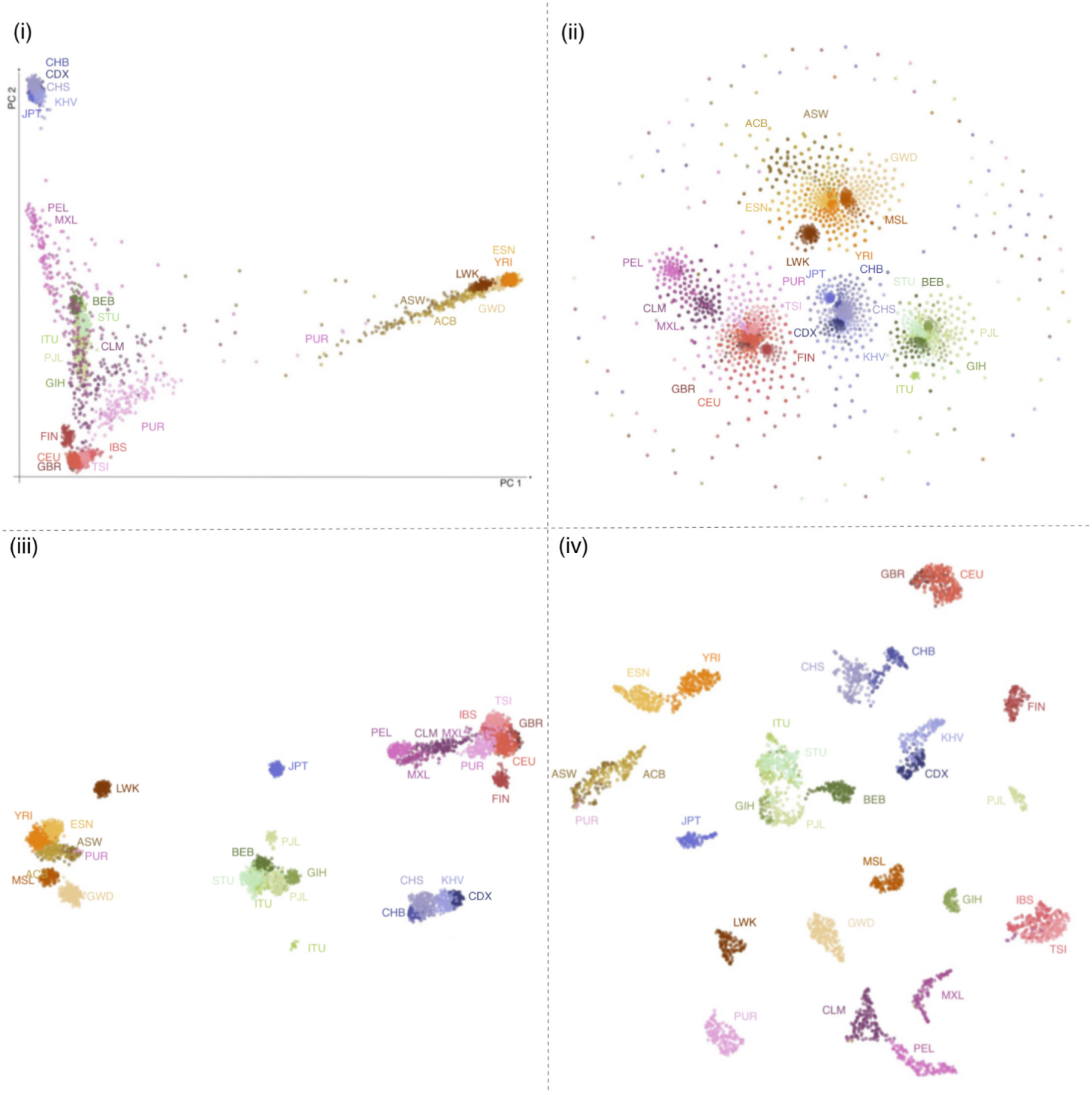
Four methods of dimension reduction of 1KGP genotype data with population labels (i) PCA maps individuals in a triangle with vertices corresponding to African, Asian/Native American, and European continental ancestry. Discarding lower-variance PCs leads to overlap of populations with no close affinity, such as Central and South American populations with South Asians. (ii) t-SNE forms groups corresponding to continents, with some overlap between European and Central and South American people. Smaller subgroups are visible within continental clusters. The cloud of peripheral points results from the method’s poor convergence. (iii) UMAP forms distinct clusters related to continent with clearly defined subgroups. Japanese, Finnish, Luhya, and some Punjabi and Telugu populations form separate clusters consistent with their population history(7). (iv) UMAP on the first 15 principal components forms fine-scale clusters for individual populations. Groups closely related by ancestry or geography, such as African Caribbean/African American, Spanish/Italian, and Kinh/Dai populations cluster together. Results using t-SNE on principal components are presented in figure S1. Axes in UMAP and t-SNE are arbitrary. ACB, African Caribbean in Barbados; ASW, African Ancestry in Southwest US; BEB, Bengali; CDX, Chinese Dai; CEU, Utah residents with Northern/Western European ancestry; CHB, Han Chinese; CHS, Southern Han Chinese; CLM, Colombian in Medellin, Colombia; ESN, Esan in Nigeria; FIN, Finnish; GBR, British in England and Scotland; GWD, Gambian; GTH, Gujarati; IBS, Iberian in Spain; ITU, Indian Telugu in the UK; JPT, Japanese; KHV, Kinh in Vietnam; LWK, Luhya in Kenya; MSL, Mende in Sierra Leone; MXL, Mexican in Los Angeles, California; PEL, Peruvian; PJL, Punjabi in Lahore, Pakistan; PUR, Puerto Rican; STU, Sri Lankan Tamil in the UK; TSI, Tuscani in Italy; YRI, Yoruba in Nigeria

Focusing on PCA-UMAP with 15 principal components (figure 1 (iv)), we also find several population clusters that reflect shared ancestries. British individuals from England and Scotland form a cluster mixed with those from Utah who claim Northern and Western European ancestry. Toscani and Iberian individuals form a group reflecting their Mediterranean heritage. African Americans in the Southwest US, African Caribbean individuals in Barbados, and some Puerto Ricans also form a cluster. The East Asian super-population forms three sub-populations split by geography: one is largely Han and Southern Han individuals, another is comprised of the Chinese Dai in southern China and the Kinh from Vietnam, and the third is the Japanese population. Looser geographical groupings include Colombians and Peruvians, and the Esan and Yoruba populations of Nigeria; both groupings appear as connected sub-clusters. The South Asian super-population also forms a loose grouping.

Only a few individuals cluster differently than the majority of individual bearing the same population label: a few Mexican individuals cluster with Spanish and Italian individuals, and a few Puerto Ricans cluster with the African Americans and African Caribbeans, likely resulting from ancestry proportions that differ from the majority. One Gambian-identified individual is present in a cluster that is otherwise entirely Mende people from Sierra Leone. Only two populations form multiple clusters: Gujarati Indians in Houston, Texas and Punjabi people in Lahore, Pakistan. This clustering is robust to, e.g., the choice of the number of PCs considered (see figure S2).

Finally, contrasting UMAP and t-SNE, we find that UMAP preserves more of the global structure of the data than t-SNE, and is more robust to choices of data pre-processing (figure S2).

### The genetic continuum of admixed populations

The 1KGP sampled individuals from relatively distinct population groups across the world, which makes the data particularly easy to cluster. Most medical cohorts comprise larger numbers of individuals sampled across extended geographical areas.

For example, the HRS contains genotype data of 12,454 American individuals across all 50 states who have provided racial identity (10,434 White, 1,652 Black, 368 Other) as well as whether they identify as Hispanic (1,203 total) and, if so, whether they identify as Mexican-American (705 total)(13). We crossed these three variables to form a composite self-reported ethnicity resulting in 10 categories (e.g. White Hispanic Mexican-American), and considered birth regions based on the 10 census regions and divisions used by the US Census Bureau. Admixture proportions for each individual were estimated in (15) by assuming ancestral African, Asian/Native American, and European populations using RFMIX (16). We have scaled these three proportions to values between 0 and 255, to color individual points by their estimated admixture represented by RGB where red, green, and blue respectively correspond to African, European, and Asian/Native American ancestry. Using the first 10 principal components and UMAP, we demonstrate projections that present a collection of sub-populations and a continuum of genetic variation.

The HRS forms several large groupings and clusters, reflecting both ethnicity (figure S33) and admixture proportions (figure 2). Gradients in admixture proportion are visible within the predominantly Hispanic cluster, but not within the predominantly Black cluster, perhaps because the variance in ancestry proportions is greater among Hispanics. The “White Not Hispanic” (WNH) group forms several interconnected clusters, and these do not correspond to broad geographical areas (figure S34). The clarity of the interconnected clusters varies by parameterization, but they consistently form a large, roughly connected group.

**Fig. 2.**
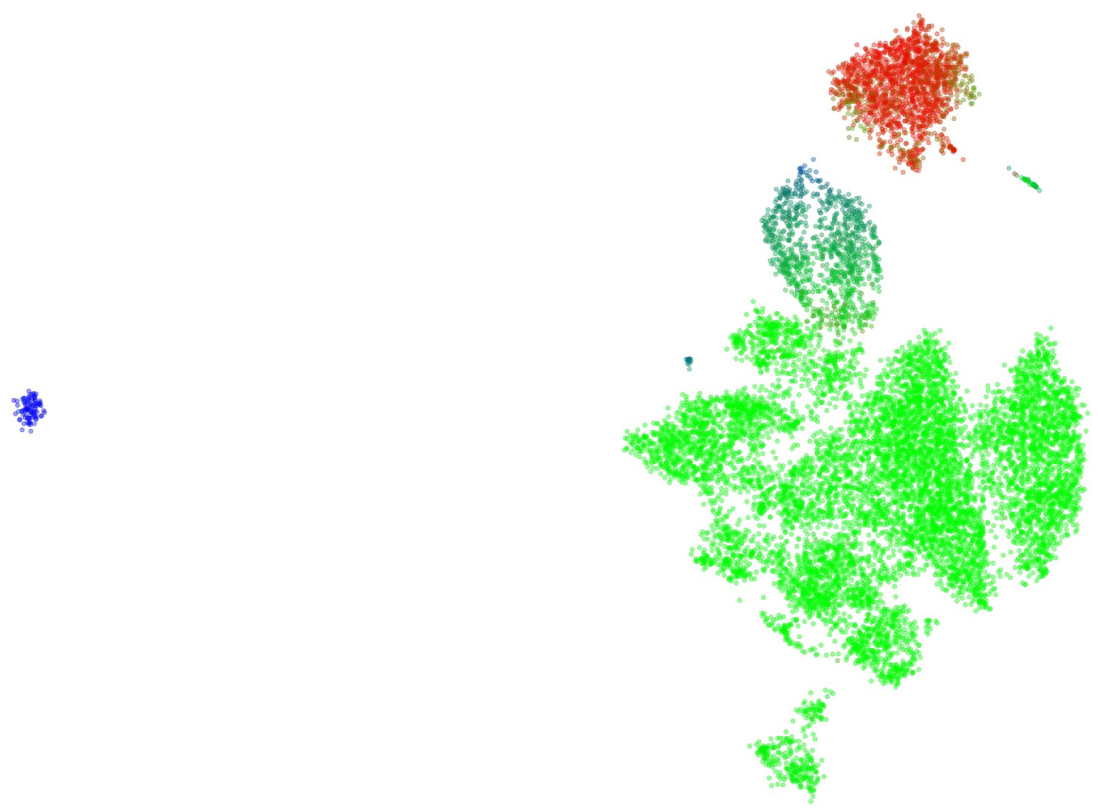
UMAP on the first 10 principal components of HRS data. Coloring individuals by estimated admixture from three ancestral populations reveals considerable diversity in the Hispanic population. This projection colored by self-identified race and Hispanic status is presented in figure S33.

To investigate possible ancestries related to populations in the 1KGP we took two approaches. In the first we generated PC axes and a UMAP embedding using UMAP and 1KGP data together (figure S36). In the second we used the PC axes and UMAP embedding generated in figure 2 and projected 1KGP data onto it (figure S35). Both approaches reveal substructure within the Hispanic cluster, groupings of Finnish individuals within the WNH groups, as well as Italian and Spanish individuals grouping near the White Hispanic population. One group of WNH individuals regularly appears at the periphery of the main cluster and does not cluster with any 1KGP populations.

### Regional patterns in the American Hispanic population

In contrast to the WNH individuals, applying PCA-UMAP to self-identified Hispanic individuals reveals clear groupings related to birth region. One separate cluster, highlighted in figure 3, consists almost entirely of individuals born in the Mountain Region of the United States. This cluster is not apparent when looking at a grid of pairwise plots of the first 8 principal components, provided in figure S37, as the signal is distributed along PCs 3, 4, and 6. Even though continental admixture patterns do correlate with UMAP position (figure S38), these do not explain the Mountain Region cluster. Individuals from 1KGP populations do not appear in the cluster when projected to the UMAP embedding. The cluster possibly comprises the Hispano population of the Southwest US, who have been present in the Mountain Region area long before the more recent immigrants from Latin America, and whose ancestry is expected to reflect both distinct Native ancestry and population-specific drift relative to other Hispanic populations. A recent preprint discusses the Mountain Region Hispanics and provides a more detailed historical description (17).

**Fig. 3.**
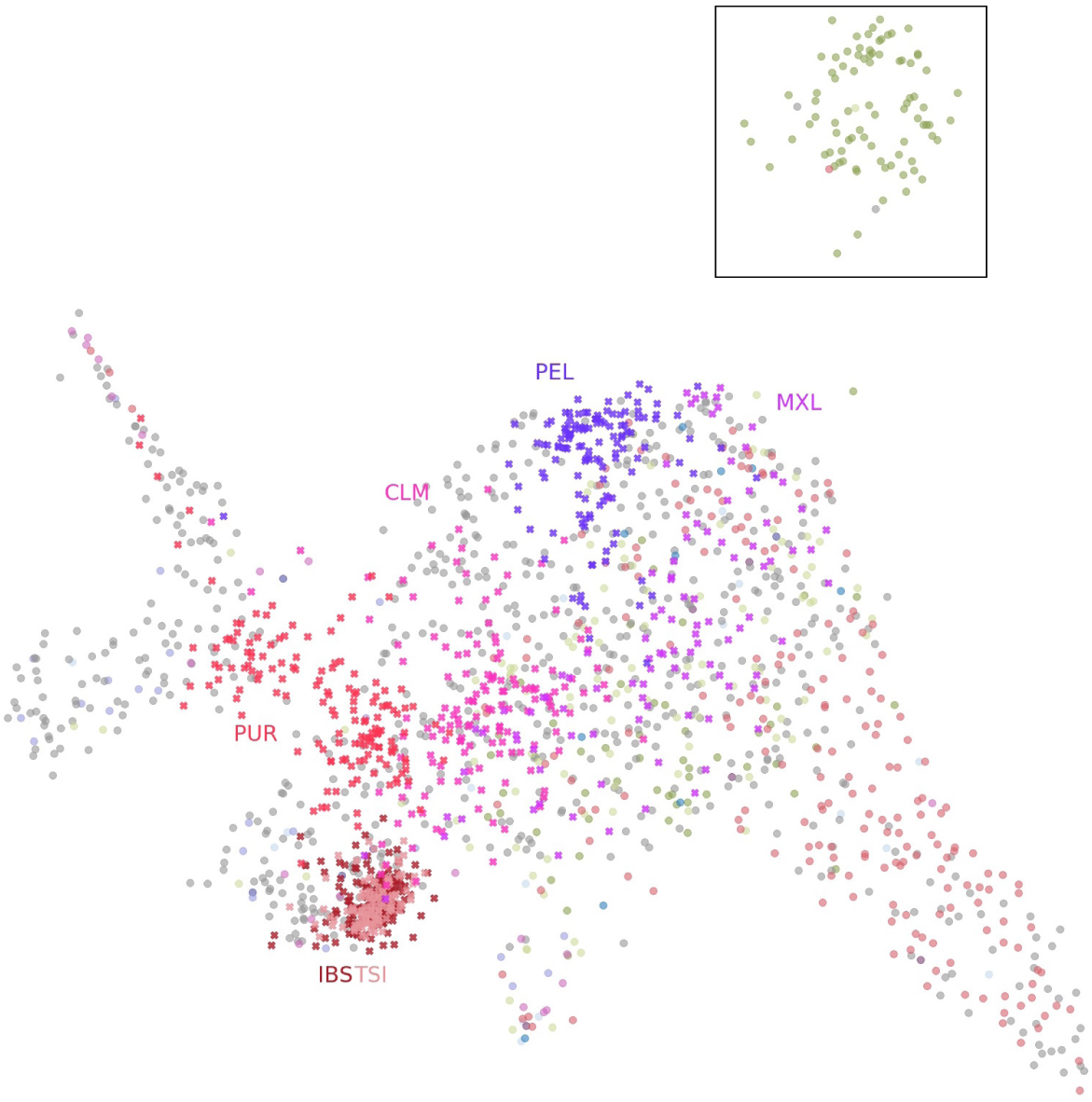
Applying UMAP to the top 7 principal components of the self-identified Hispanic population of the HRS reveals a cluster (highlighted). Coloring the points by birthplace shows they were born almost entirely in the Mountain region of the United States (New Mexico, Arizona, Colorado, Utah, Nevada, Wyoming, Idaho, and Montana). When populations from the 1KGP are projected onto the UMAP embedding they do not map to the cluster. Six 1KGP populations are presented: CLM, Colombian in Medellin, Colombia; IBS, Iberian in Spain; MXL, Mexican in Los Angeles, California; PEL, Peruvian; PUR, Puerto Rican; TSI, Tuscani in Italy. Figure S38 presents the same projection of individuals from the HRS colored by estimated admixture proportions.

### Population structure in the UKBB reflects local and global genetic variation

The UKBB provides geno-type data on 488,377 individuals along with self-identified ethnic background in a hierarchical tree-structured dictionary. Participants provided ethnic background on two occasions. We used the initial ethnicity after finding minimal differences between the two. The dataset is majority White (88.3% British, 2.6% Irish, 3.4% other), with large populations identifying as Black (1.6% either African, Caribbean, or other), Asian (1.9% either Indian, Pakistani, Bangladeshi, or other), Chinese (0.3%), an other ethnic group (0.8%), mixed ethnicity (0.6%), or an unavailable response (0.5%).

UMAP on the top 10 principal components reveals both continuous and discrete population structure (figure 4b): The patchwork of local topologies identifies continuous structure within the British population as well as admixture gradients despite the very unbalanced population sizes. The result is a comprehensive portrait of genetic variation capturing population relationships not visible using other methods, succinctly illustrating the complex structure of large and multi-ethnic datasets.

**Fig. 4.**
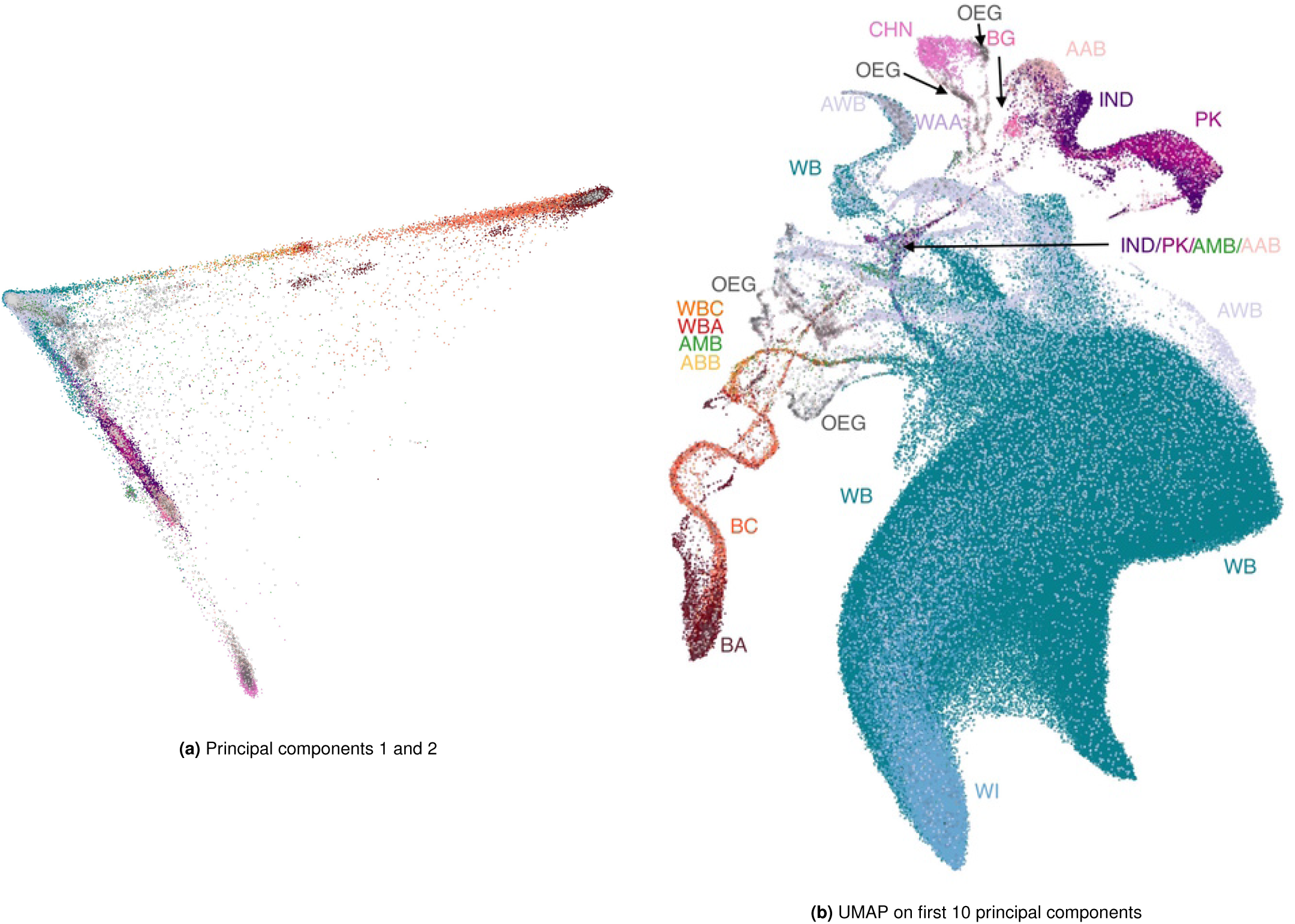
The UKBB projected onto two dimensions, colored by self-reported ethnic background. (a) The first two principal components, showing the usual triangle with vertices corresponding to African, Asian/Native American, and European ancestries, and intermediate values indicating admixture or lack of relationship to the vertex populations. (b) UMAP on the first 10 principal components. The cluster of White British and White Irish individuals is greatly expanded, with the Irish forming a distinct sub cluster mixed with the White British population. South Asian and East Asian individuals form their separate clusters, as do individuals of African or Caribbean backgrounds. Population clusters are connected by “trails” comprised of large proportions of individuals with mixed backgrounds. BA, Black African; BC, Black Caribbean; BG, Bangladeshi; CHN, Chinese; IND, Indian; PK, Pakistani; WB, White British; WI, White Irish; WBC, White and Black Caribbean; WBA, White and Black African; WAA, White and Asian; AAB, Any other Asian Background; ABB, Any other Black Background; AWB, Any other White Background; AMB, Any other Mixed Background; OEG, Other ethnic group.

The largest body in the figure consists of the White British and Irish populations. The Irish population concentrates in a portion of this group, but many individuals are also scattered throughout the British-identified population. Individuals identifying as Black African and Black Caribbean partially overlap, but admixed individuals form distinct trails leading to Asian and European clusters. Chinese individuals form a cluster, within appears to be a broader East Asian super-population; Indian, Pakistani, and Bangladeshi populations form a closely bound group as well. The East Asian and South Asian super-populations each have large clusters of individuals who identify as having an “other Asian background” or belonging to an “other ethnic group”. The patchwork of genetic neighbourhoods is connected by trails of admixed individuals. These trails come together in a nexus of individuals with a variety of ethnicities; many claim mixed ancestry, and there are clear groups of individuals who belong to an “other ethnic group”. Although their ethnicities are unknown to us, given their proximity to African, South Asian, and White individuals, possible candidates for these groups are North African, Middle Eastern, and West Asian backgrounds. Additionally, there are many individuals whose ethnicity is White but neither British nor Irish (AWB) forming clusters distinct from the British and Irish cluster.

Figure 5 presents the projection in figure 4b colored in by geographical coordinates from the Ordnance Survey National Grid (OSGB1936), with distances defined as a north or east position relative to the Isles of Scilly. UMAP coordinates within the “White British” cluster broadly map to geographic coordinates, as has been observed in Europe-wide data(4). Most admixture lines connect to the South East corner of this cluster, corresponding to the position of the city of London and reflecting its high migrant population.

**Fig. 5.**
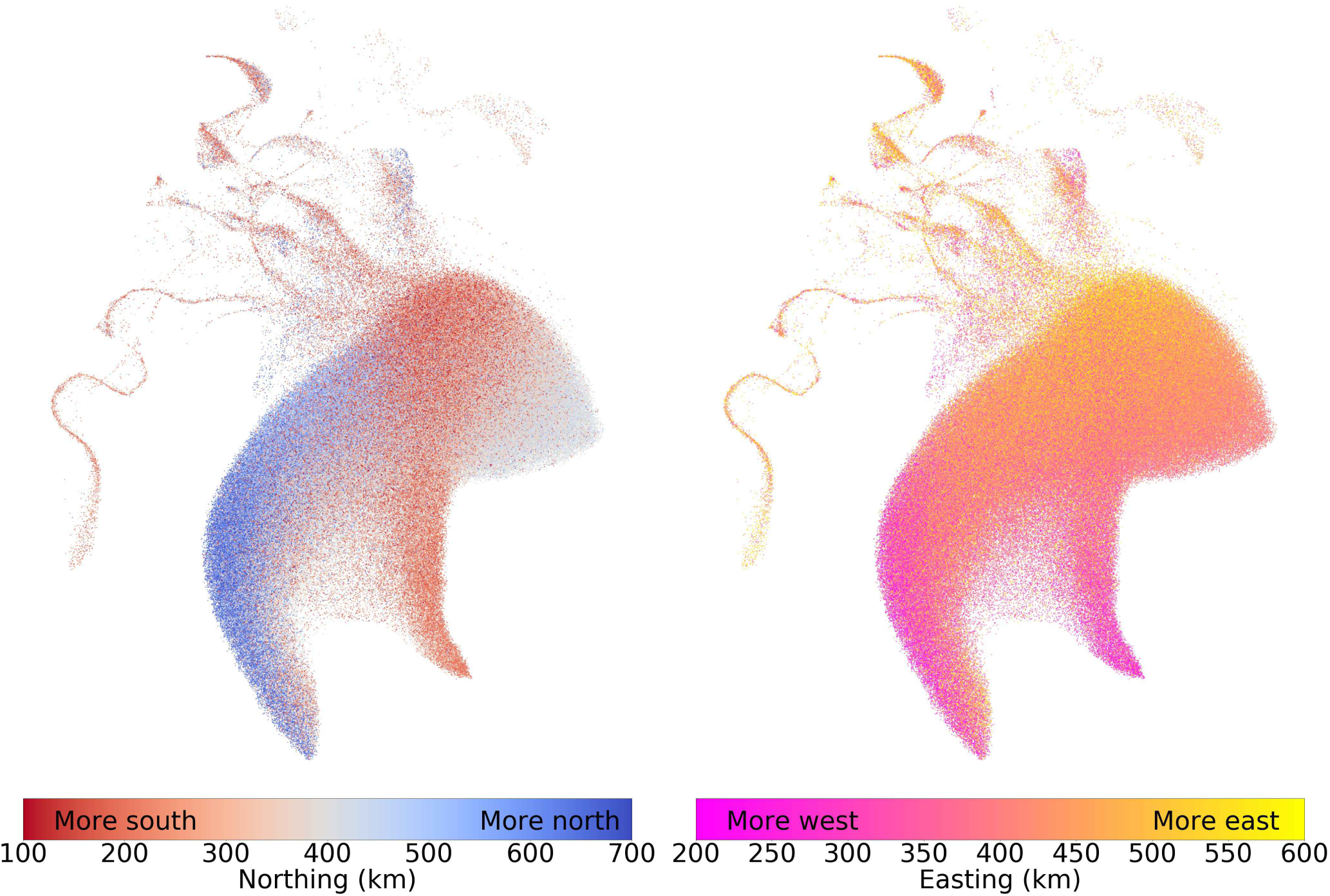
The UKBB projected onto two dimensions using PCA-UMAP with each individual colored by their geographical coordinates of residence. Coordinates follow the UKBB’s OSGB1936 geographic grid system and represent distance from the Isles of Scilly, which lie southwest of Great Britain. The left image colors individuals by their north-south (“northing”) coordinates, and the right image colors them by their east-west (“easting”) coordinates. Adding more components creates finer clusters. (figures S4 and S5). Individuals with missing geographic data are not shown. To prevent outlying individuals from washing out the color scheme, northing values were truncated between 100km and 700km, and easting values were truncated between 200km and 600km. To protect participant privacy, data has been randomized as explained in the materials and methods section.

The detailed shape of extended clusters is not stable as we vary the number of PCs included. Figure S3 shows a UMAP plot using the top 20 PCs from the UKBB. The shape of the “White British Cluster” is notably different, and we observe finer patterns of geographic variation, yet the qualitative observations made above are maintained. As an alternate visualization of diversity’s correlation with geography, we performed a 3D UMAP projection and converted the normalized UMAP values into RGB values, allowing us to plot individuals on a map of Great Britain, emphasizing both spatial gradients of genetic relatedness and increased diversity in urban centers (figure 7). The patterns in rural areas observed are similar to those reported in (18) using the haplotype-based CHROMOPAINTER on British individuals whose grandparents lived nearby.

**Fig. 6.**
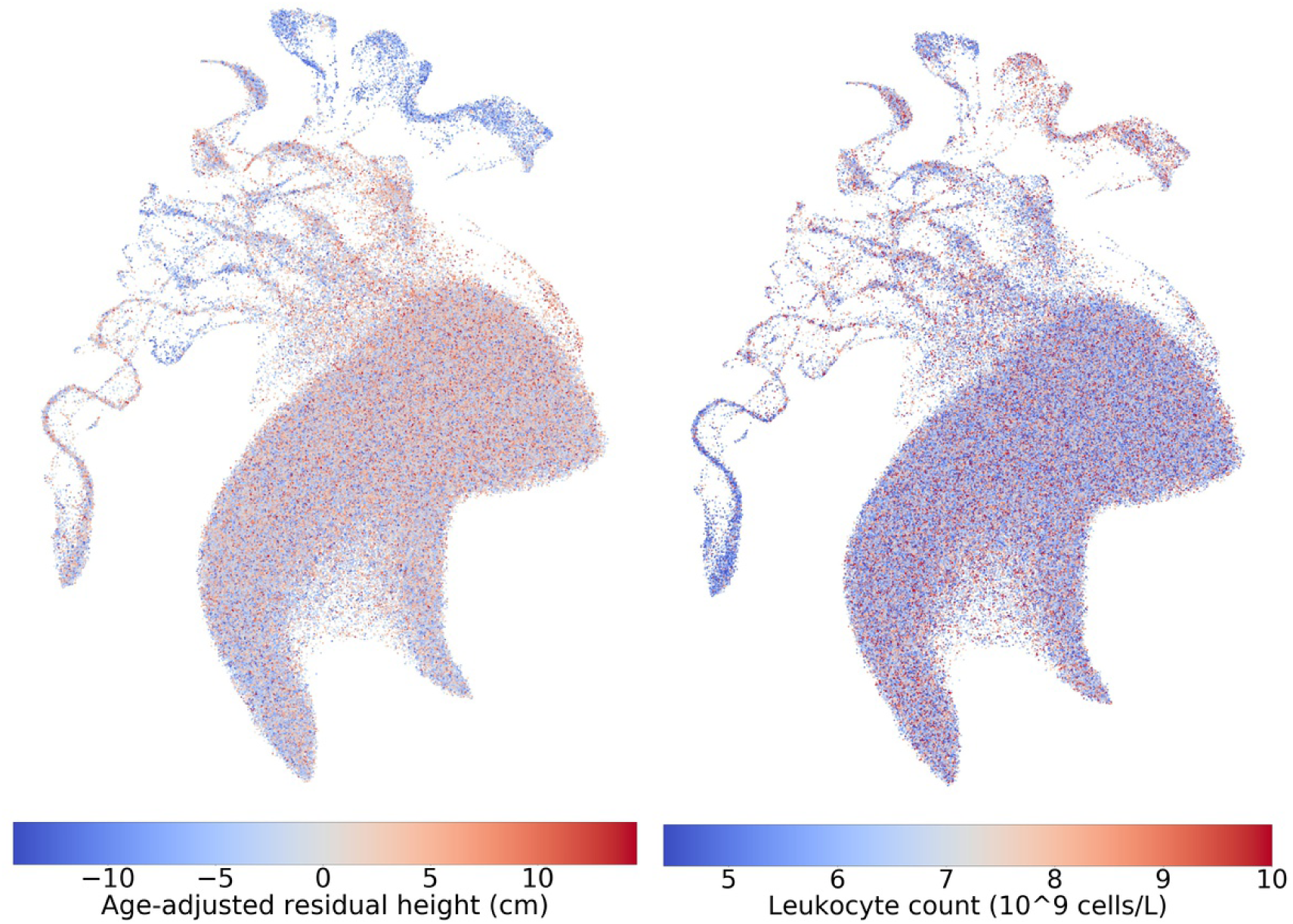
The UKBB projected onto two dimensions using PCA-UMAP (as in 4b), with females colored by age-adjusted difference from mean population height (left) and leukocyte counts (right). To protect participant privacy, data has been randomized as explained in the materials and methods section.

**Fig. 7.**
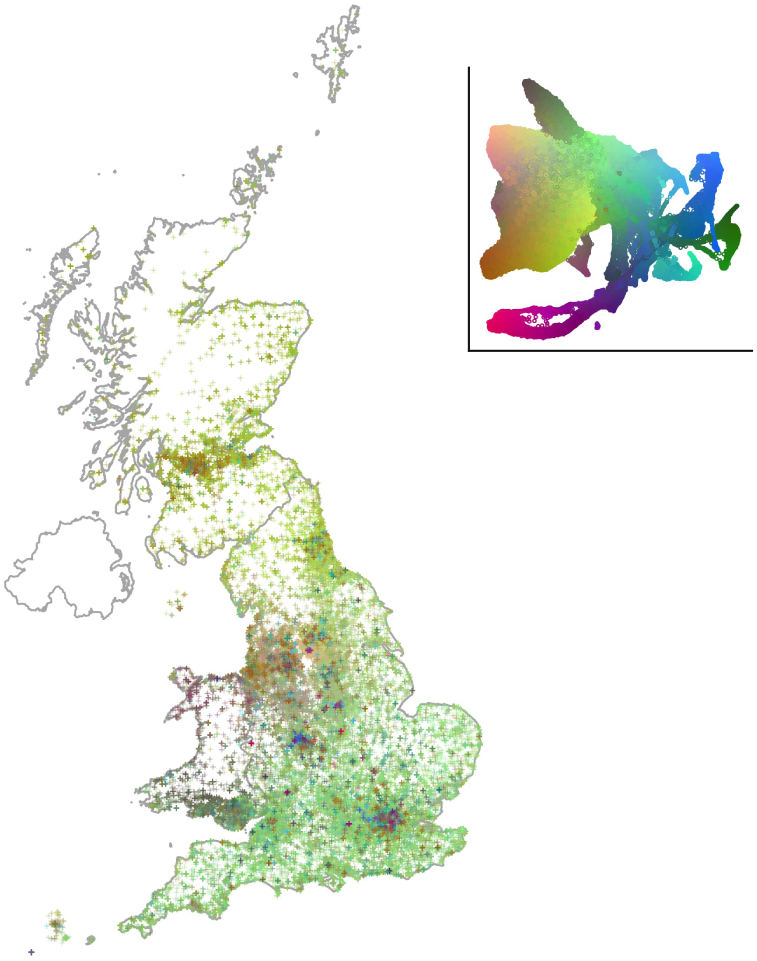
A map of Great Britain colored by a 3D UMAP projection. Each individual is assigned a 3D RGB vector based on 3D UMAP coordinates (inset): Individuals who are closer to each other in the projection will appear to be closer in color. Patterns in genetic similarity are visible in Scotland, South England, the East and West Midlands, and major urban centres. A flattened 2D view of of the 3D projection used for coloring is presented in the top right. To protect participant privacy, data has been randomized as explained in the materials and methods.

Similarly to UMAP, t-SNE applied to the UKBB data both displays diversity within the “White British” population and identifies clusters among other groups. However, it has three drawbacks: it is much slower, requiring 2.26 hours for its first thousand iterations alone on 10 principal components against UMAP’s 14 minutes; it fails to find a global optimum, which results in a scattering of individuals and groups that are not stable across independent runs; and it does not identify continuity between different continental groups resulting from admixture (figure S40).

### Patterns in phenotype related to population structure

Our involvement with the UK biobank data is through a project on autoimmune disease and asthma. More than in geographic coordinates, we are interested in whether genetic population structure correlates with phenotypes and covariates of interest.

Covariates such as height (figure 6) and autoimmune and asthma-related measures (figures S7 to S18) correlate strongly with both discrete and continuous population structure. Several populations in figure 6, including South Asian, East Asian, African, and several unidentified ethnic groups have noticeably lower-than-average heights. More subtle patterns are also visible: the area of the projection in figures 4b with the cluster of White Irish people appears more blue than the main body of White British individuals; an unpaired two sample t-test of self-identified White Irish and White British individuals reveals statistically significant differences in age-adjusted mean height between the populations, with British males being taller on average by 0.846cm (p-value 2.10 × 10^-23^) and British females by 0.763cm (p-value 3.65 × 10^-23^) (see figures S25 and S26 for boxplots). Height differences between Irish and British populations have been previously observed but the direction of the difference is not consistent(19, 20).

Forced expiratory volume in 1 second (FEV1) (figures S11, S12) also shows strong correlations with certain populations — South Asian, African, and Caribbean — having considerably lower measurements on average (see figures S23 and S24 for boxplots and p-values). Notably, there appears to be a juncture in the admixture continuum, highlighted in figure S31, where the distribution of FEV1 changes. This roughly corresponds to the transition from Black African/Caribbean individuals to those who identified having mixed backgrounds. Boxplots and statistical testing suggest that relative to White British populations FEV1 values are significantly lower for Black African and Black Caribbean populations, but not for White and Black Caribbean and White and Black African populations. Unidentified populations highlighted in figure S32 suggest that one ethnic group close to the Chinese may have higher than average FEV1 values compared to the relatively low values of the Chinese themselves; while another close to European and British populations has lower values relative to the population mean. These results merit further investigation and underscore the exploratory value of PCA-UMAP — these populations are largely unidentified and there is no straightforward way to separate them within the data otherwise.

### Comparing t-SNE and UMAP

Assessing the relative merits of data visualization methods is notoriously difficult, because different users might give different importance to different features of the data. Methods differ in their choice of a similarity metric between high- and low-dimensional representation, which are hard to assess, but also in their strategy to optimize the metrics, which are amenable to direct comparison. This is especially true with UMAP and t-SNE which, despite different mathematical motivations, have computationally similar similarity matrices. Given that t-SNE generated a smattering of clusters for the minority populations when applied to the UKBB, we hypothesized that this was due to a convergence issue rather than a property of the global optimum of the t-SNE metric.

To compare UMAP to t-SNE directly and assess the role of convergence, we ran UMAP and Python’s sklearn implementation of t-SNE, and a hybrid approach where the early exaggeration phase of t-SNE (typically the first 250 iterations) has been replaced by a full UMAP run (600 epochs for the 1KGP data and 230 epochs for the HRS and UKBB data). The three approaches produce similar results in smaller datasets such as the 1KGP and HRS, with the main differences being in run-time and some qualitative factors such as clusters being more interconnected when using UMAP (figures S41, S42). When applied to the UKbiobank, we find (perhaps subjectively) that UMAP provides a more compelling visualization of the data compared to both version of t-SNE. We also find that the hybrid approach produces a better value for the t-SNE metric than the default t-SNE approach (figure S46), confirming that the default t-SNE optimization fails to identify the global t-SNE minimum. Qualitatively, the standard t-SNE initialization generated a large number of arbitrary clusters, while initializing t-SNE with UMAP preserved much of the large scale structure created by the UMAP embedding but broke the global continuity between clusters (figure S43). An animation of t-SNE iterations starting from the UMAP embedding can be seen in the supplementary files. Given the many desirable properties of UMAP, such as computational performance and intuitive placement of individuals within a global genetic continuum, we conclude that UMAP is a superior alternative to t-SNE for large and diverse genomic datasets.

## Discussion

Understanding population structure is important to identify confounders in medical genomics and studies of anthropology and human evolution. PCA of genomic data reflects genealogical and geographic data, but visualization in large datasets still requires scanning through a large number of pairwise plots. UMAP condenses these components and comprehensively illustrates information — phenotypic, geographic, and ancestral — contained within genotypes on fine-scale levels and within the context of a global population structure. In large datasets where the number of significant PCs is large, the resulting representation has important advantages over PCA alone and provides a superior visualization to t-SNE.

Examinations of clustering in the three datasets provided many intriguing clusters that would otherwise have been difficult to identify. In particular, several areas from figure 4b, highlighted in figure S6, show multiple unidentified groups related to each of the East Asian and South Asian super-populations, as well as to either or both of African or admixed populations. Additionally, the Hispanic population of the HRS contains a geographically-restricted cluster that could not have been identified from pairwise examinations of principal components. The 1KGP — frequently used in medical and population studies — contained splits in the Gujarati and Punjabi population samples that were not visible PCA or Admixture analysis alone (although a split among Gujarati is arguably visible in the Admixture analysis with K=12 in (7)).

Application to the UKBB underscores the strength of PCA-UMAP in large cohorts. We see clear, fine relationships between genotype and phenotype and geography, and this is presented in a visualization that accounts for natural genetic clustering. Figures 6, S11, and S6 demonstrate phenotypic variation within and across clusters, with phenotypes such as height showing continuous variation across admixture edges, as expected from genetically controlled traits, and others, such as leukocyte counts or FEV1, showing sharper boundaries, as expected from environmentally determined traits.

Importantly, using UMAP is straightforward and fast. Most of the plots presented in this article were generated directly from the PCA data using UMAP with default parameters, except that we set the “minimum distance” parameter to 0.5 which made fine features on UMAP more visible (results with default parameter 0.1 provided qualitatively similar results). Given PCA data and a desktop computer, UMAP can be performed in 15 to 25 minutes on a sample of hundreds of thousands of individuals over tens of dimensions.

There are downsides to using nonlinear approaches to visualize the data. Both UMAP and t-SNE are sensitive to sample size, and spend more visual real estate for populations with larger sample sizes compared to PCA. This is useful to identify significant patterns in a cohort, but it makes comparing visualization across cohorts difficult. Nonlinearity also complicates the interpretation of results. Distances in UMAP or tSNE space should not be used as a proxy for genetic distance. We did not assign meaning to wiggles in UMAP figures, which occurred consistently in the UKBB but may be an artifact of the dimensional reduction strategy rather than a meaningful feature of the data. Hand-waving interpretations of pretty plots has a history of getting population geneticists in trouble (as pointed out, e.g., in (21)): visualization is not a replacement for statistical testing.

## Conclusion

With these caveats in mind, a priori data visualization plays a central role in quality control, hypothesis generation, and confounder identification for a wide range of genomic applications. Nonlinear approaches, despite their limitations, become increasingly useful as the size of datasets increases: We have shown that UMAP, in particular, reveals a wide range of features that would not be apparent using linear maps. Given its ease of use, breadth of results, and low computational cost, we propose that UMAP should become a default companion to PCA in large genomic cohorts.

## Materials and Methods

We used genotype data from 12,454 individuals from the Health and Retirement Study (HRS), genotyped on the Illumina Human Omni 2.5M platform(13). Principal components were computed in PLINK v1.90b5.2 64-bit(22) using variants with a minor allele frequency greater than 0.05, Hardy-Weinberg p-value of more than 1 × 10^-6^, and genotype missing rate of less than 0.1, and sample with genotype missing rate of less than 0.1. We used the principal components of genotype data from 488,377 individuals in the UK BioBank (UKBB) as computed by the cohort (14). We used genotype data from 3,450 individuals from the 1KGP project using Affy 6.0 genotyping(7).

Scripts for all tests and plotting functions can be found on https://github.com/diazale/gt-dimred. A demo version using freely available 1KGP data is available at https://github.com/diazale/1KGP_dimred. PCA and standard t-SNE were done with Scikit-learn(23). UMAP was performed using a Python implementation(11). Statistical testing was done in SciPy(24) and StatsModels(25).

Both UMAP and t-SNE feature a number of adjustable parameters. Among the parameters that we varied, the number of PCs used in pre-processing of the data has the largest effect for both methods (see figures S1 and S2).

We tested different choices for perplexity in t-SNE. The default value of 30 provided comparable performance to other parameter choices. Similarly, we tested different parameter choices for UMAP, with the clearest results generated by specifying 15 nearest neighbours (the default value) and a “minimum distance” between points in low dimensions of 0.5. UMAP developers described “sensible” values for nearest neighbours as between 5 and 50 and minimum distance between 0.5 and 0.001.

UMAP and t-SNE projections were carried out on an iMac with a 3.5GhZ Intel Core i7 processor, 32 GB 1600 MHz DDR3 of RAM, and an NVIDIA GeForce GTX 775M 2048 MB graphics card.

To reduce the potential risks for re-identification from results in this publication, data has been randomly permuted so that the population characteristics are preserved but individual-level data is not presented directly in the figures. We rounded each attribute to an attribute-specific number of bins, and then permuted the data in the following way: For each point (i.e. each individual) in UMAP visualizations, and each attribute, we identified the 9 nearest neighbouring points, and copied the attribute from a randomly selected neighbor (thus allowing for the possibility of one value being printed twice). Because this process is done independently for each visualization, a given point shown on the figure will copy values from different randomly selected individuals. Additionally, spatial coordinates have random noise added (normally distributed about 0 with a standard deviation of 50km) before binning to the nearest 50km.

For each point in figure 7 we identified the nearest 50 neighbouring individuals and copied the color value from a randomly selected neighbour.

## Declarations

### Ethics approval and consent to participate

HRS data was under IRB Study No. A11-E91-13B - The apportionment of genetic diversity within the United States. UKBB data was accessed under accession number 6728.

### Consent for publication

Not applicable.

### Availability of data and material

All data is publicly available to researchers.

### Authors’ contributions

A.D.P. and S.G. designed the research. A.D.P. carried out dimension reduction and analysis. S.G. provided analysis. A.D.P and S.G. wrote the paper. L.A.T. provided the UK map visualization and analysis.

## Acknowledgements

We thank all participants in the HRS, UKBB, and 1KGP for providing their genetic data as well as the teams who generated and assembled the dataset. We also thank Chief Ben-Eghan, Jose Sergio Hleap, Mark Lathrop, Dominic Nelson, Markus Munter, Stephen Sawcer, and Audrey Grant for useful discussions about science, programming, and data access, and Selin Jessa for introducing us to UMAP. This research has been conducted using the UK Biobank Resource under Application Number 6728.

## Supporting Information (SI)

### SI Figures

**Fig. S1.**
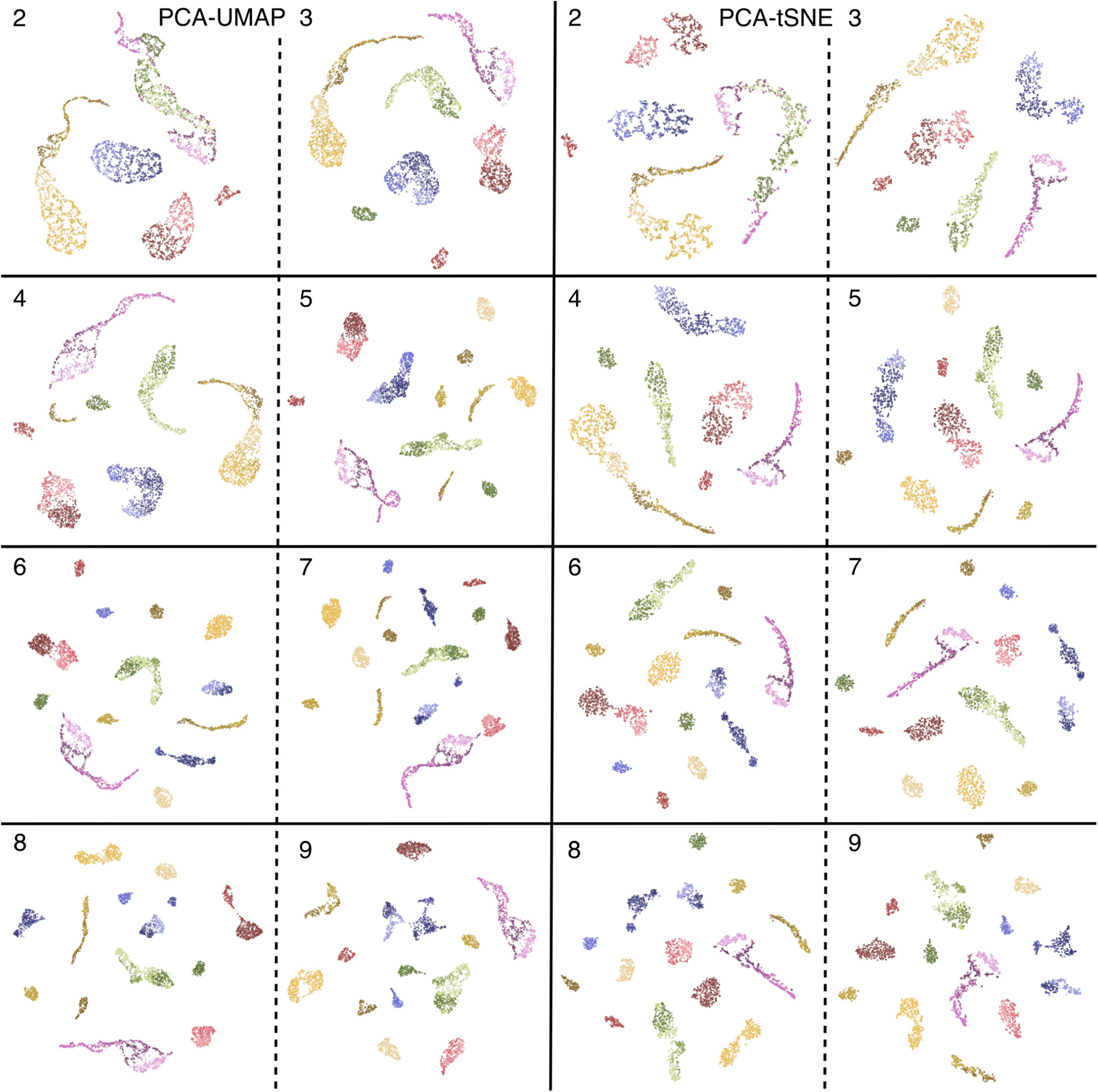
UMAP (left two columns) and t-SNE (right two columns) applied to the top principal components of the 1KGP labelled by the number of components used. Adding more components results in progressively finer population clusters using both methods.

**Fig. S2.**
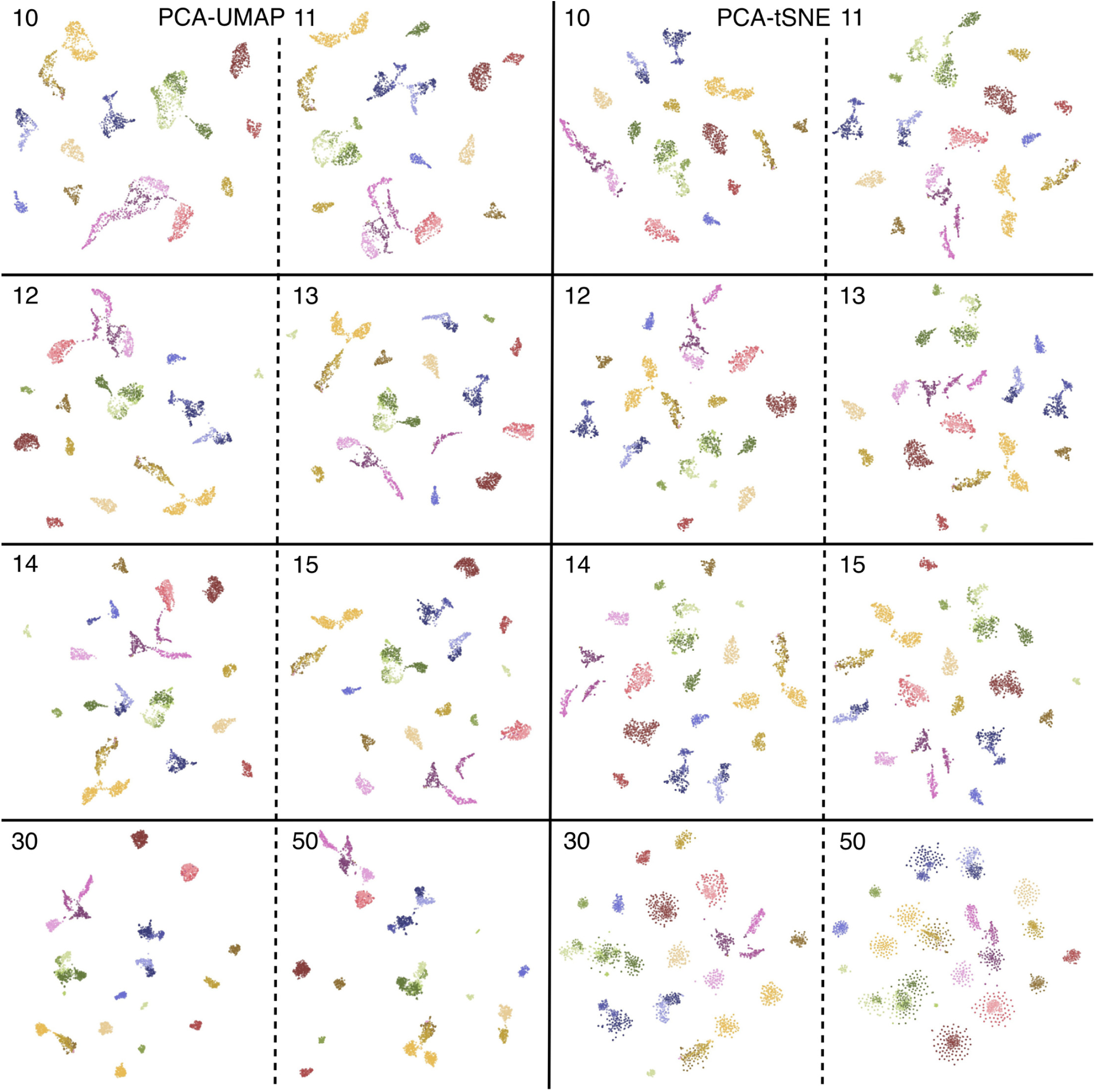
UMAP (left two columns) and t-SNE (right two columns) applied to the top principal components of the 1KGP labelled by the number of components used. Results are similar until approximately 11 components, where t-SNE breaks apart clusters of South Asian (in green) and Central and South American populations (in pink) while UMAP preserves them. At approximately 30 components populations begin to drift together with UMAP and disperse with t-SNE.

**Fig. S3.**
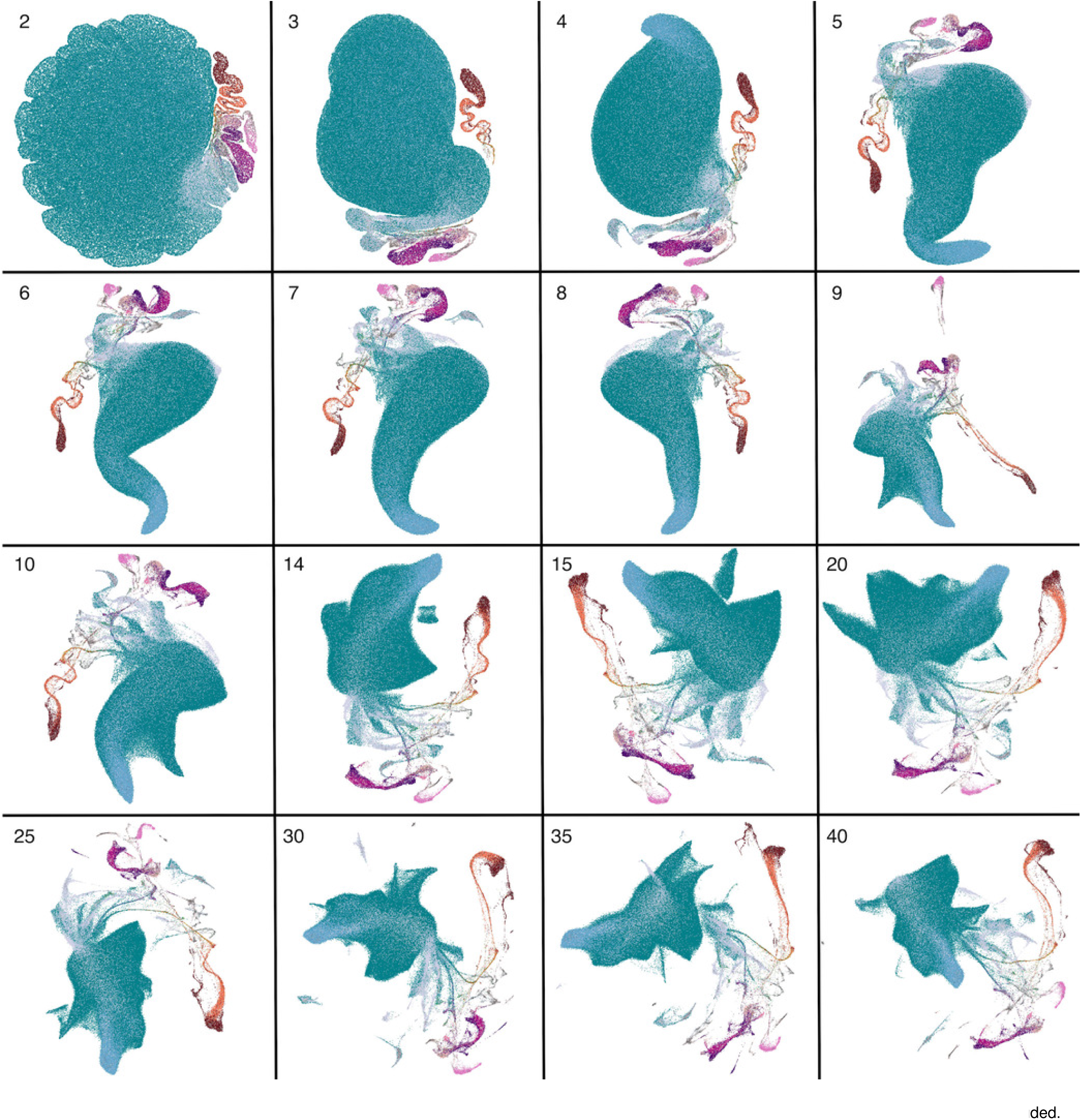
PCA-UMAP on UKBB data, colored by self-identified ethnic background. Images are labelled by the number of components included.

**Fig. S4.**
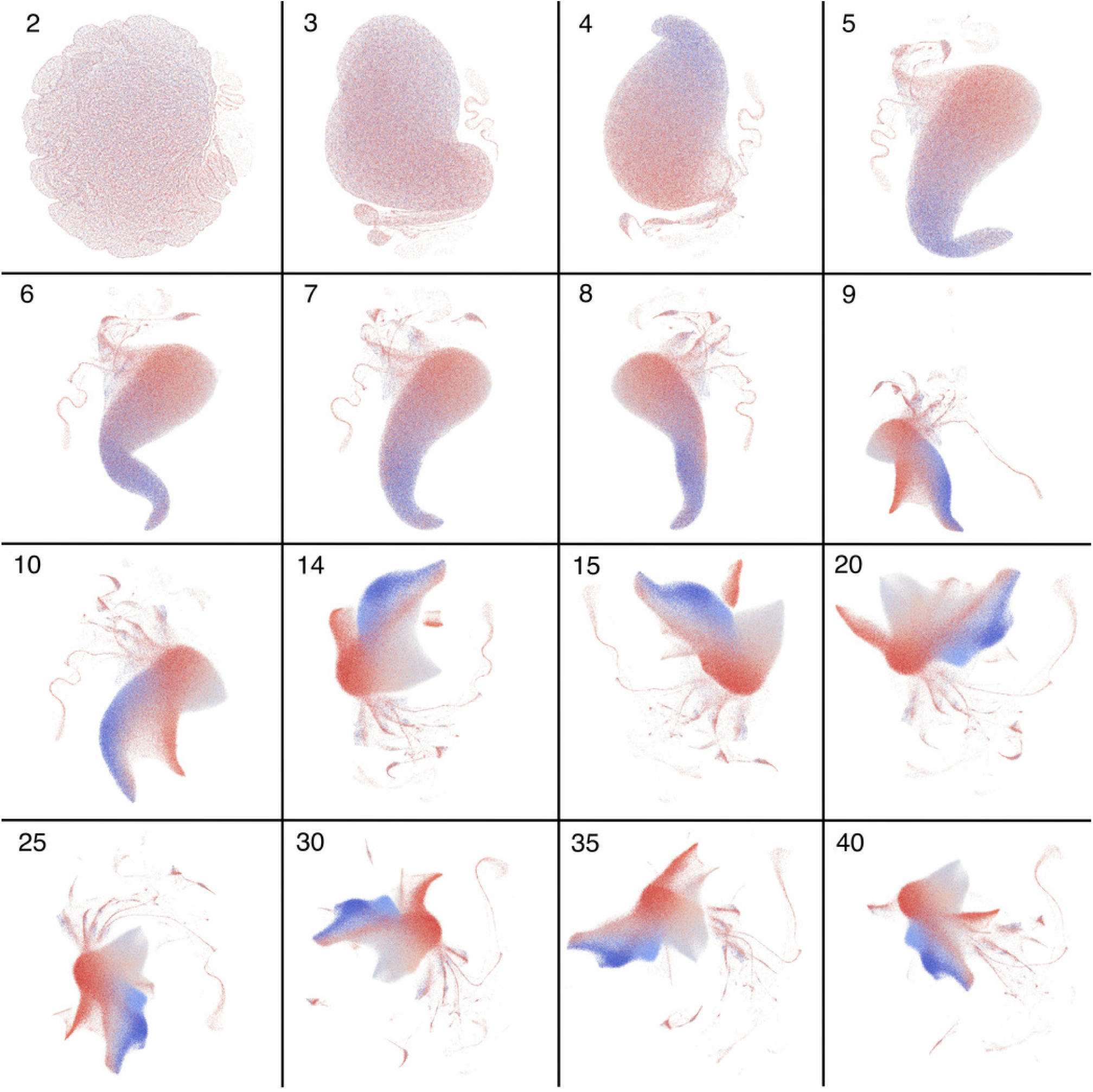
PCA-UMAP on UKBB data, colored by northing values, with more blue representing more northern coordinates and more red representing more southern coordinates. Images are labelled by the number of components included. Data has been randomized as explained in the materials and methods section.

**Fig. S5.**
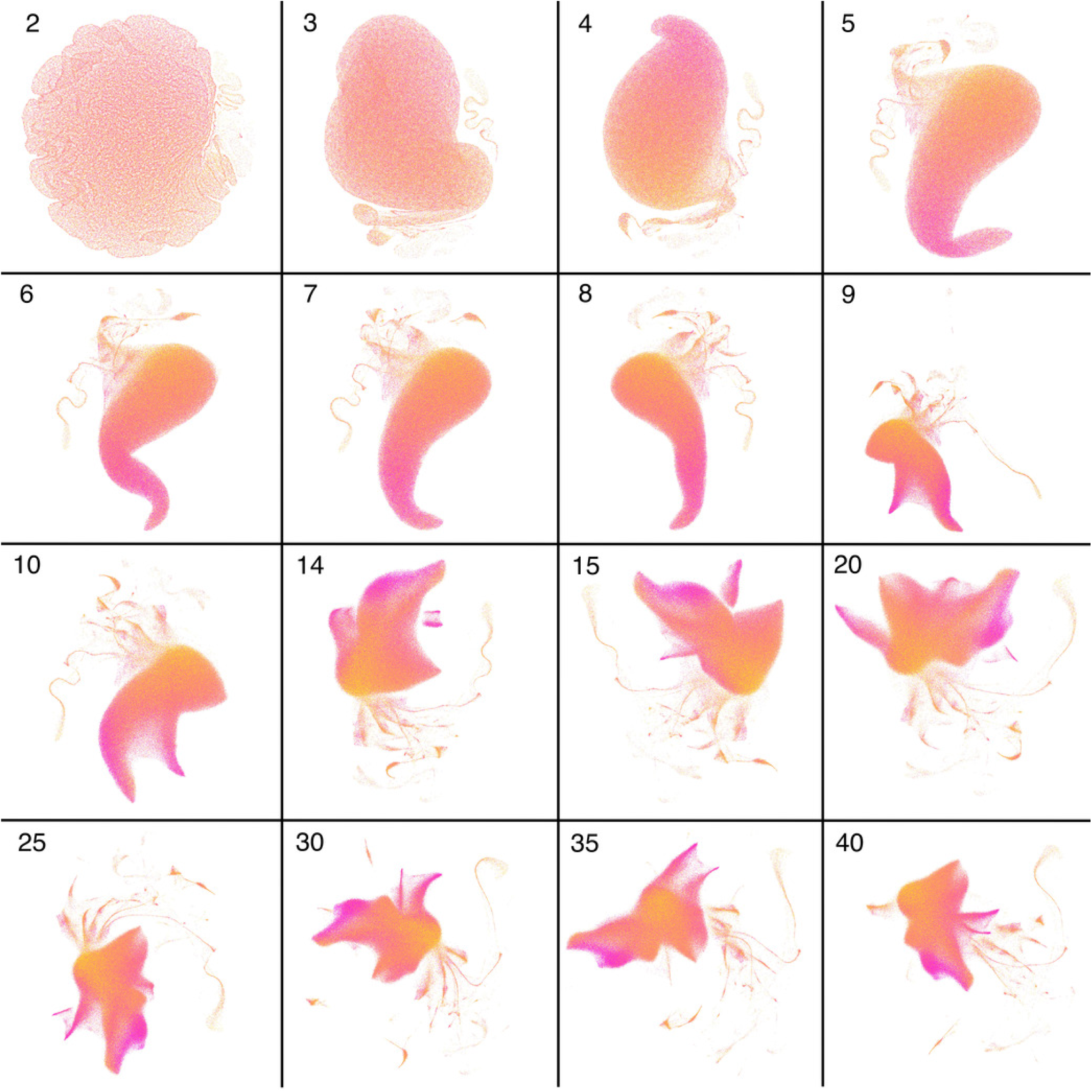
PCA-UMAP on UKBB data, colored by easting values, with more yellow representing more eastern coordinates and more pink representing more western coordinates. Images are labelled by the number of components included. Data has been randomized as explained in the materials and methods section.

**Fig. S6.**
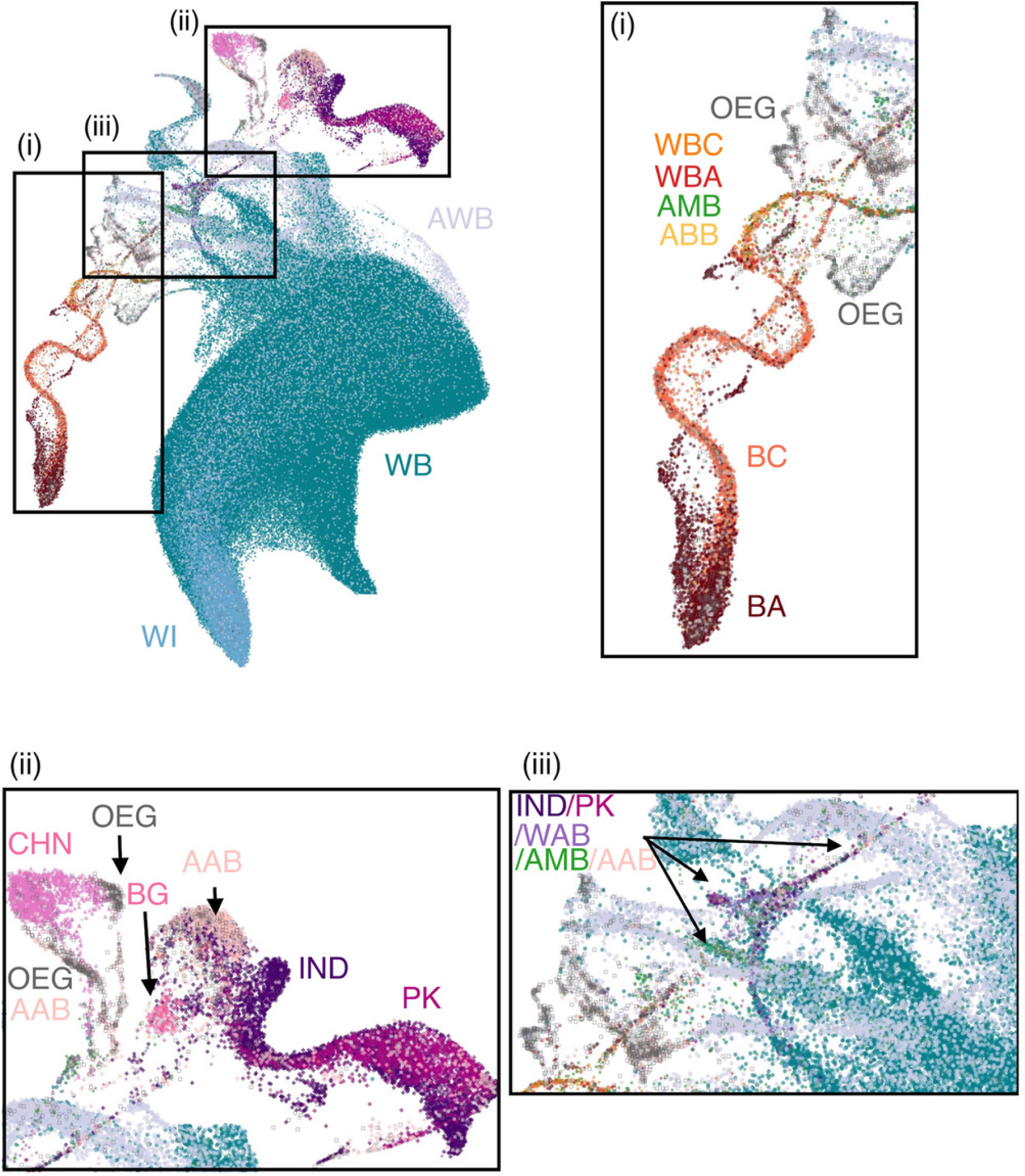
Zoomed in areas of figure 4b. Sections (i) and (ii) respectively focus on the African and Asian superpopulations, and section (iii) focuses on an area with individuals from many ethnic backgrounds. Noticeable clusters of unidentified ethnic backgrounds appear and are labelled “OEG” “(Other Ethnic Group)”.

**Fig. S7.**
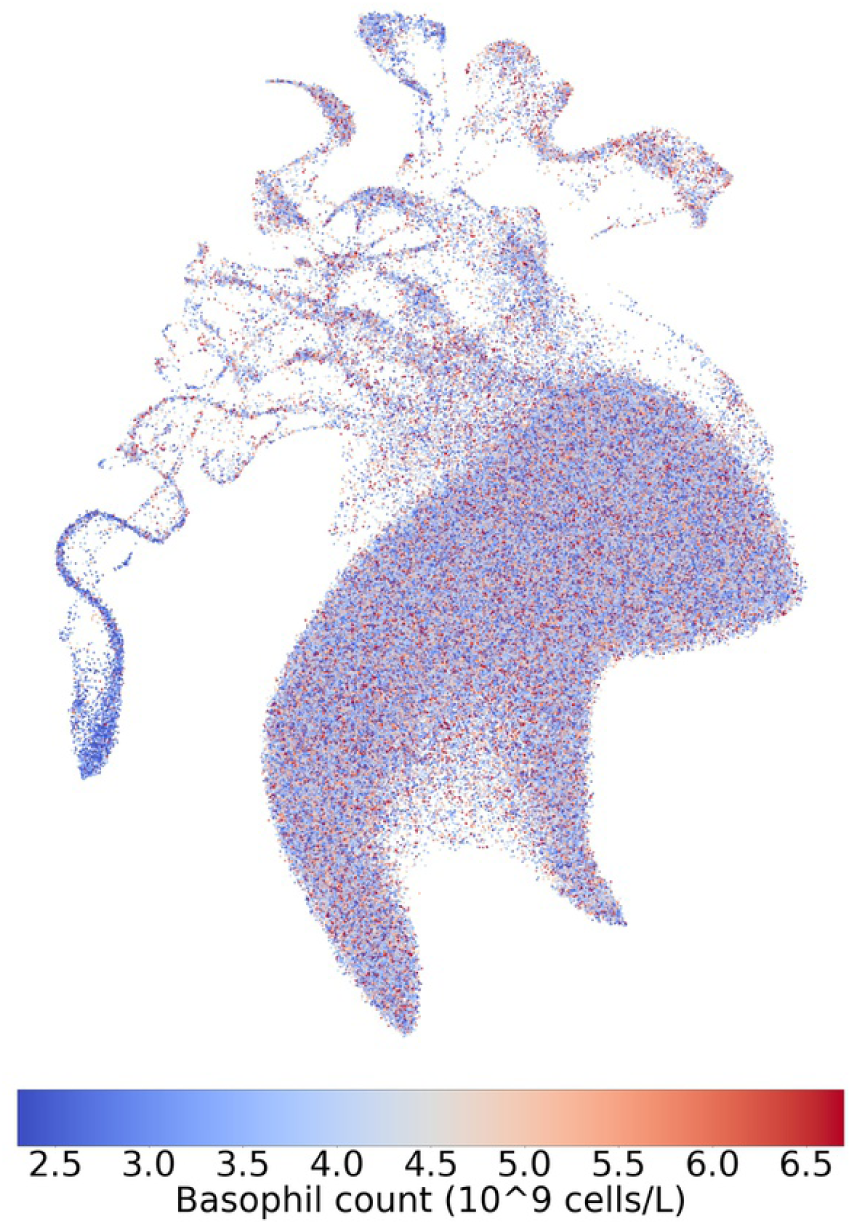
PCA-UMAP on the top 10 principal components of the UKBB colored by basophil count (female). Data has been randomized as explained in the materials and methods section.

**Fig. S8.**
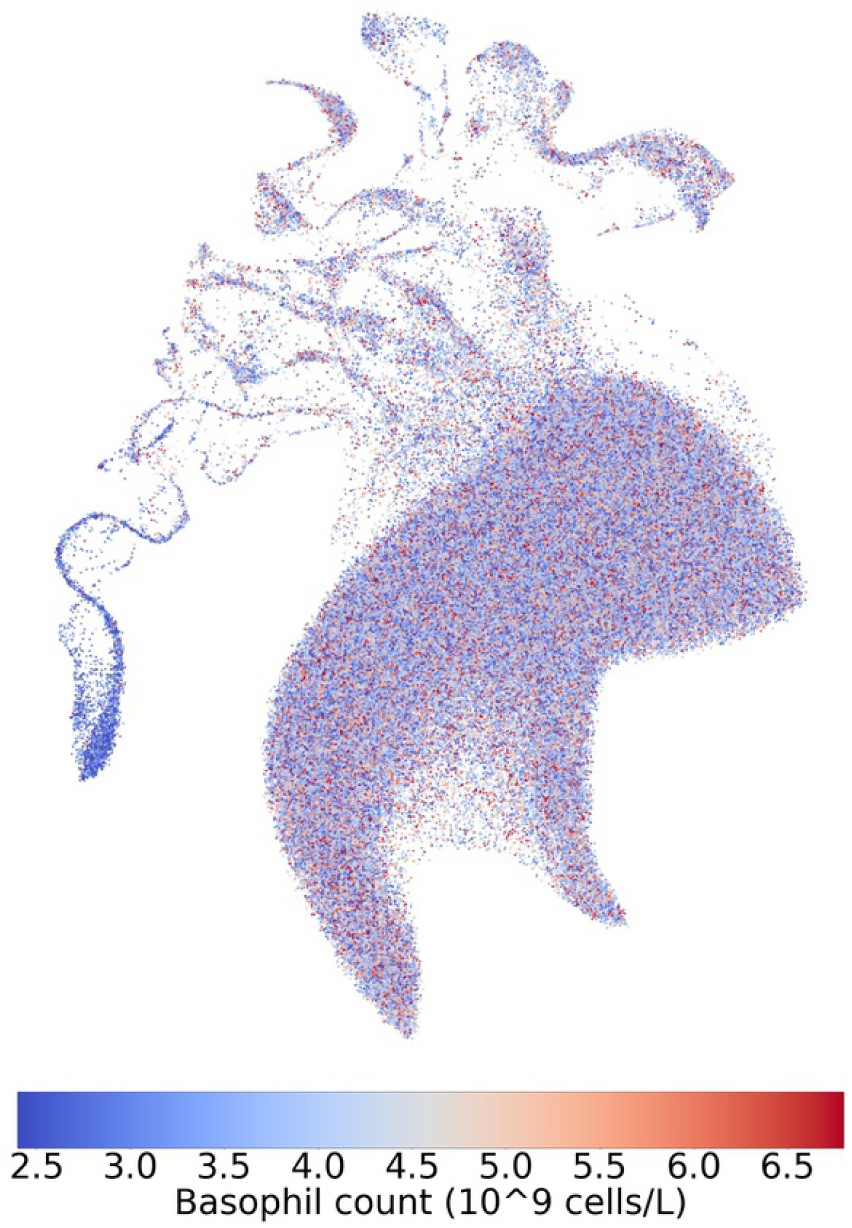
PCA-UMAP on the top 10 principal components of the UKBB colored by basophil count (male). Data has been randomized as explained in the materials and methods section.

**Fig. S9.**
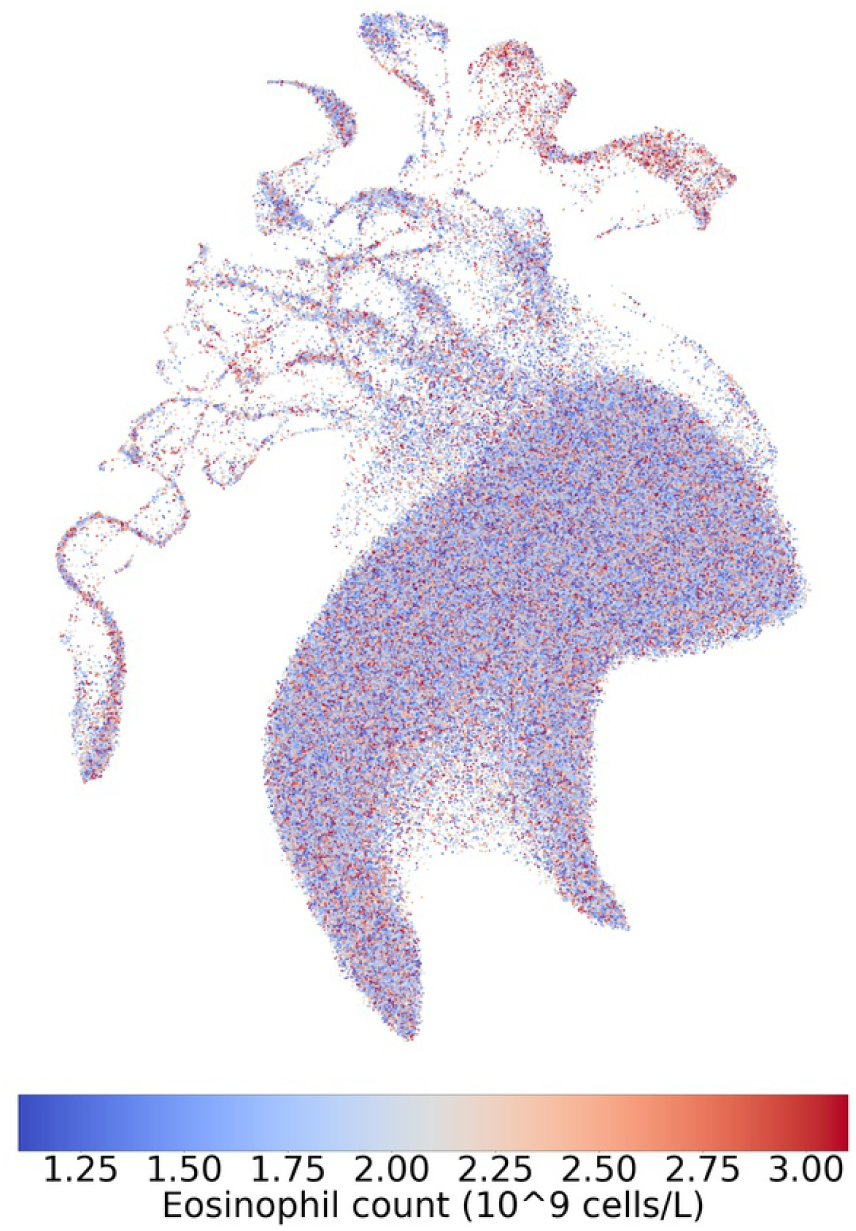
PCA-UMAP on the top 10 principal components of the UKBB colored by eosinophil count (female). Data has been randomized as explained in the materials and methods section.

**Fig. S10.**
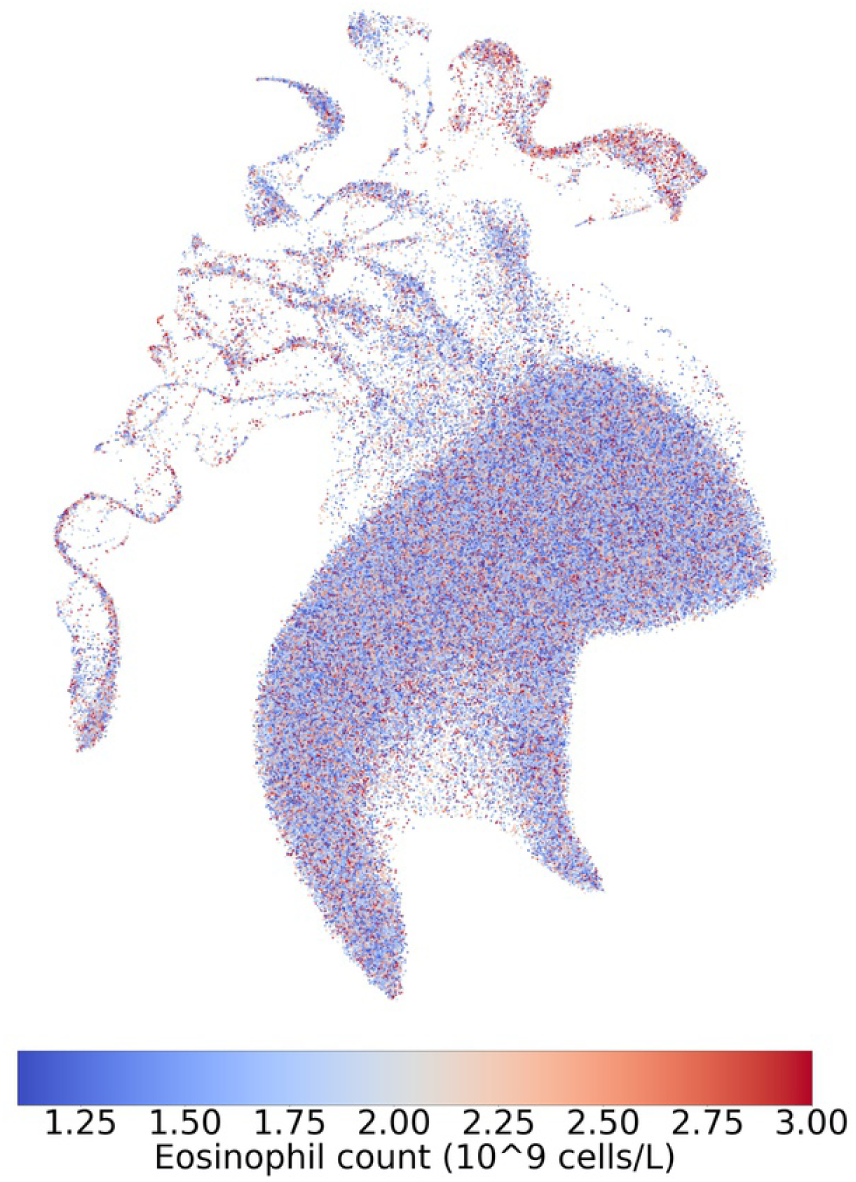
PCA-UMAP on the top 10 principal components of the UKBB colored by eosinophil count (male). Data has been randomized as explained in the materials and methods section.

**Fig. S11.**
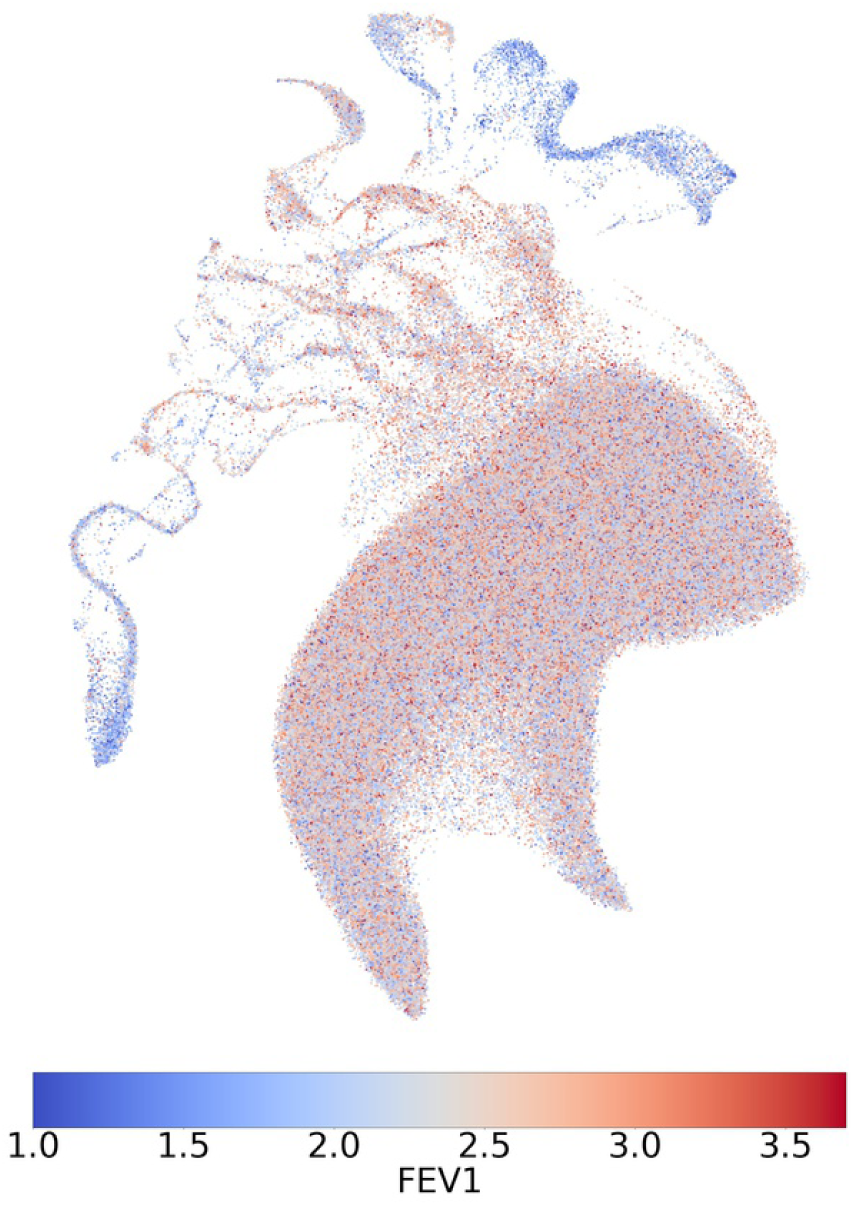
PCA-UMAP on the top 10 principal components of the UKBB colored by FEV1 (female). Data has been randomized as explained in the materials and methods section.

**Fig. S12.**
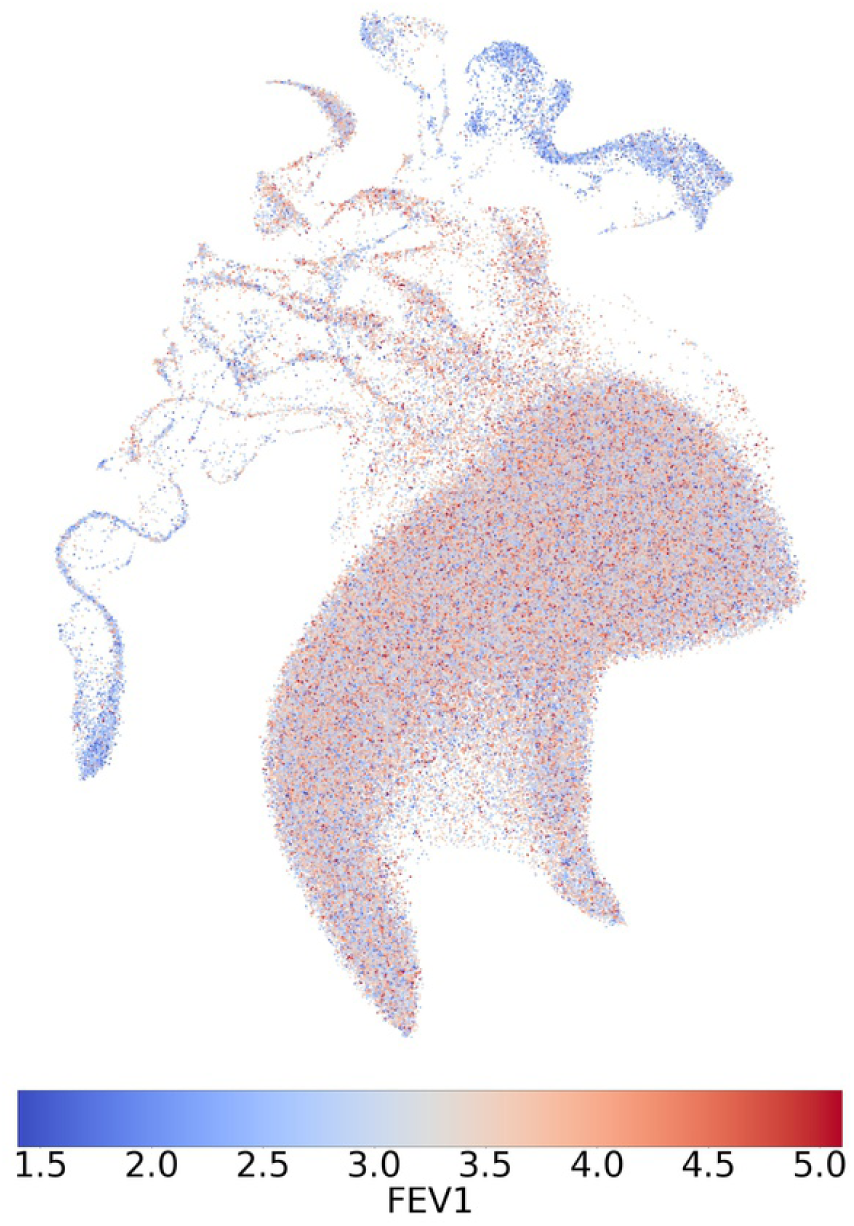
PCA-UMAP on the top 10 principal components of the UKBB colored by FEV1 (male). Data has been randomized as explained in the materials and methods section.

**Fig. S13.**
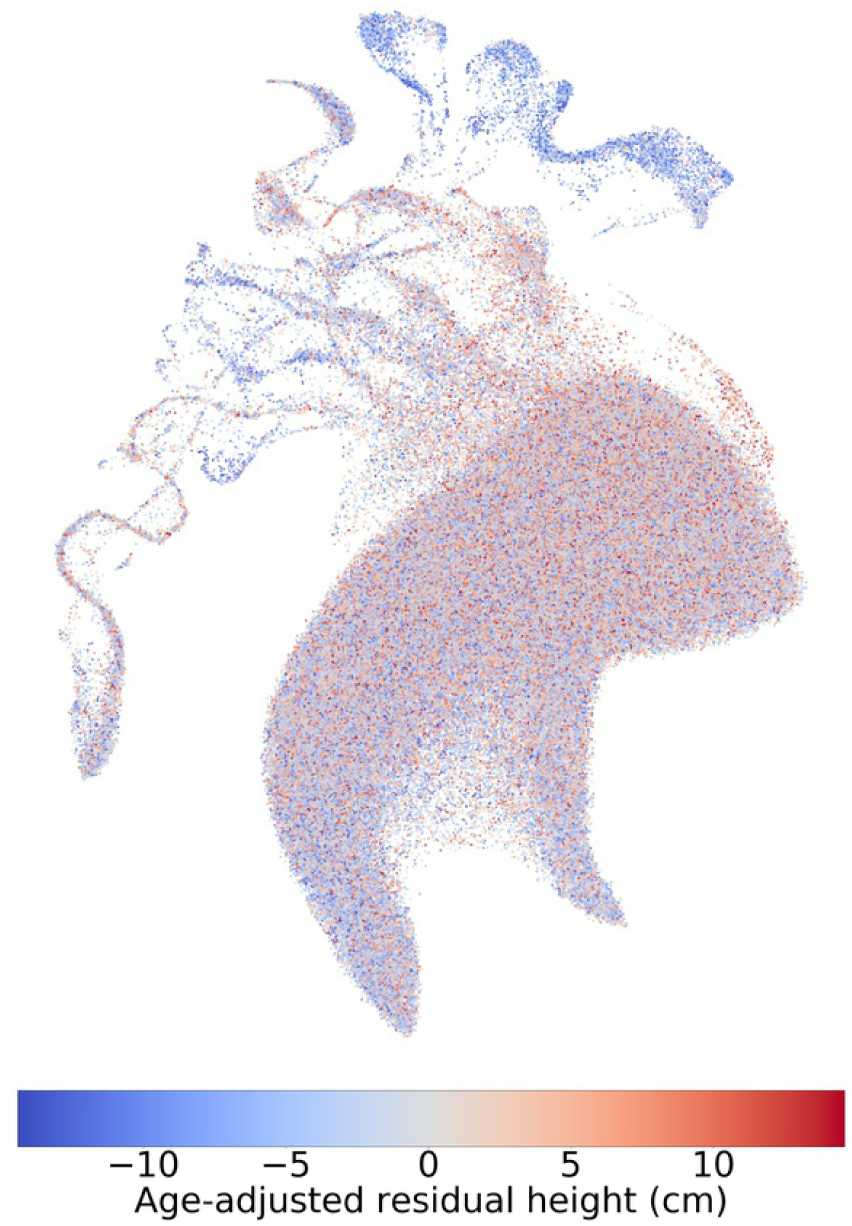
PCA-UMAP on the top 10 principal components of the UKBB colored by height (female). Data has been randomized as explained in the materials and methods section.

**Fig. S14.**
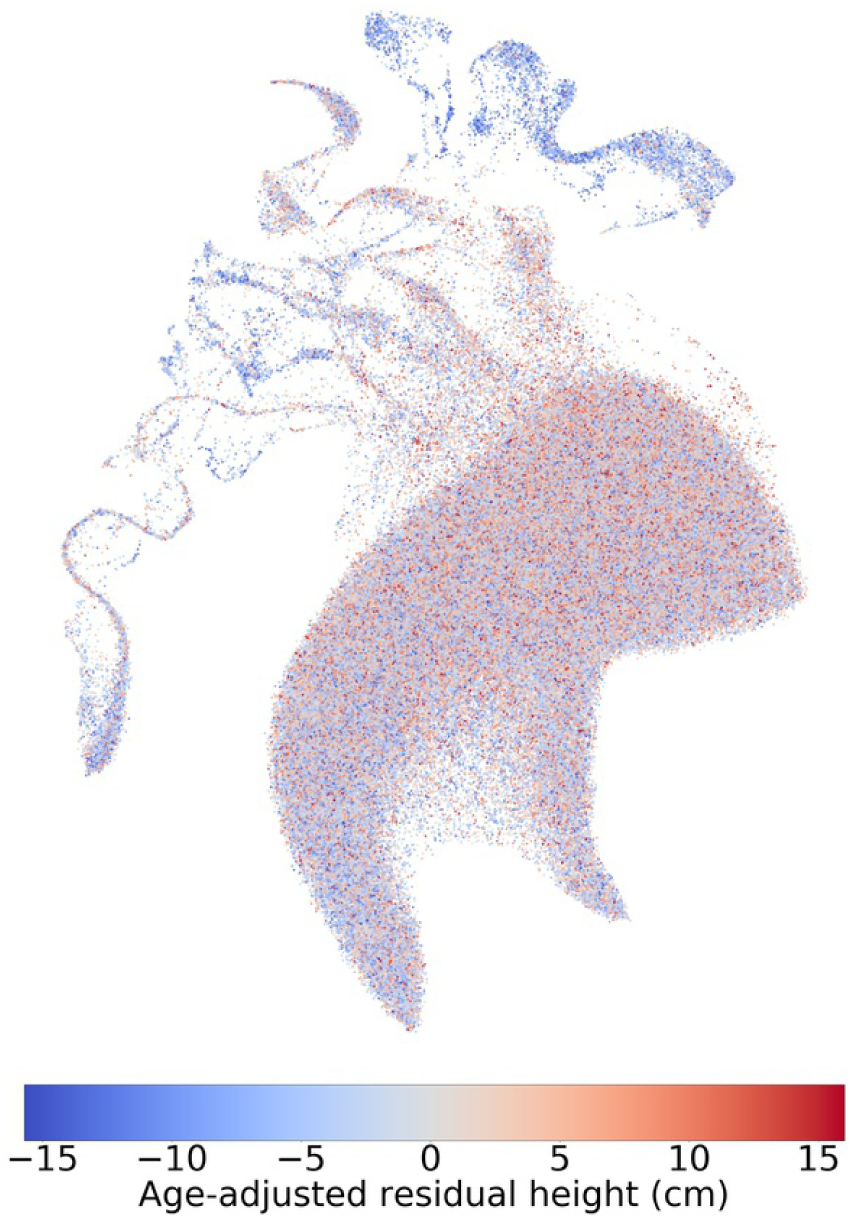
PCA-UMAP on the top 10 principal components of the UKBB colored by height (male). Data has been randomized as explained in the materials and methods section.

**Fig. S15.**
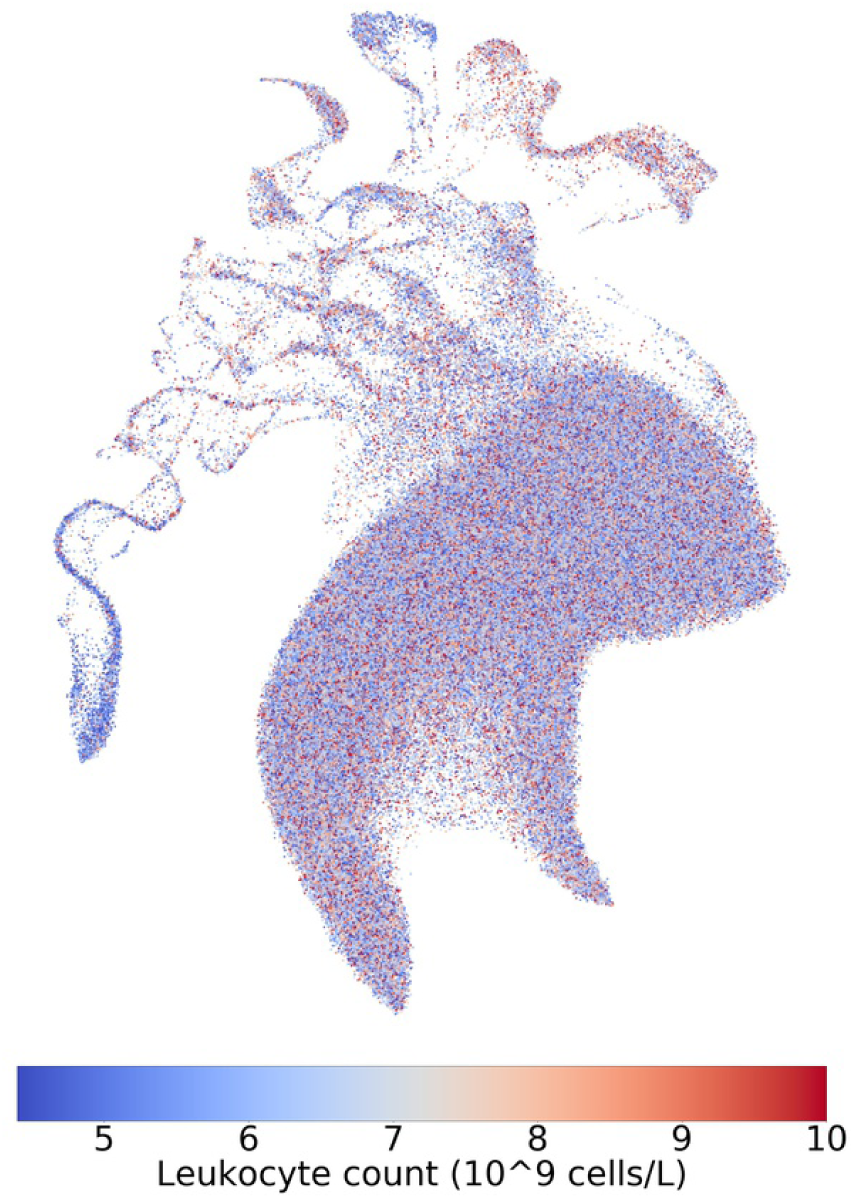
PCA-UMAP on the top 10 principal components of the UKBB colored by leukocyte count (female). Data has been randomized as explained in the materials and methods section.

**Fig. S16.**
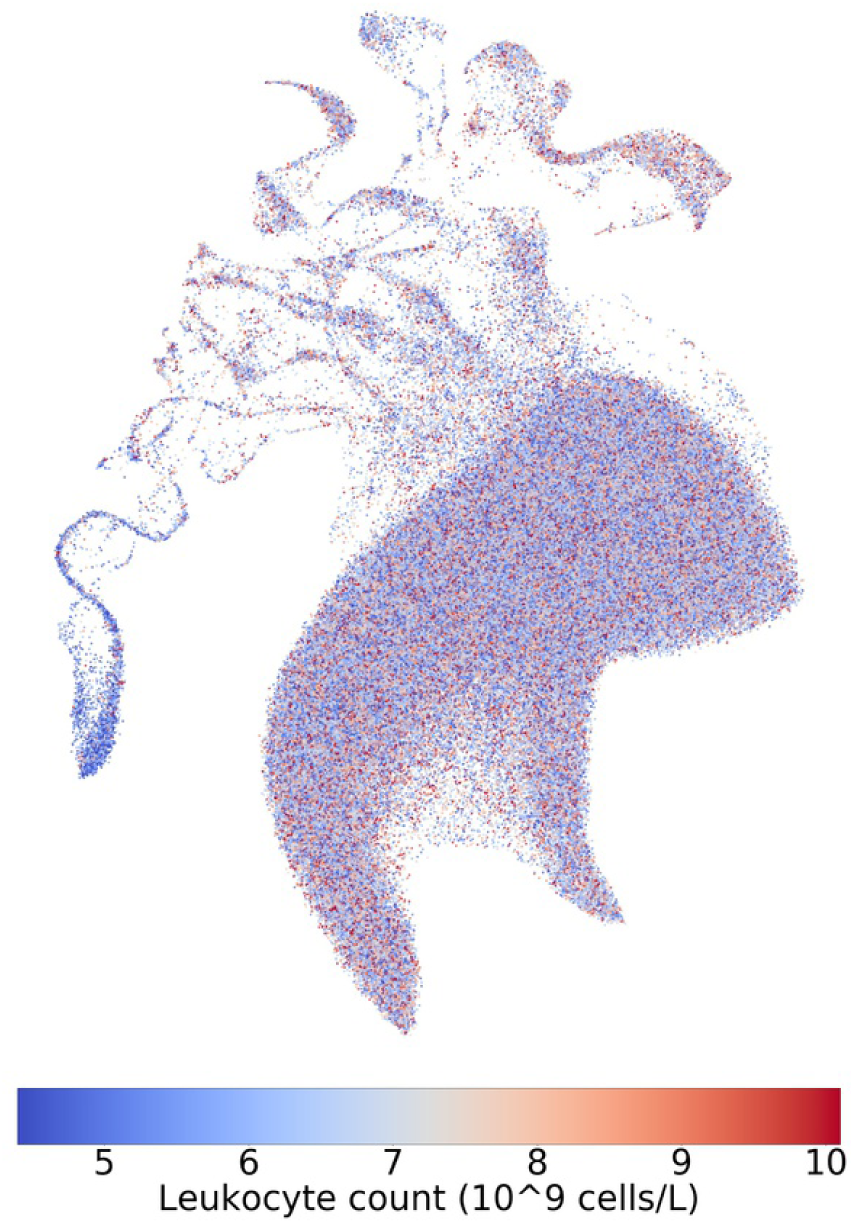
PCA-UMAP on the top 10 principal components of the UKBB colored by leukocyte count (male). Data has been randomized as explained in the materials and methods section.

**Fig. S17.**
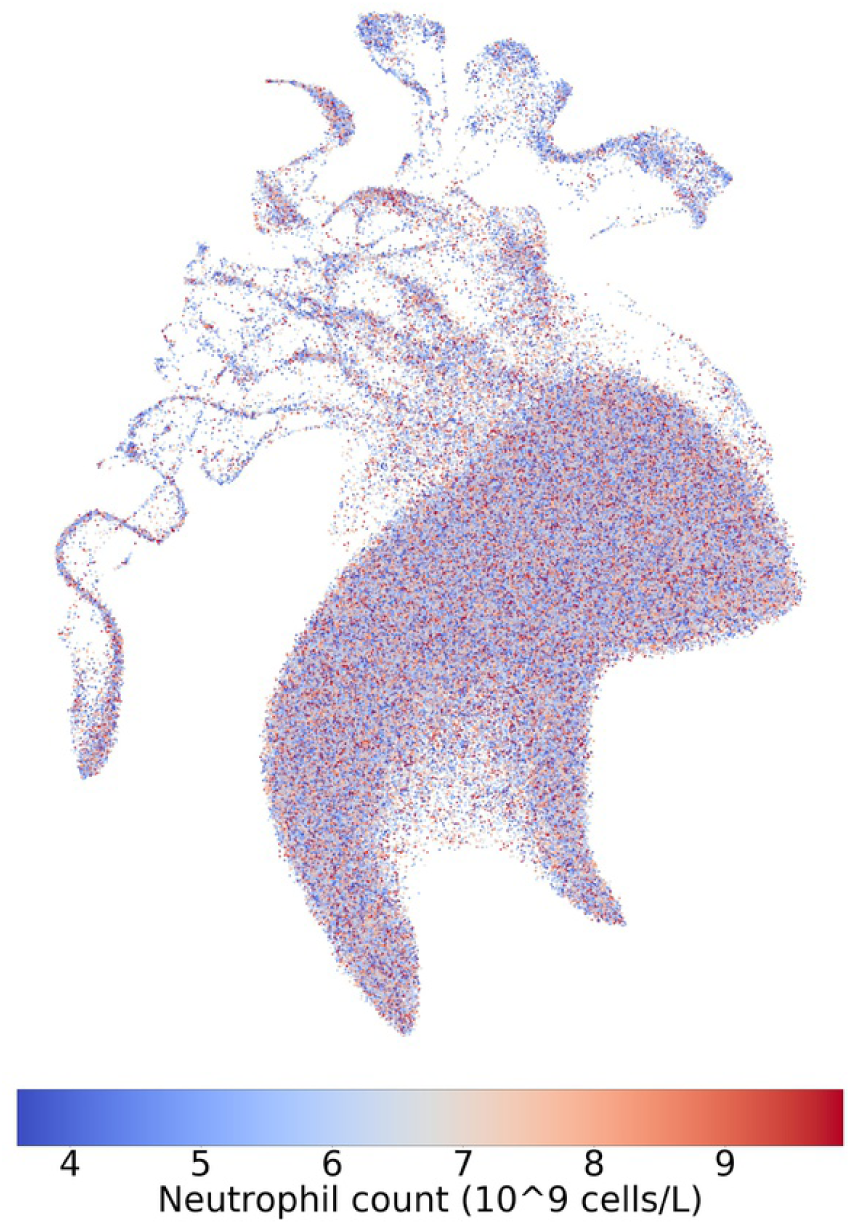
PCA-UMAP on the top 10 principal components of the UKBB colored by neutrophil count (female). Data has been randomized as explained in the materials and methods section.

**Fig. S18.**
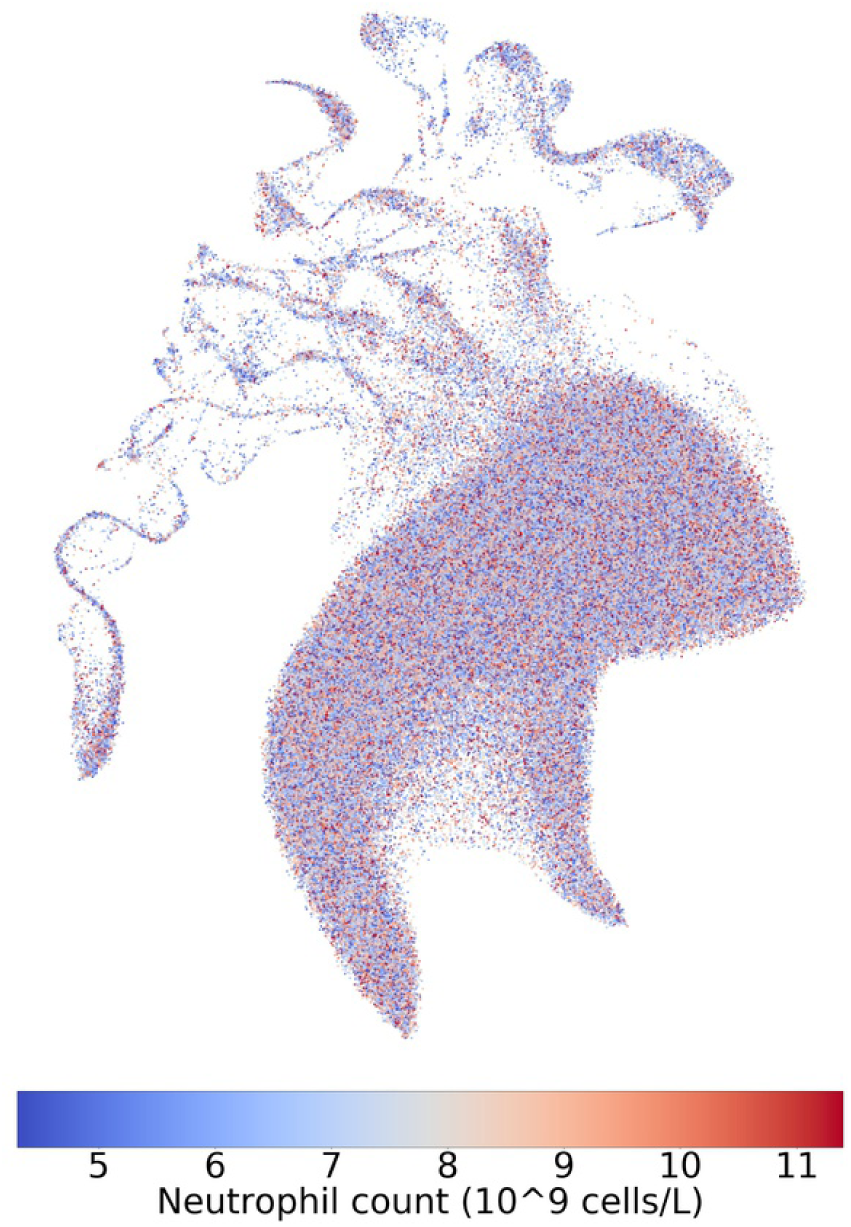
PCA-UMAP on the top 10 principal components of the UKBB colored by neutrophil count (male). Data has been randomized as explained in the materials and methods section.

**Fig. S19.**
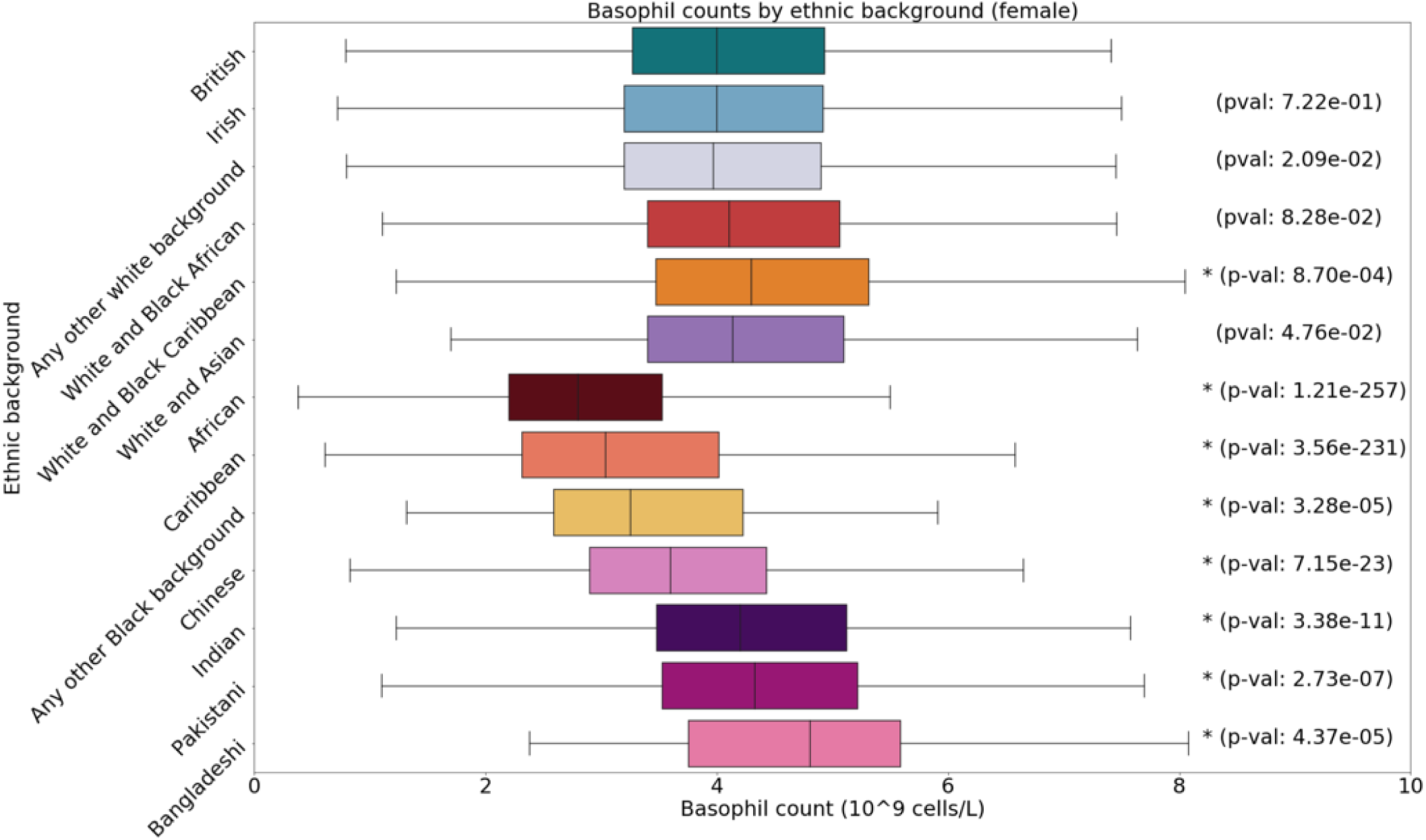
Boxplot of basophil counts by sex and ethnic group, annotated with p-values. Asterisks indicate significant difference from the White British group with a Bonferroni correction for 12 groups.

**Fig. S20.**
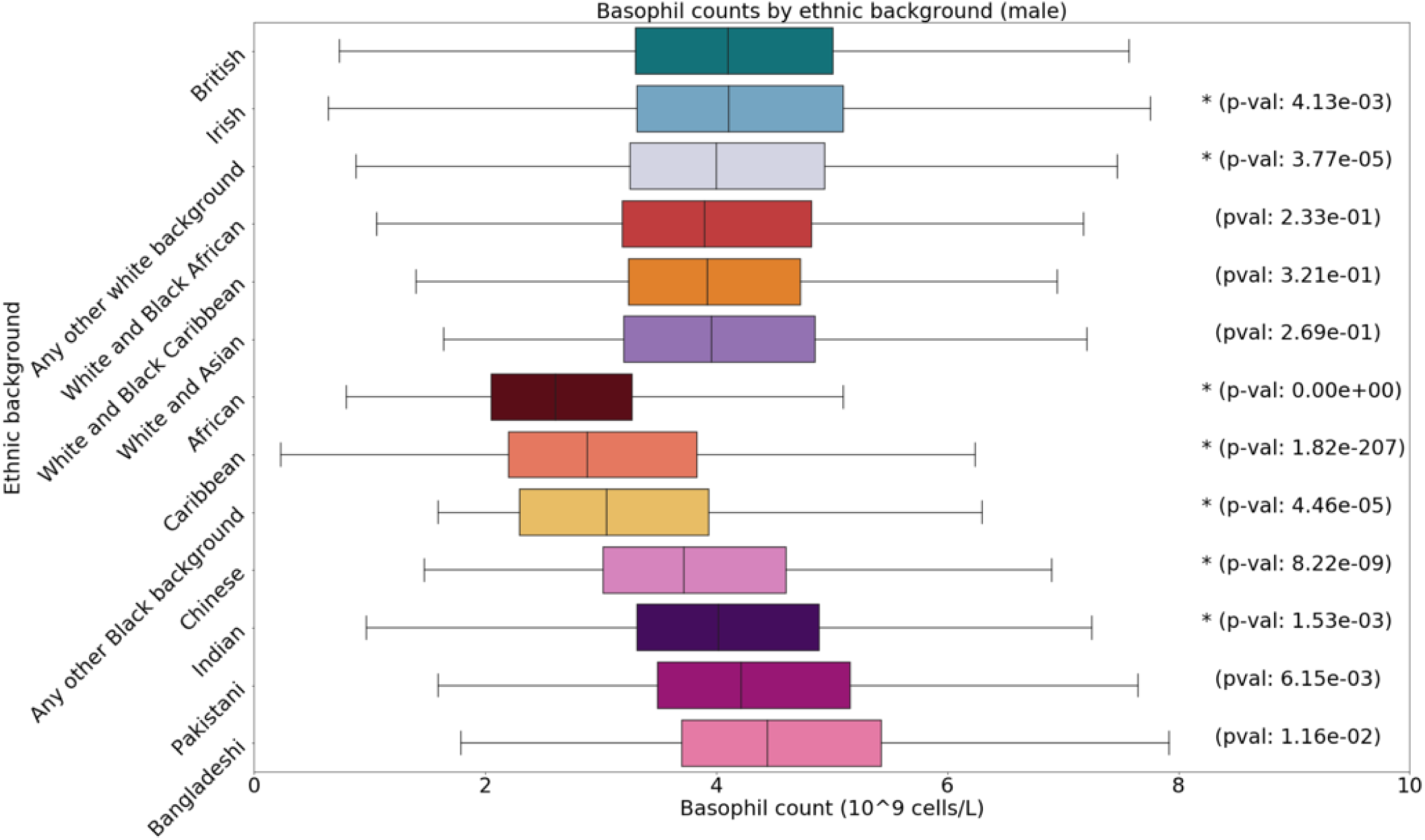
Boxplot of basophil counts by sex and ethnic group, annotated with p-values. Asterisks indicate significant difference from the White British group with a Bonferroni correction for 12 groups.

**Fig. S21.**
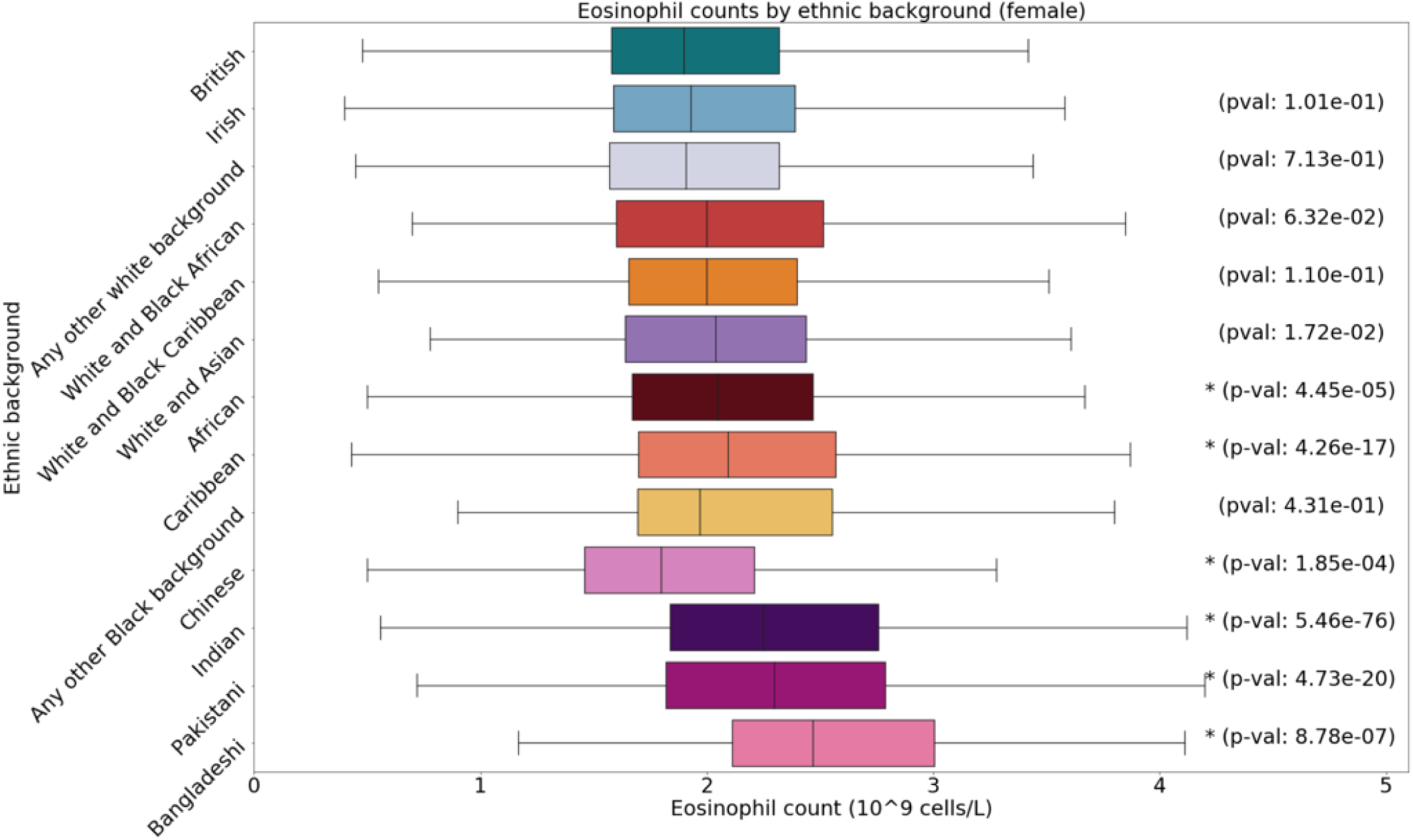
Boxplot of eosinophil counts by sex and ethnic group, annotated with p-values. Asterisks indicate significant difference from the White British group with a Bonferroni correction for 12 groups.

**Fig. S22.**
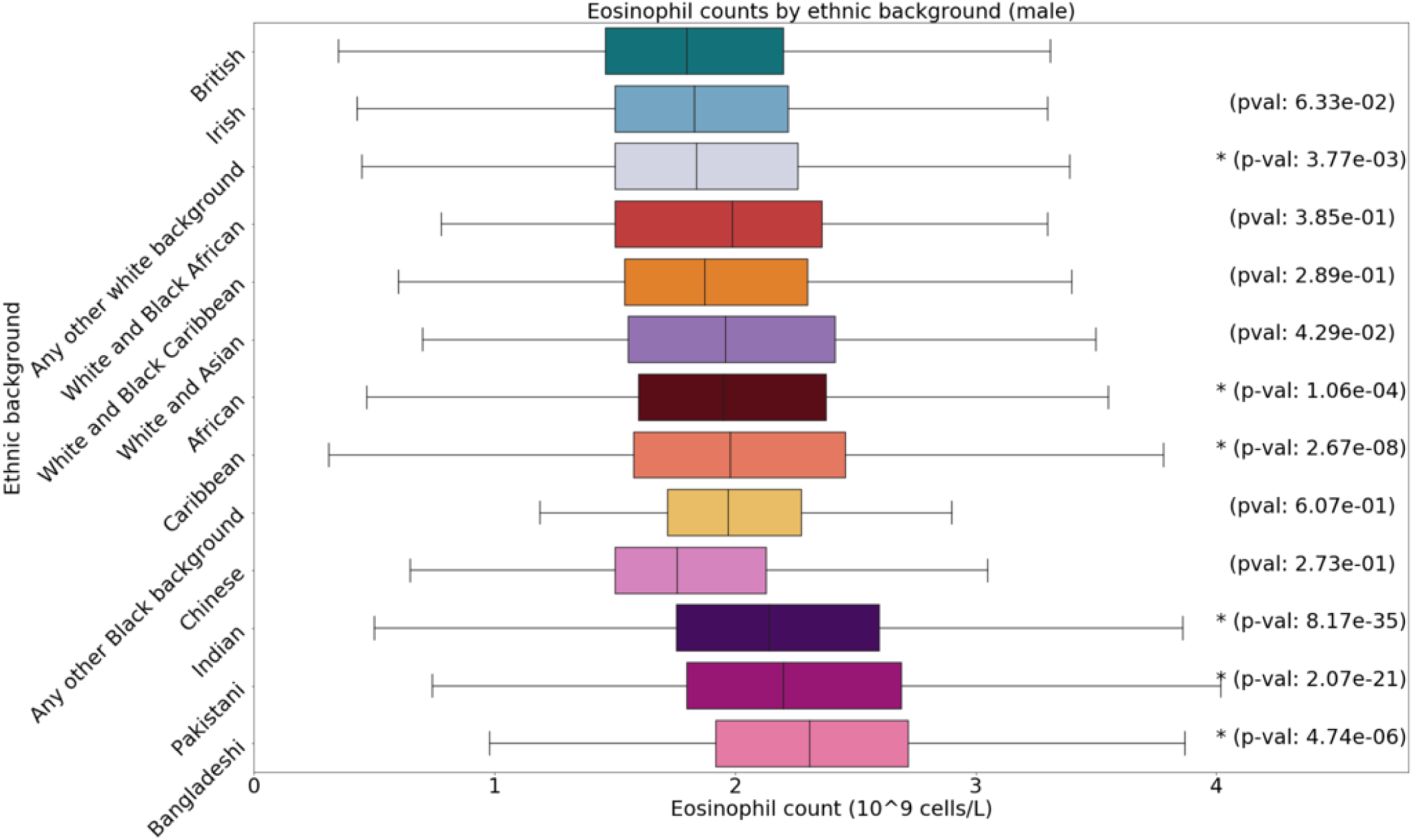
Boxplot of eosinophil counts by sex and ethnic group, annotated with p-values. Asterisks indicate significant difference from the White British group with a Bonferroni correction for 12 groups.

**Fig. S23.**
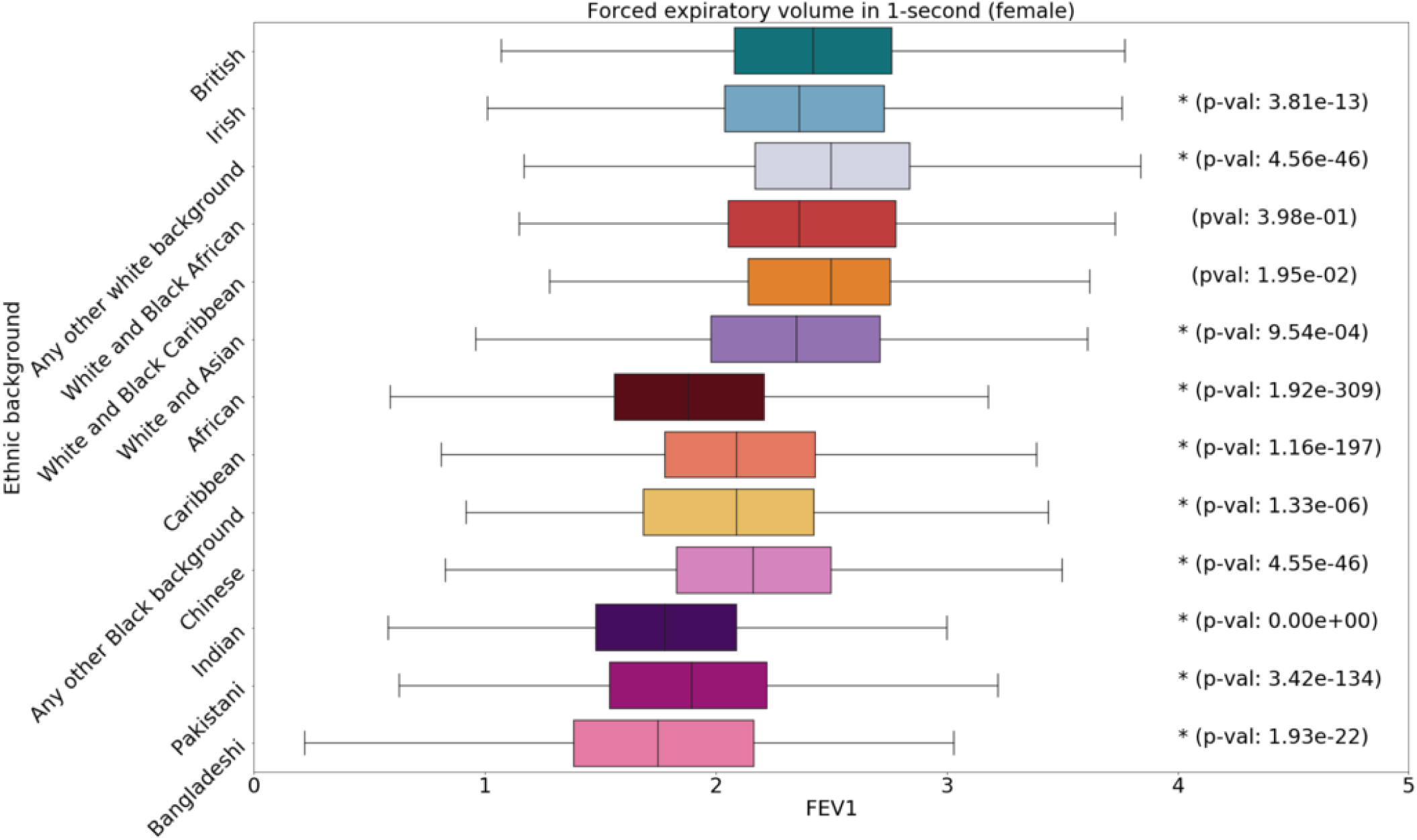
Boxplot of FEV1 by sex and ethnic group, annotated with p-values. Asterisks indicate significant difference from the White British group with a Bonferroni correction for 12 groups.

**Fig. S24.**
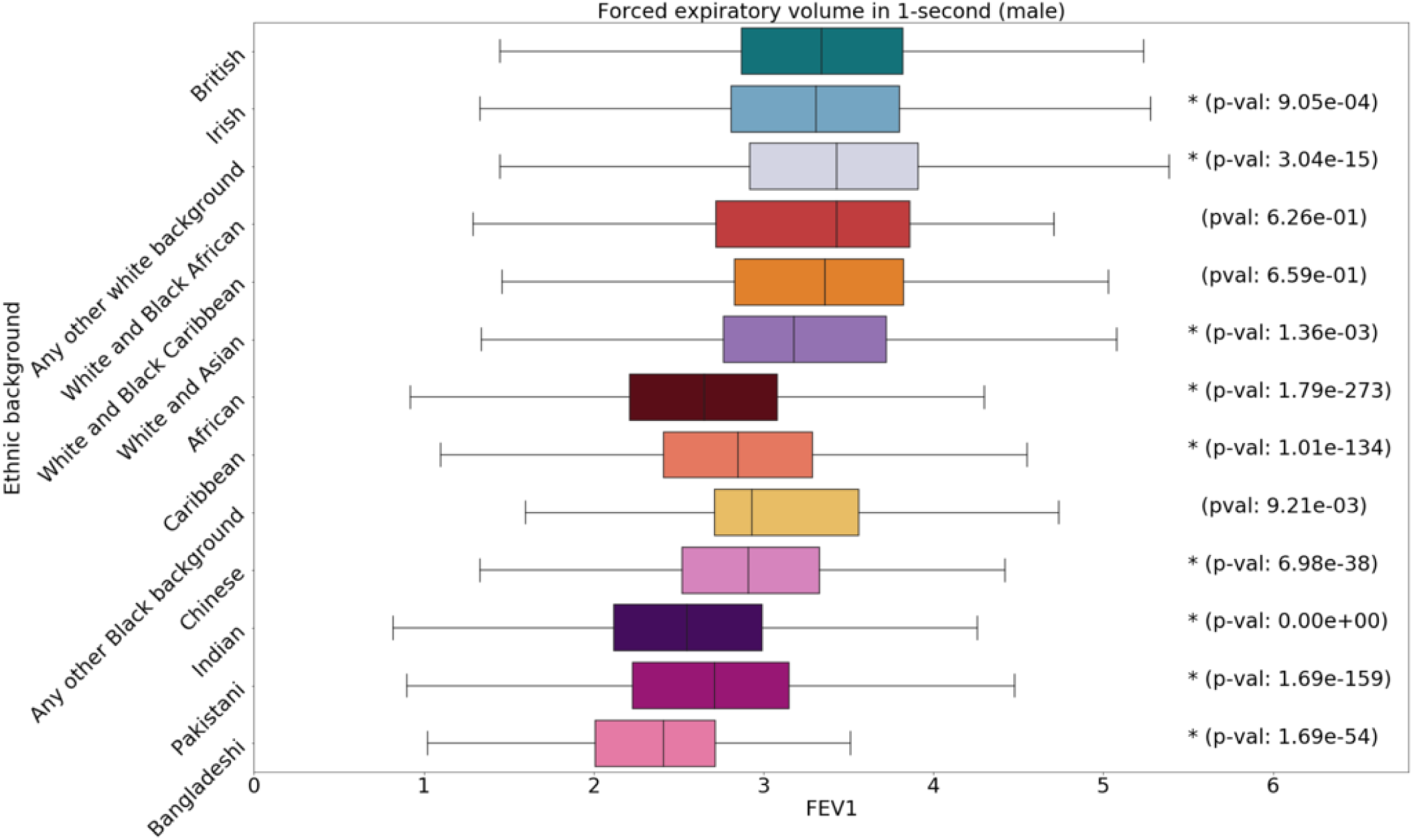
Boxplot of FEV1 by sex and ethnic group, annotated with p-values. Asterisks indicate significant difference from the White British group with a Bonferroni correction for 12 groups.

**Fig. S25.**
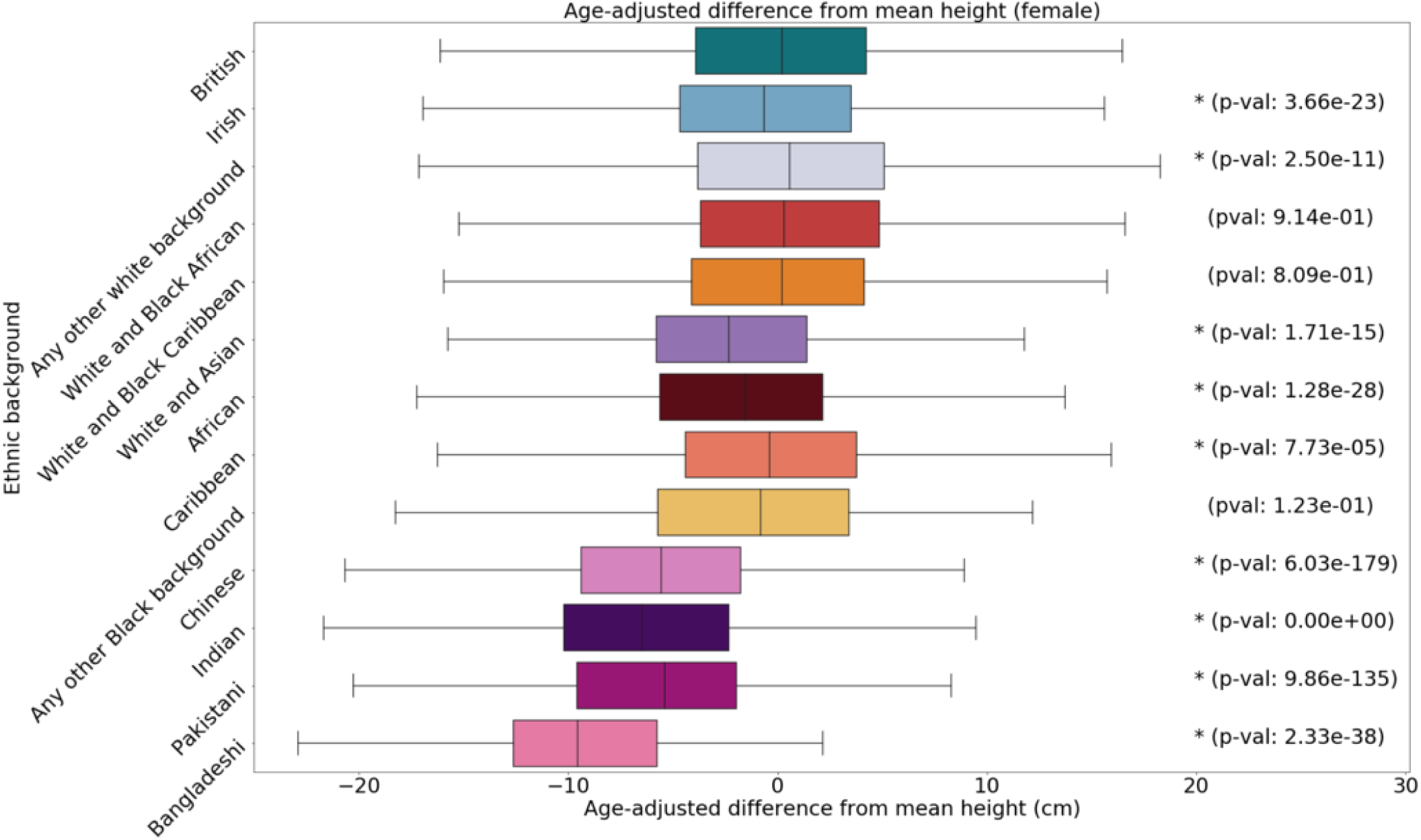
Boxplot of height by sex and ethnic group, annotated with p-values. Asterisks indicate significant difference from the White British group with a Bonferroni correction for 12 groups.

**Fig. S26.**
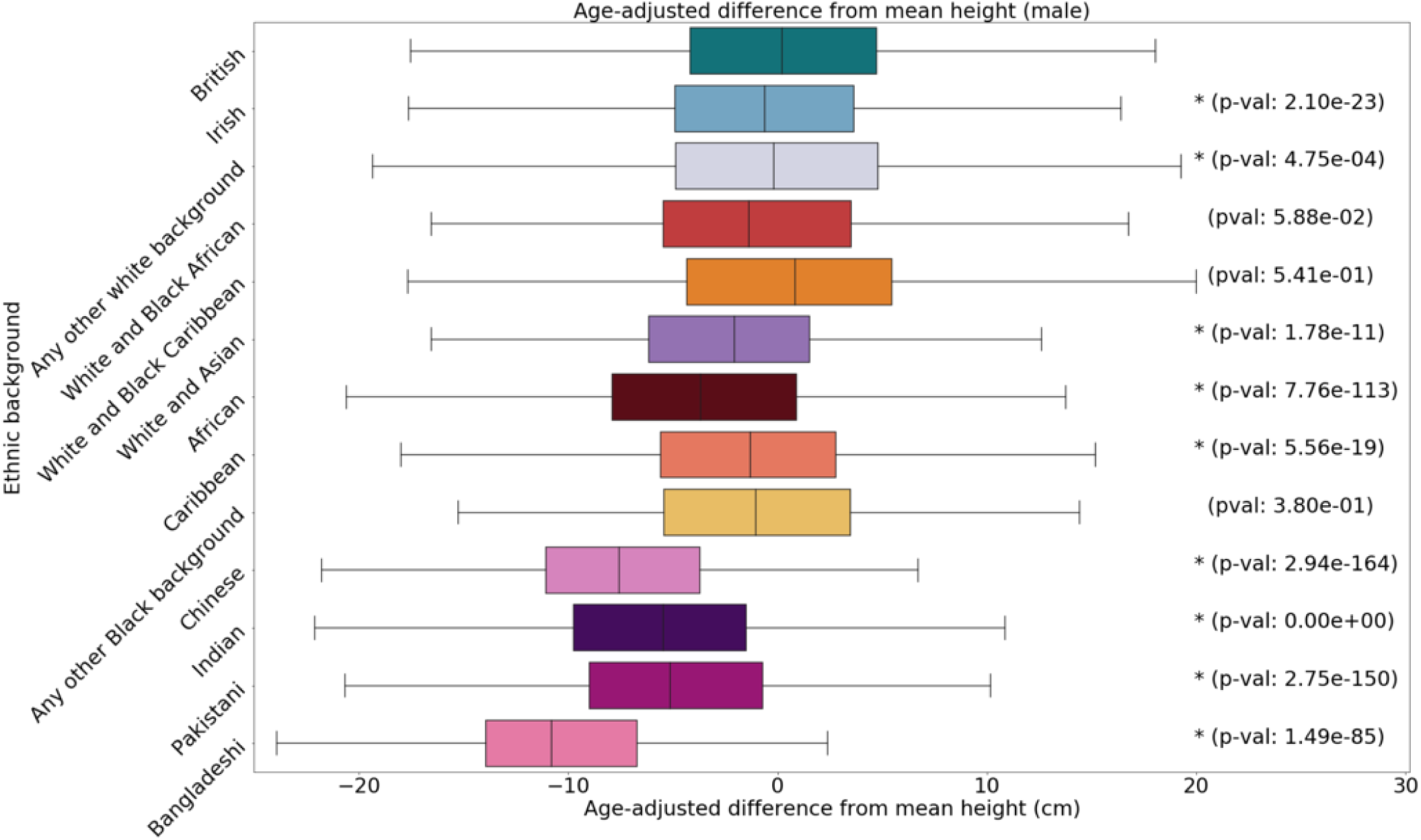
Boxplot of height by sex and ethnic group, annotated with p-values. Asterisks indicate significant difference from the White British group with a Bonferroni correction for 12 groups.

**Fig. S27.**
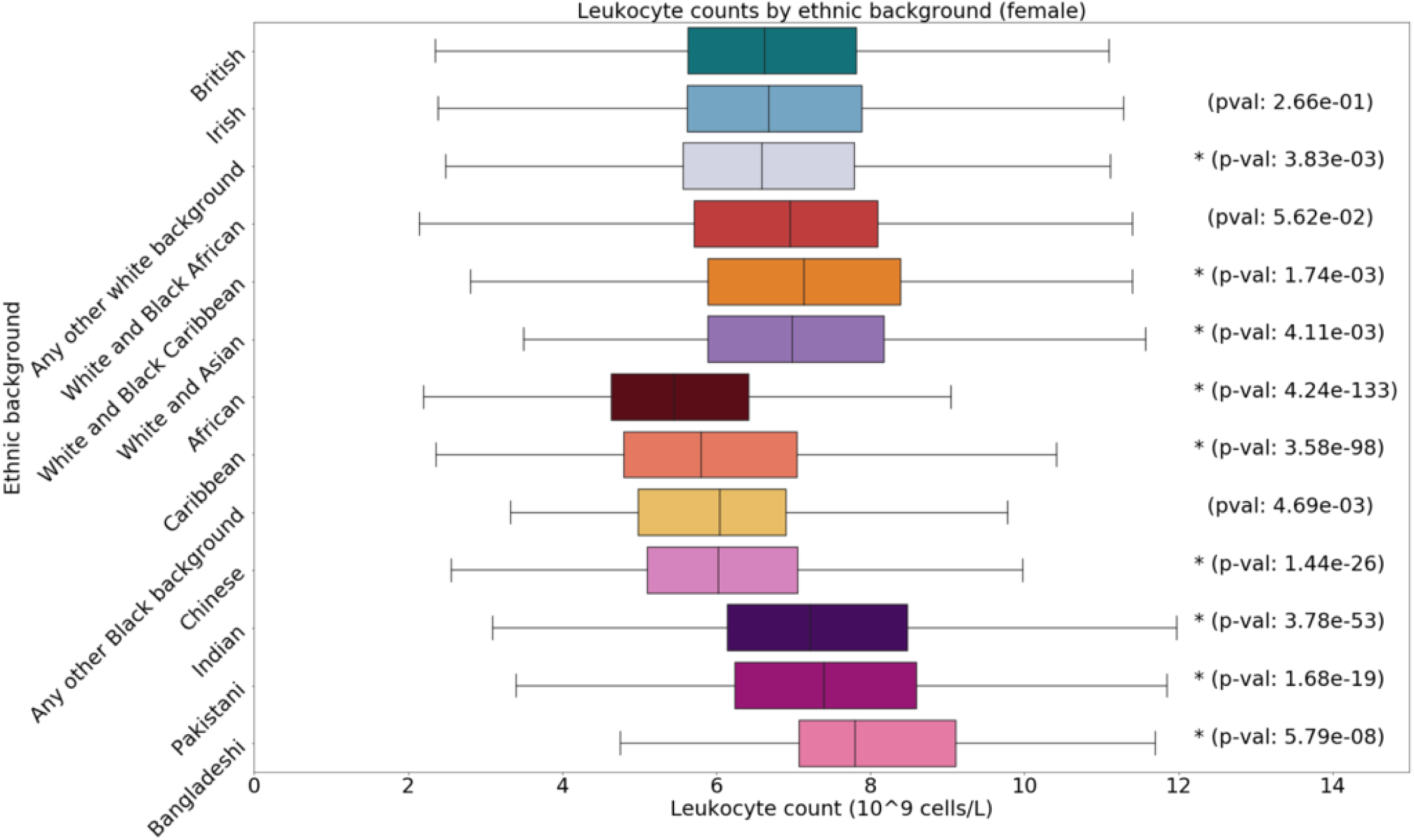
Boxplot of leukocyte counts by sex and ethnic group, annotated with p-values. Asterisks indicate significant difference from the White British group with a Bonferroni correction for 12 groups.

**Fig. S28.**
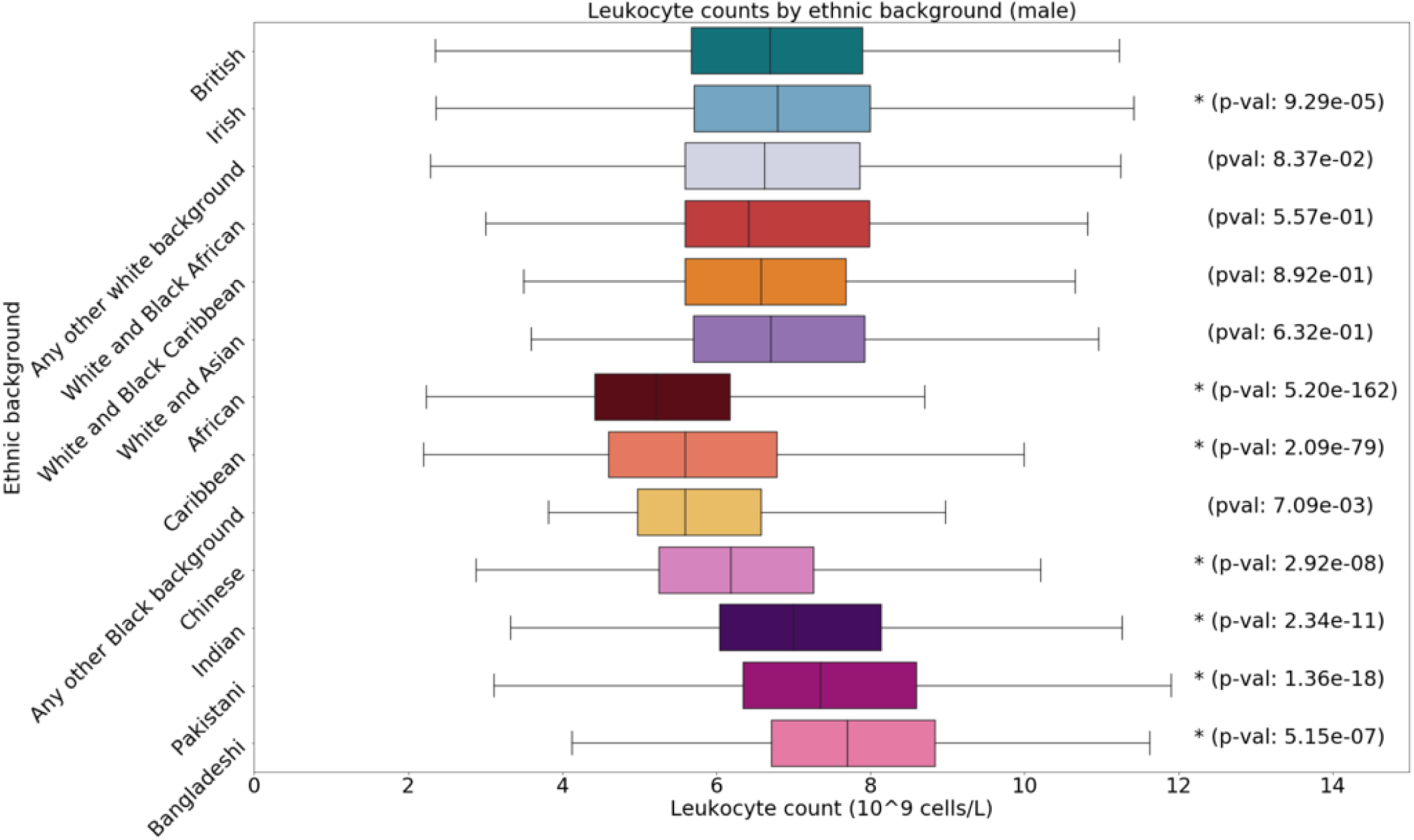
Boxplot of leukocyte counts by sex and ethnic group, annotated with p-values. Asterisks indicate significant difference from the White British group with a Bonferroni correction for 12 groups.

**Fig. S29.**
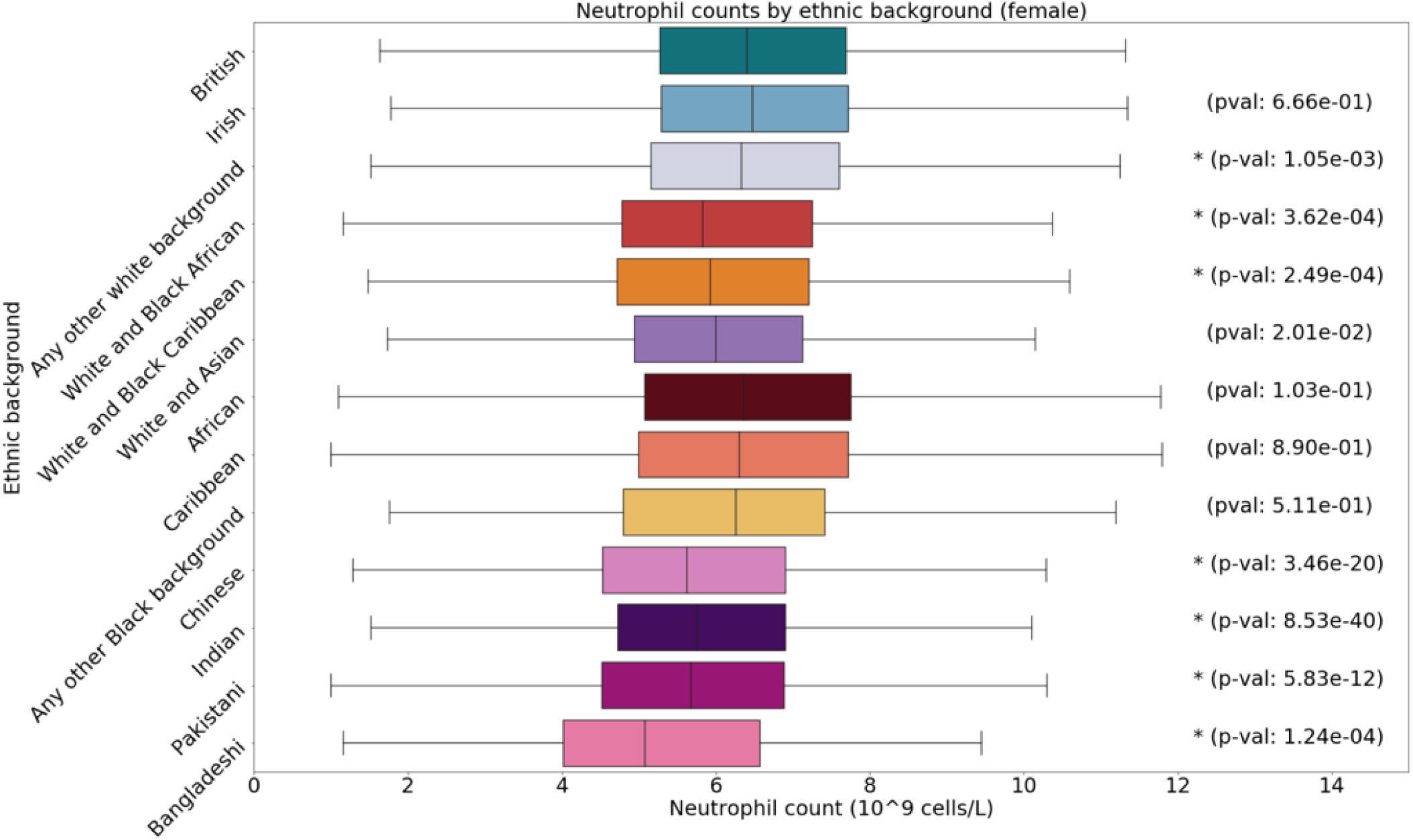
Boxplot of neutrophil counts by sex and ethnic group, annotated with p-values. Asterisks indicate significant difference from the White British group with a Bonferroni correction for 12 groups.

**Fig. S30.**
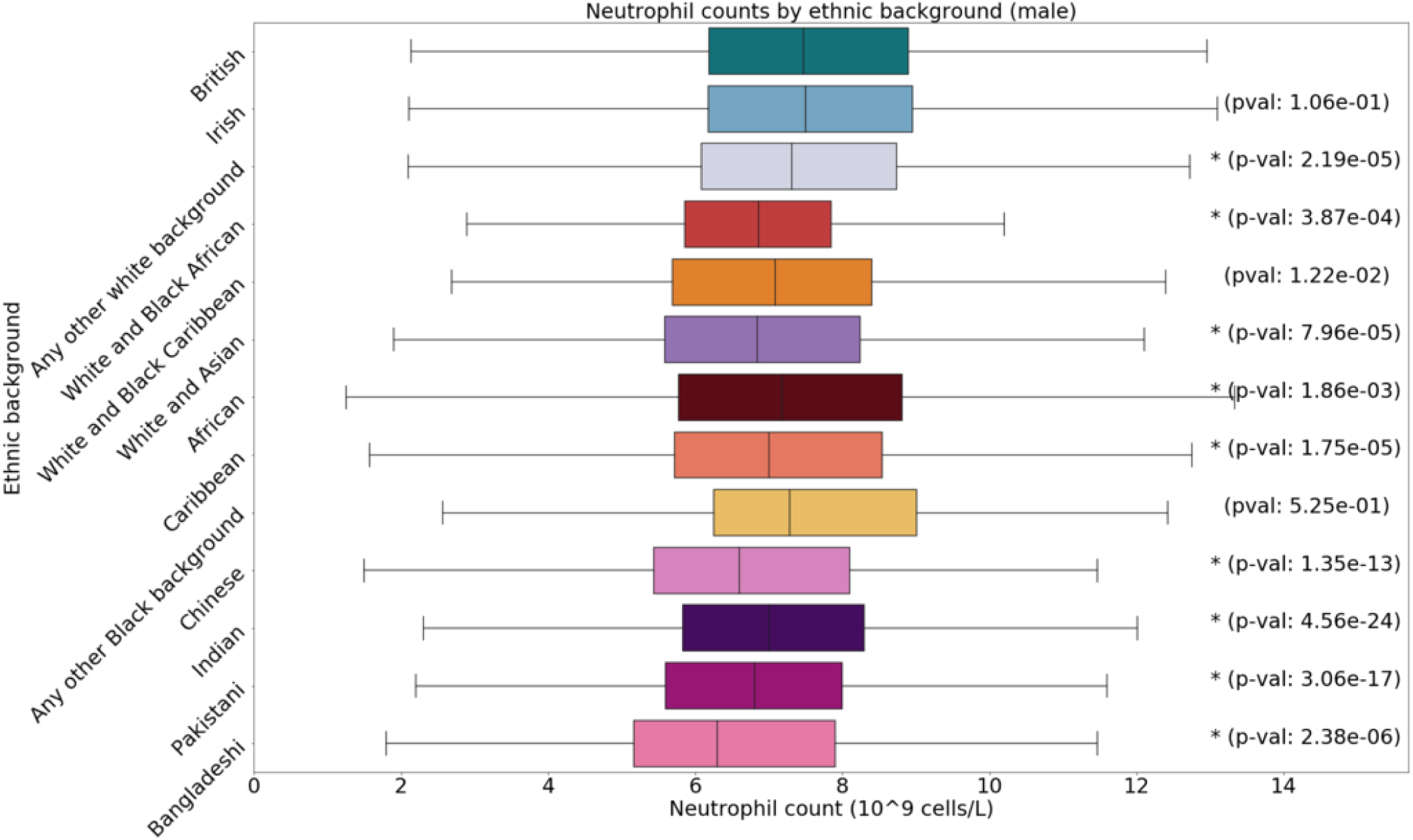
Boxplot of neutrophil counts by sex and ethnic group, annotated with p-values. Asterisks indicate significant difference from the White British group with a Bonferroni correction for 12 groups.

**Fig. S31.**
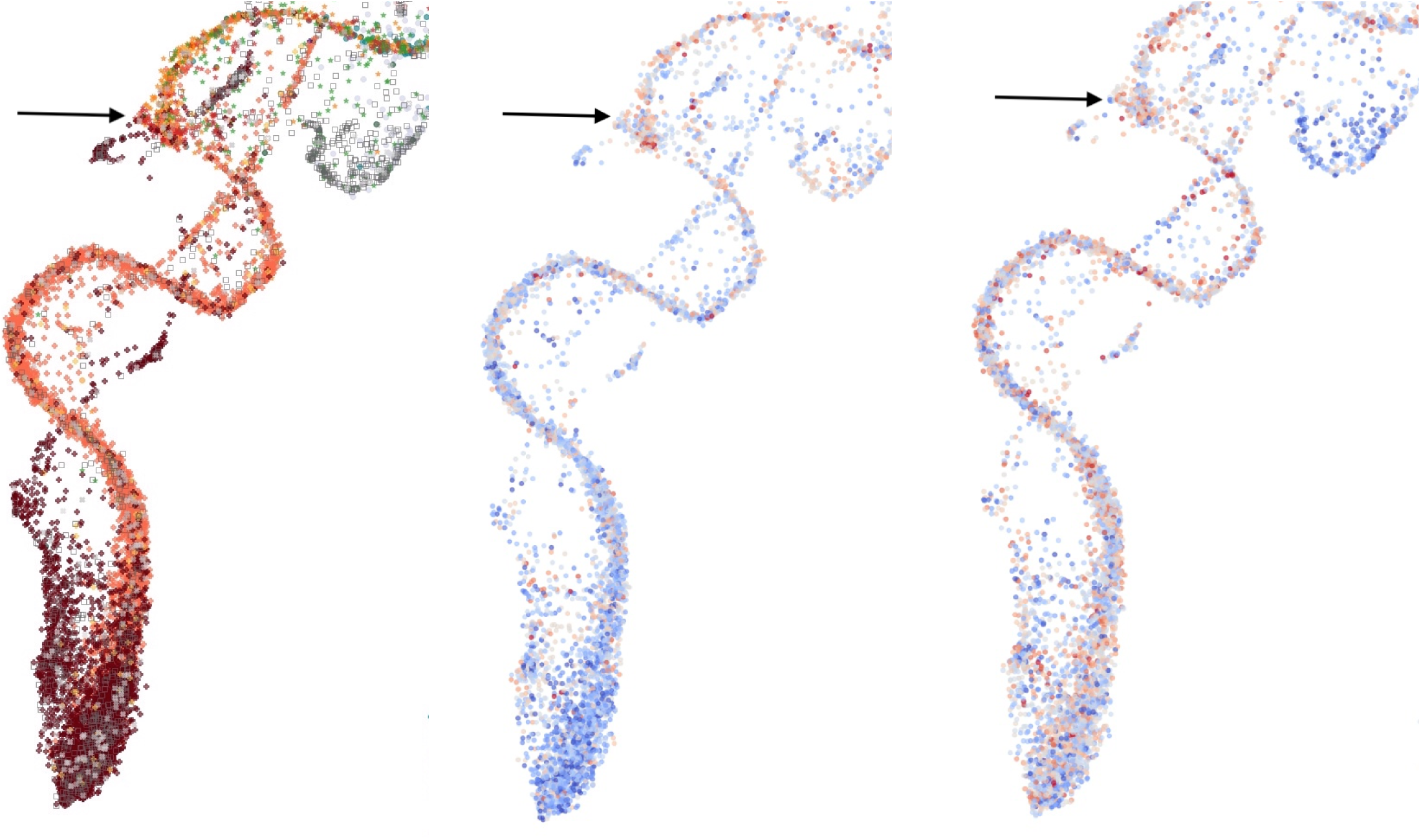
Individuals of Black African, Black Caribbean, and mixed backgrounds (primarily White and Black Caribbean/African) colored by self-identified ethnic background (left, from figure 4b), FEV1 (middle), and age-adjusted height (right). An arrow points to an area where the FEV1 distribution appears to change, corresponding to where the clusters contain more people with self-identified mixed backgrounds.

**Fig. S32.**
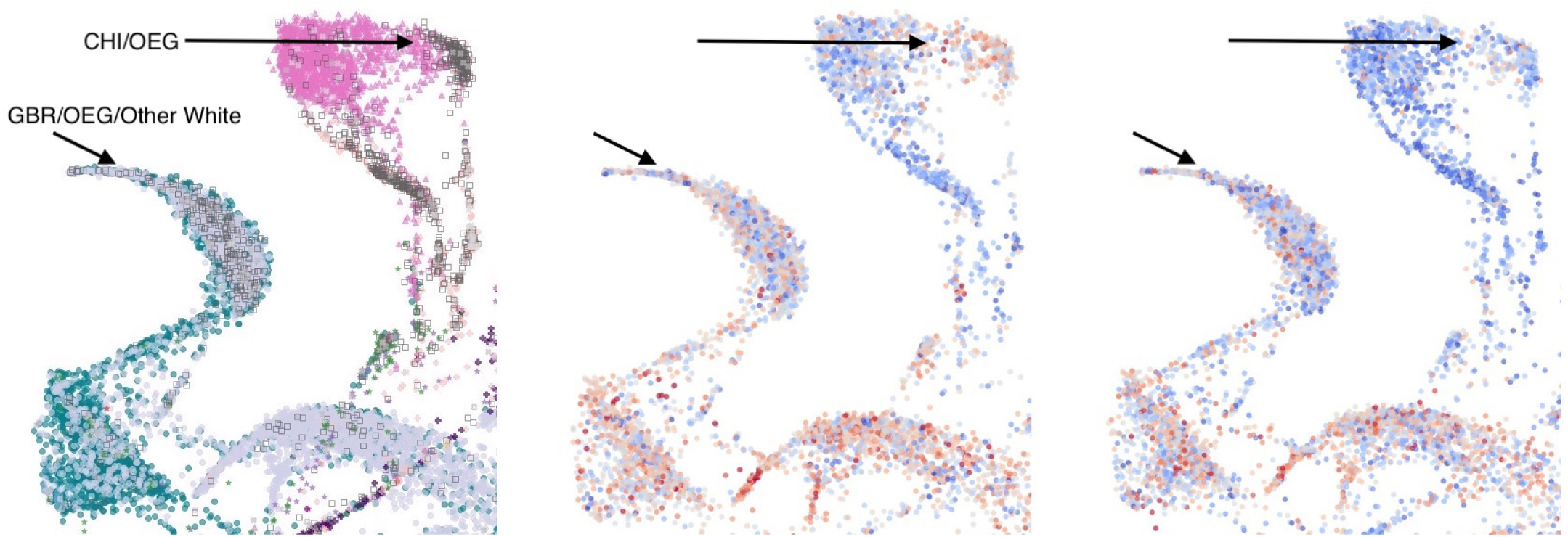
Zoomed in section of figure 4b focused on individuals with Chinese (CHI), White British (GBR), any other white background, or any other ethnic group (OEG) colored by ethnicity (left), FEV1 (middle), and age-adjusted height (right). The OEG cluster next to the Chinese cluster is colored differently, suggesting this population may have different FEV1 characteristics. A cluster of OEG/other white individuals is more blue, suggesting they may have lower than average FEV1 values relative to the rest of the British or white population.

**Fig. S33.**
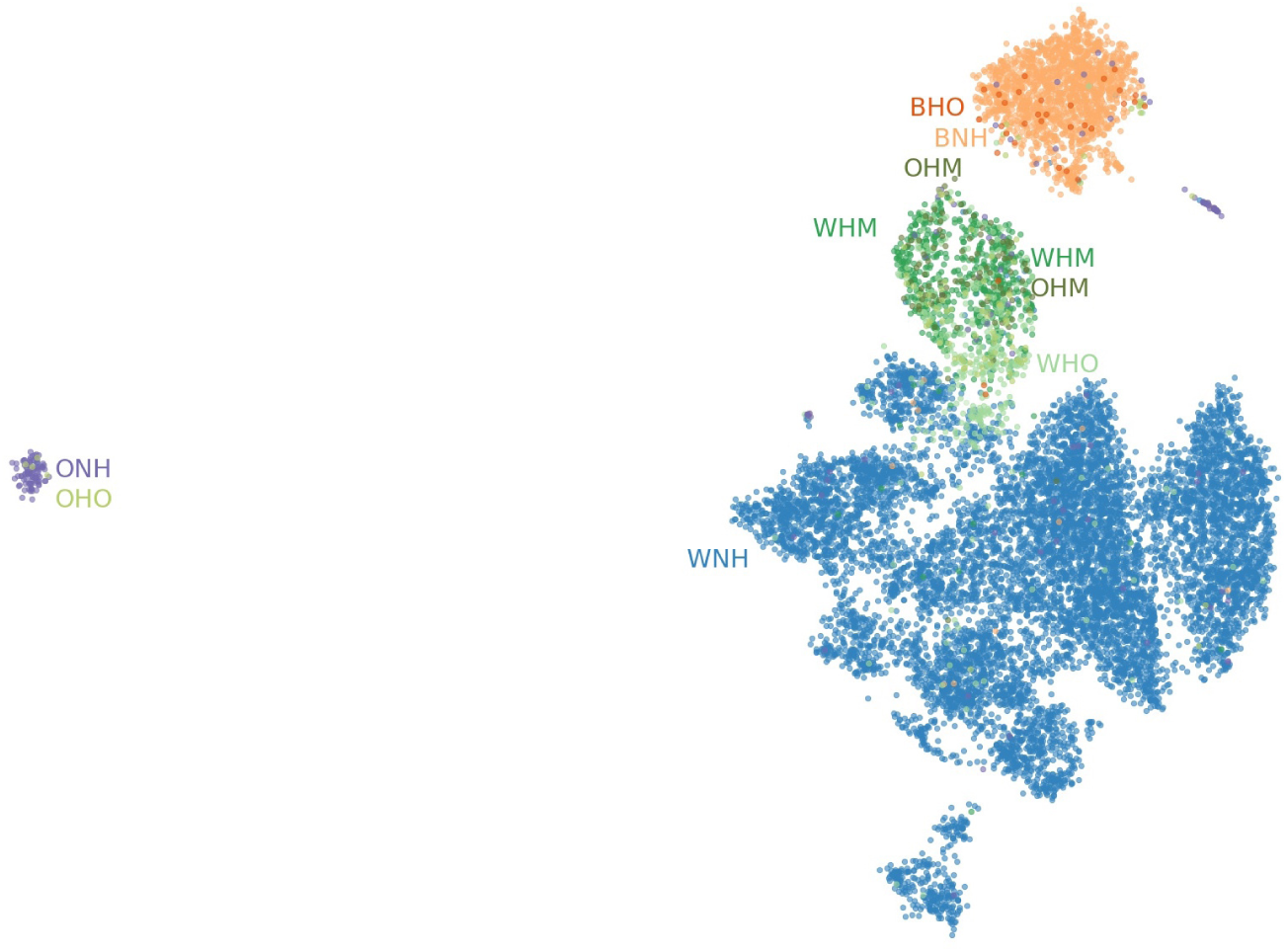
UMAP applied to the first 10 principal components of HRS data. Points colored by self-identified race, Hispanic status, and Mexican-American status. The cluster on the left is mostly people who identify as neither Black nor White and were born outside the contiguous United States or in the Pacific census region. Clustering with the 1KGP data places them with Asian-identified populations. BNH, Black (not Hispanic); BHO, Black (Hispanic, Other); WNH, White (not Hispanic); WHM, White (Hispanic, Mexican-American); WHO, White Hispanic (Other); ONH, Other (not Hispanic); OHM, Other (Hispanic, Mexican-American); OHO, Other (Hispanic, Other).

**Fig. S34.**
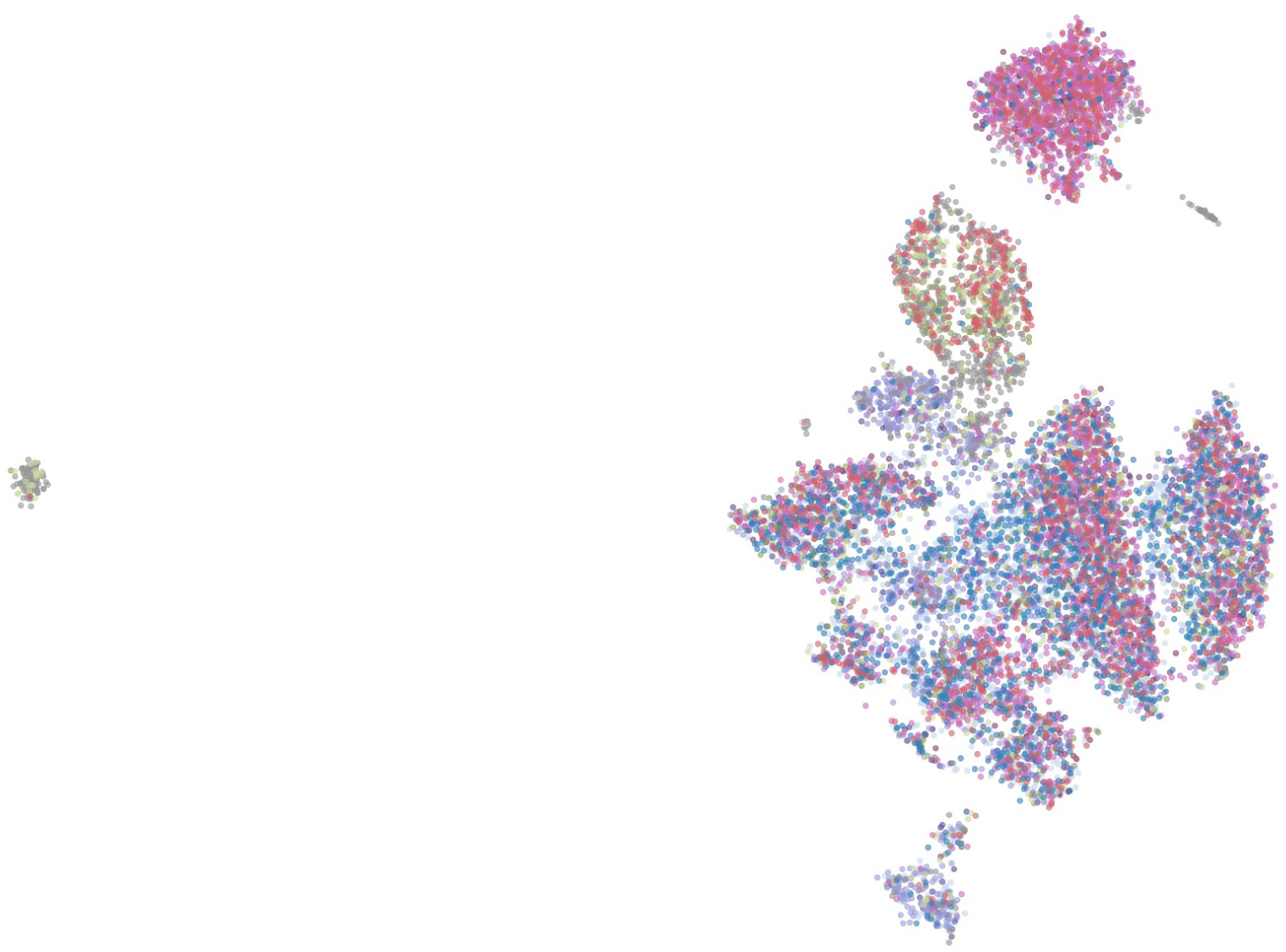
UMAP on the top 10 principal components of the HRS dataset, colored by Census Bureau birth region. There is no obvious pattern in the clusters of majority “White Not Hispanic” individuals.

**Fig. S35.**
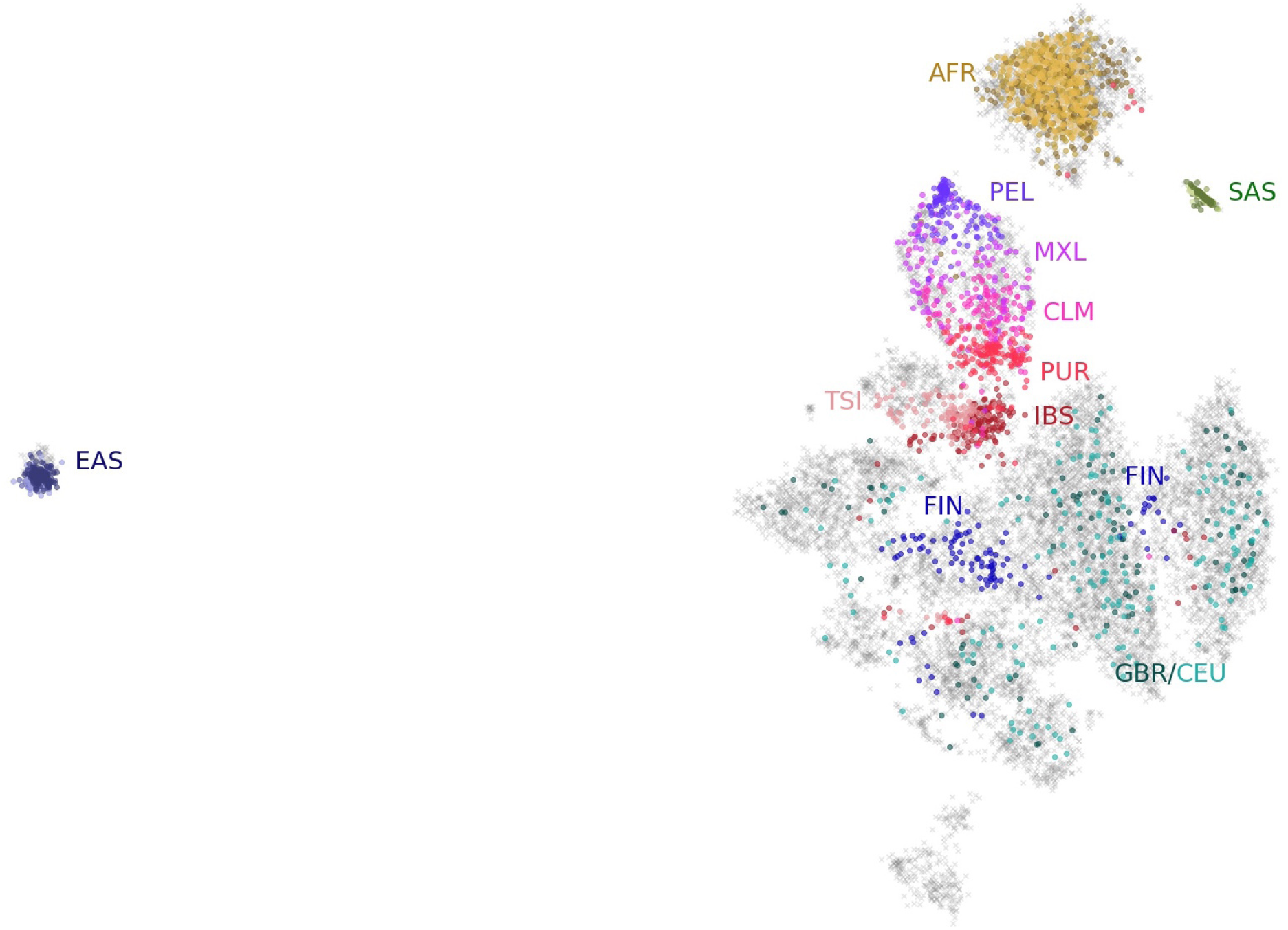
UMAP on the top 10 principal components of the HRS data, with 1KGP data projected onto the embedding. Individuals from the HRS are grey. British (GBR) and other European (CEU) individuals are scattered throughout the “White Not Hispanic” clusters. Finns (FIN) form clear groupings. Spanish (IBS) and Italian (TSI) individuals cluster near the Hispanic grouping. There are sub-groups in the Hispanic cluster formed of Puerto Ricans (PUR), Colombians (CLM), Mexicans (MXL), and Peruvians (PEL). Populations with African ancestry (AFR) appear with Black individuals. East Asian (EAS) populations comprising Chinese, Kinh, and Japanese individuals cluster together with what appears in figure 2 as a population of mostly Asian ancestry. South Asian (SAS) populations with Indian, Pakistani, and Sri Lankan ancestry cluster in a separate area. One “White Not Hispanic” cluster at the bottom does not cluster with any 1KGP populations.

**Fig. S36.**
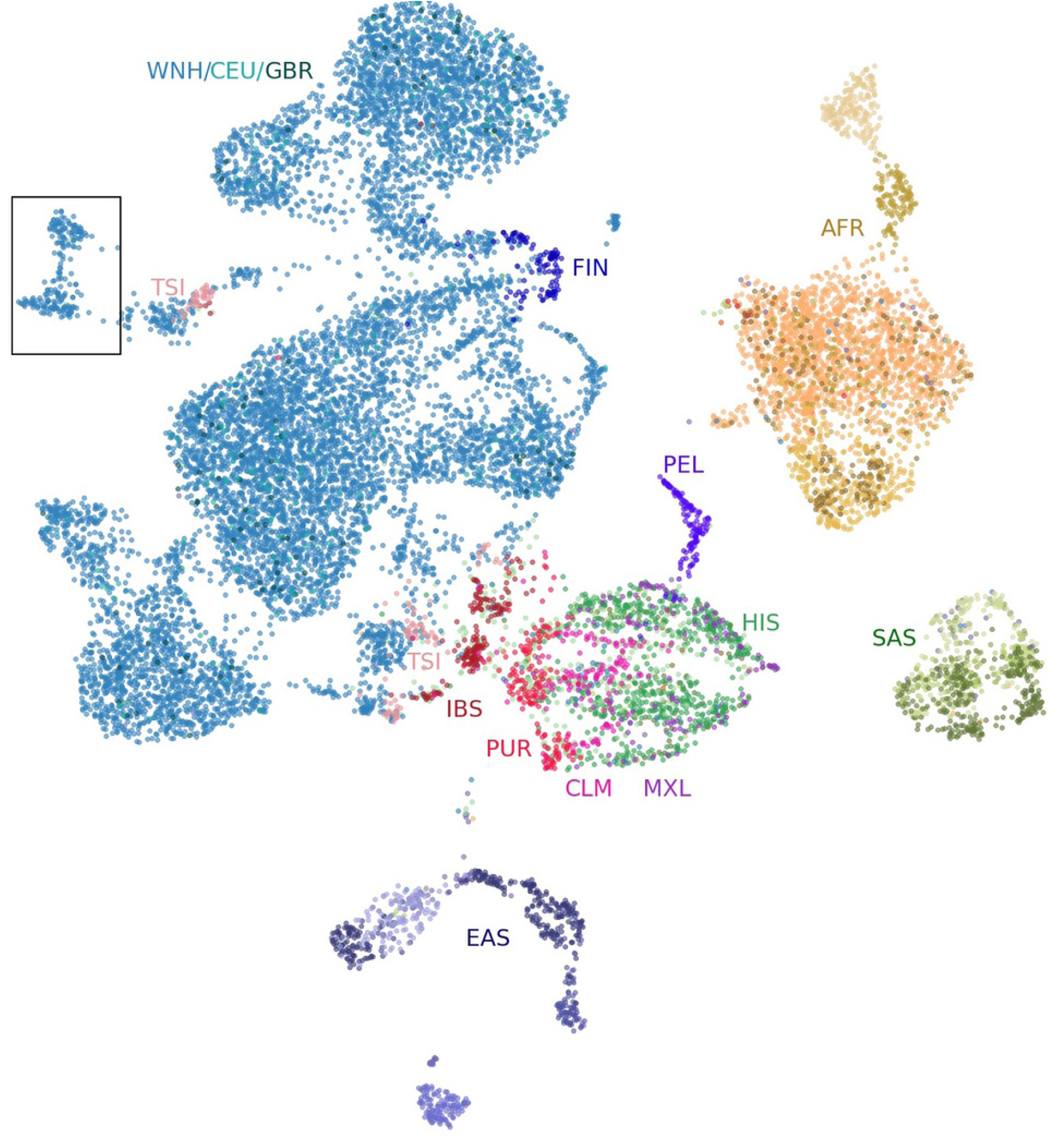
UMAP projection of the top 10 principal components of the combined HRS and 1KGP datasets. One cluster (in the box) does not group with any of the 1KGP populations. A cluster of Finnish (FIN) individuals consistently appears in the “White Not Hispanic” (WNH) group. Groups of Central and South American populations from the 1KGP (CLM, Colombian; MXL, Mexican; PEL, Peruvian; PUR, Puerto Rican) form nearby or within the HRS Hispanic cluster (HIS). Iberian individuals (IBS) cluster near the Hispanic population. Toscani individuals (TSI) form some small clusters and sometimes appear near the Iberian and Hispanic populations. Individuals with British/Scottish (GBR) or Northern/Western European ancestry (CEU) are scattered throughout the WNH clusters. Individuals with African ancestry from the 1KGP group with Black Americans from the HRS (AFR). Similar population groupings occur with South Asian (SAS) and East Asian (EAS) individuals.

**Fig. S37.**
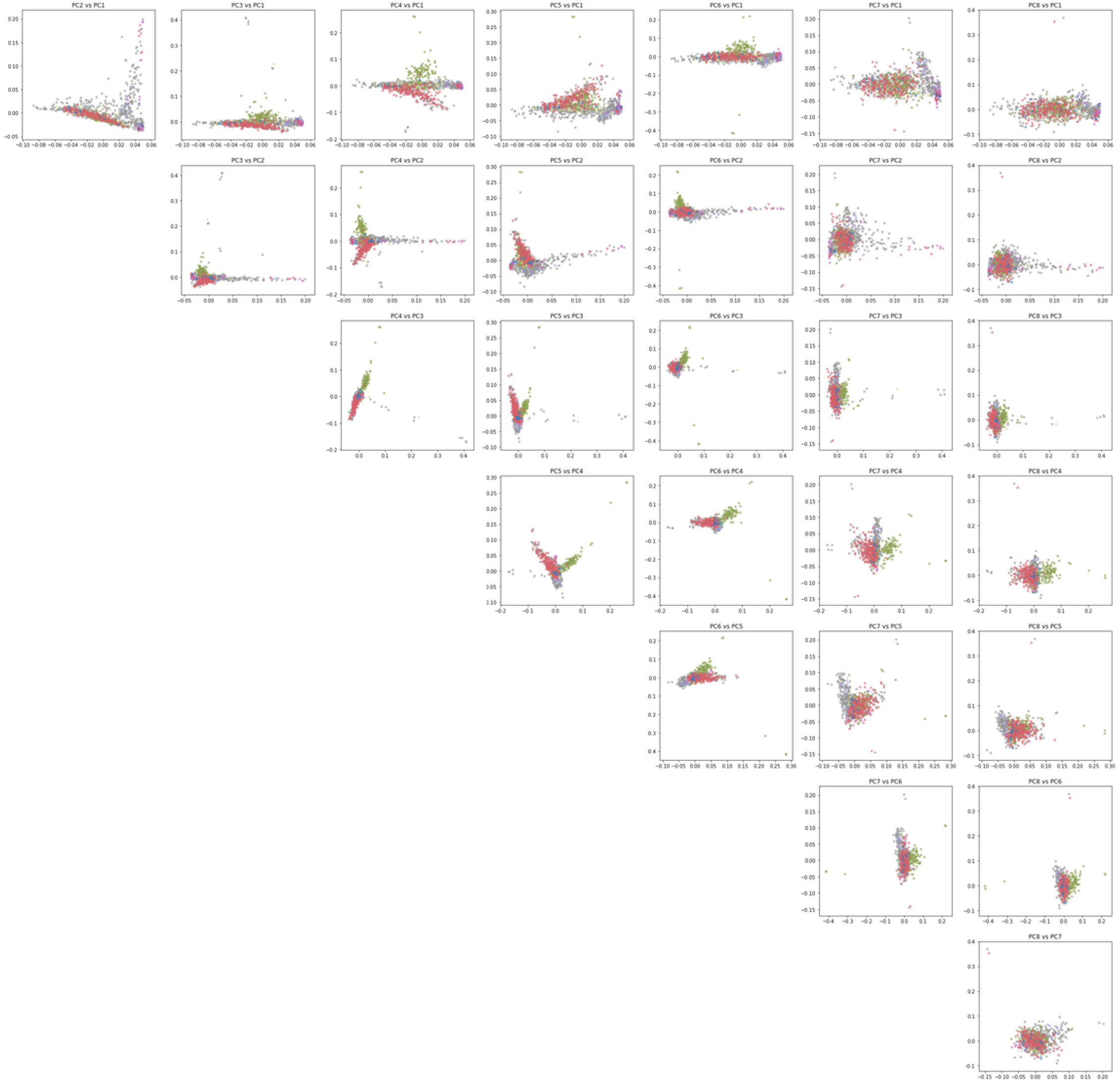
Pairwise plots of the first 8 principal components of the Hispanic subset of the HRS. Those born in the Mountain region are colored green.

**Fig. S38.**
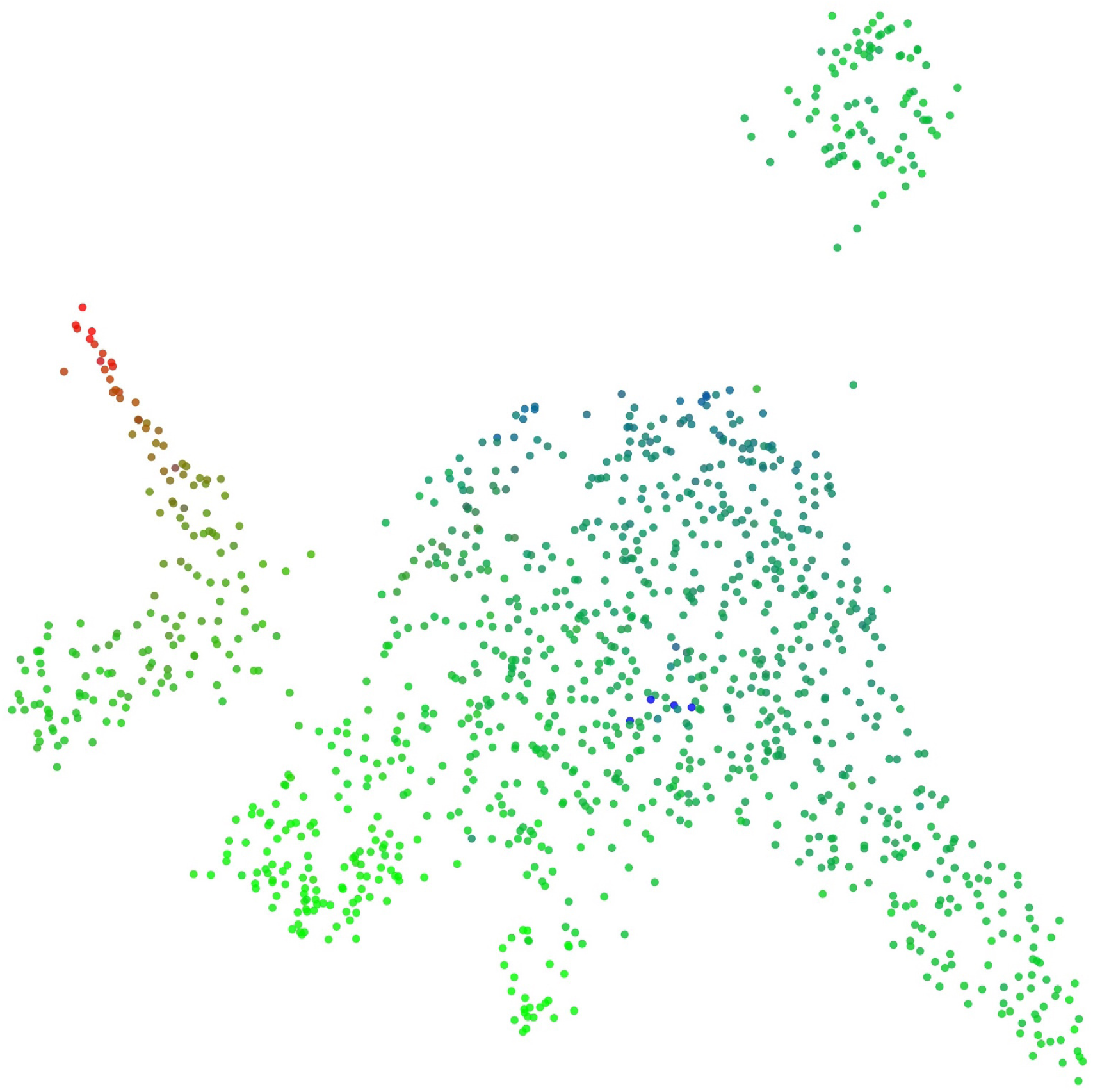
UMAP of the first 7 principal components of the Hispanic population of the HRS, colored by estimated admixture proportions.

**Fig. S39.**
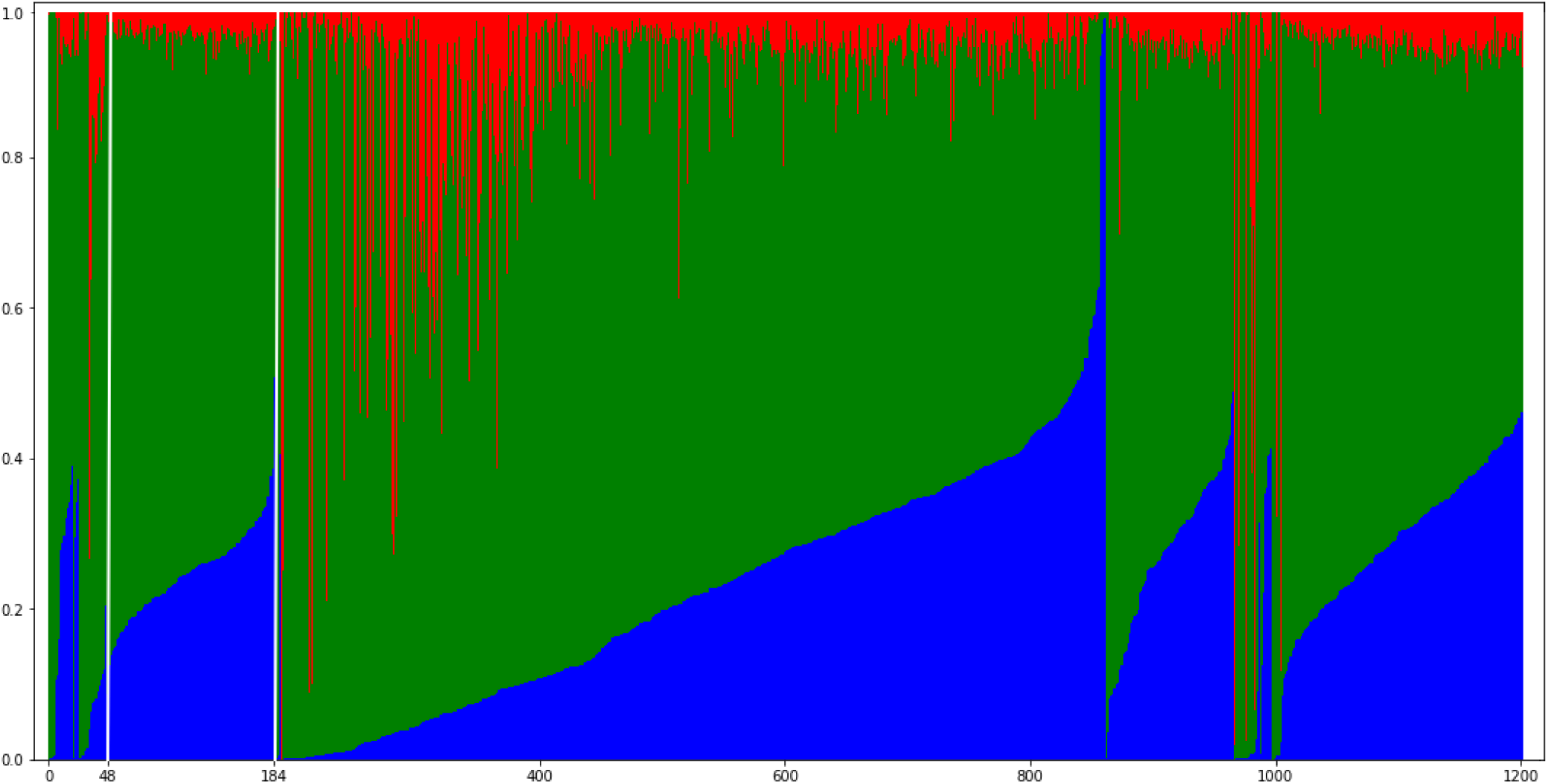
Admixture plot of Hispanic individuals in the HRS. Those born in the Mountain census region fall between the white lines (indices 48 to 184)

**Fig. S40.**
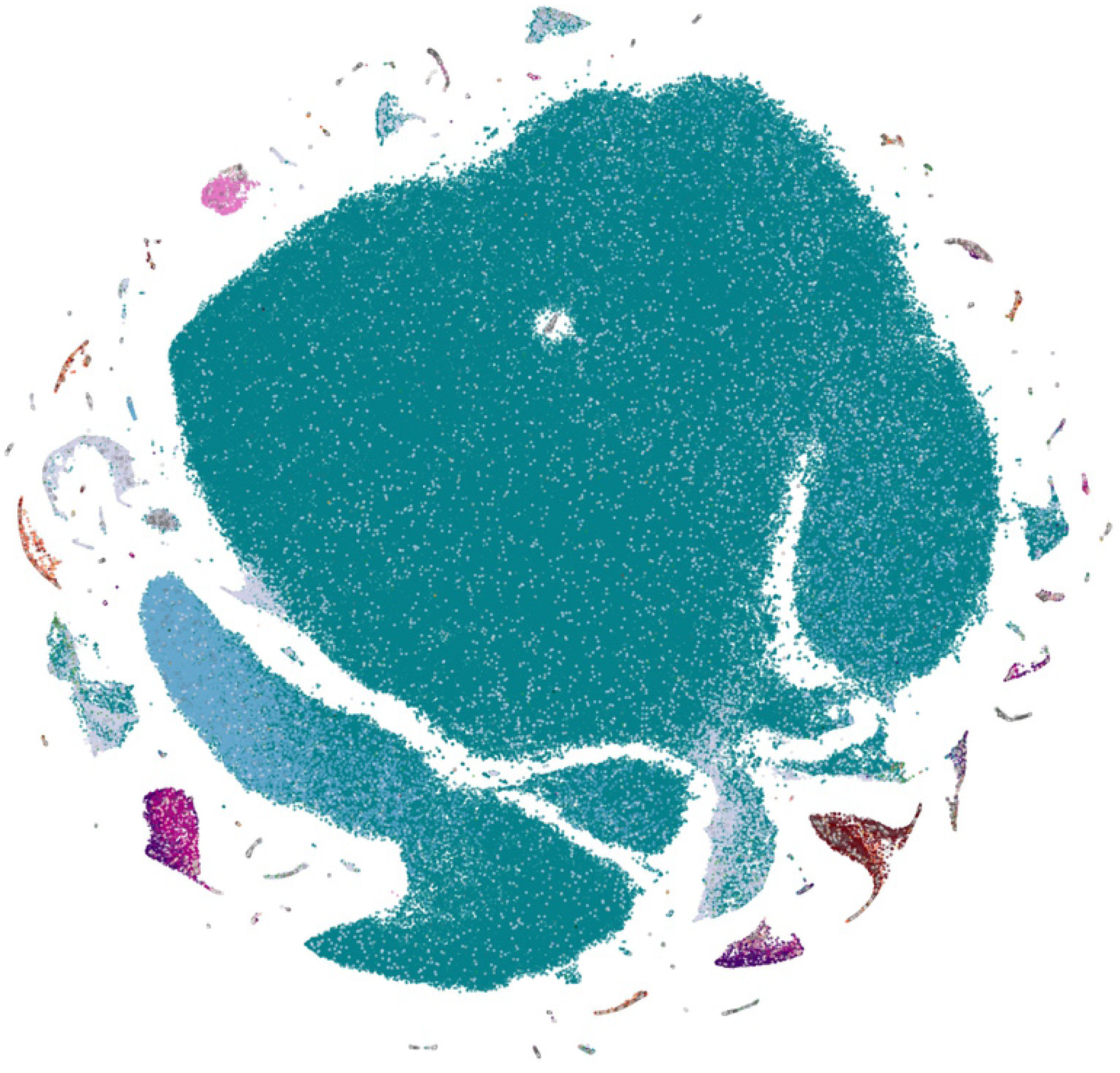
t-SNE applied to the top 10 principal components of the UKBB, colored by ethnic background. The unbalanced populations resulted in many individuals and populations being orphaned along the periphery of the main cluster.

**Fig. S41.**
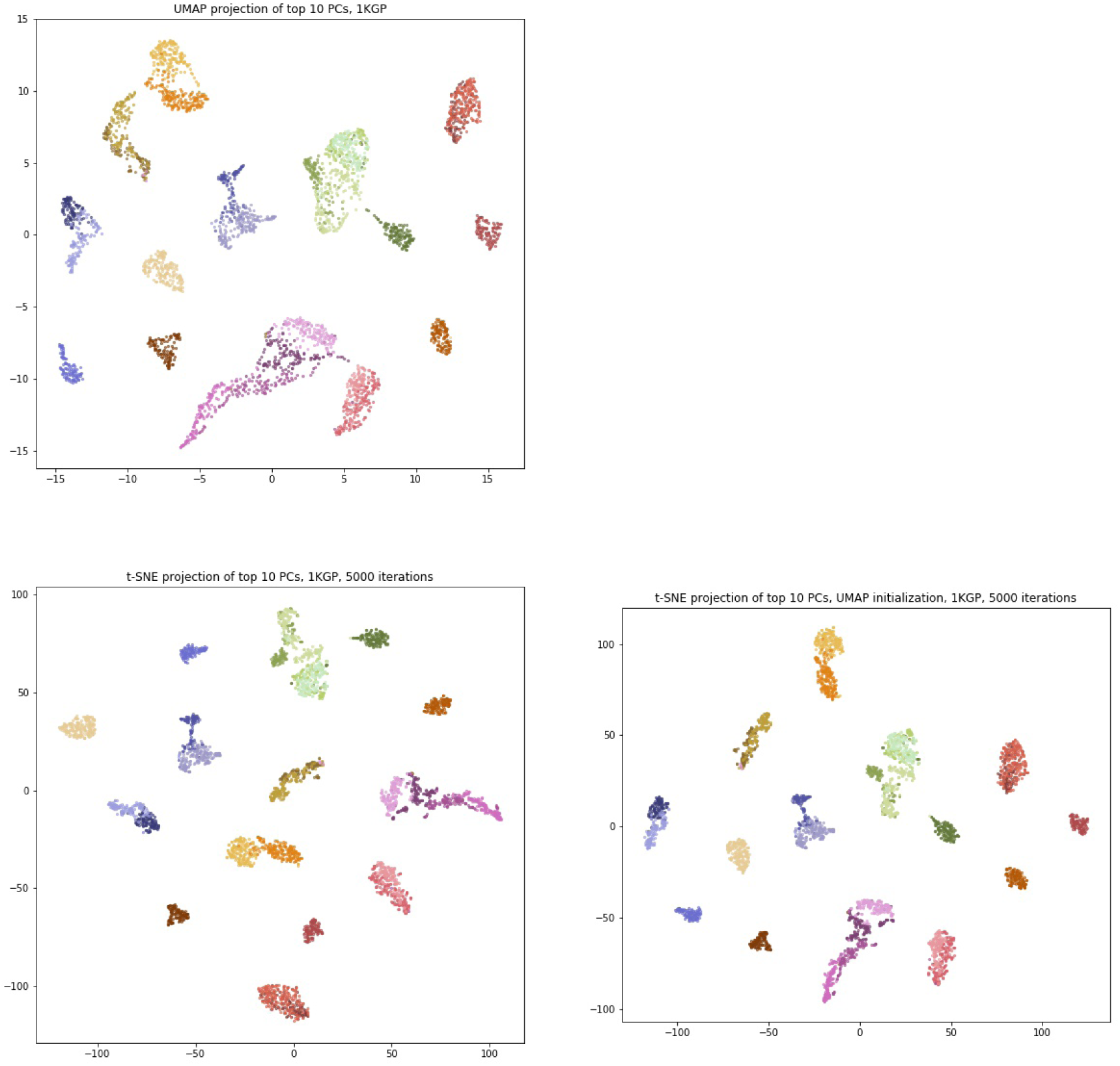
Comparing the visualizations of UMAP, standard t-SNE, and t-SNE initialized with a UMAP projection, on the top 10 principal components of the 1KGP. t-SNE used 5000 iterations.

**Fig. S42.**
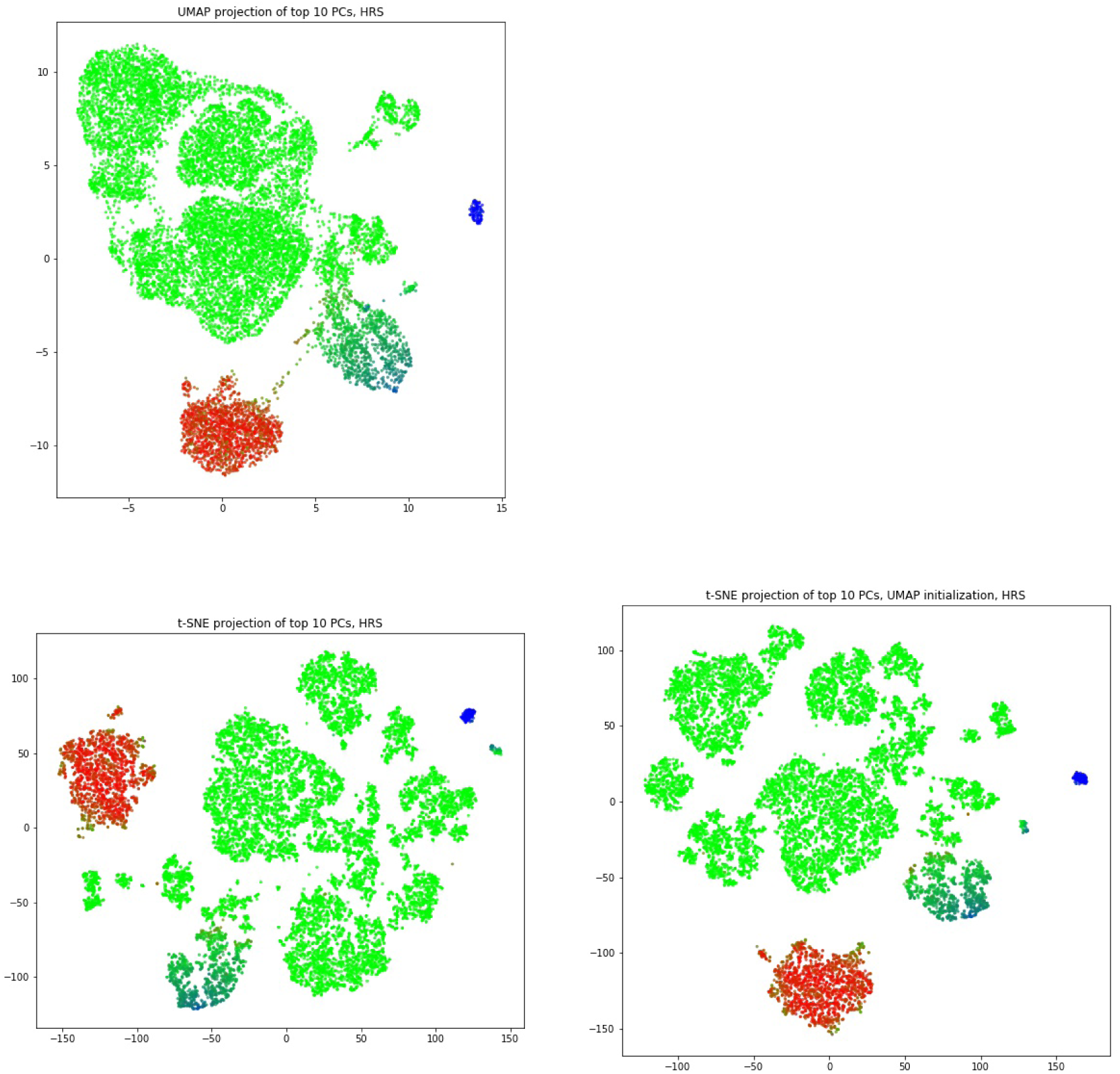
Comparing the visualizations of UMAP, standard t-SNE, and t-SNE initialized with a UMAP projection, on the top 10 principal components of the HRS. t-SNE used 5000 iterations.

**Fig. S43.**
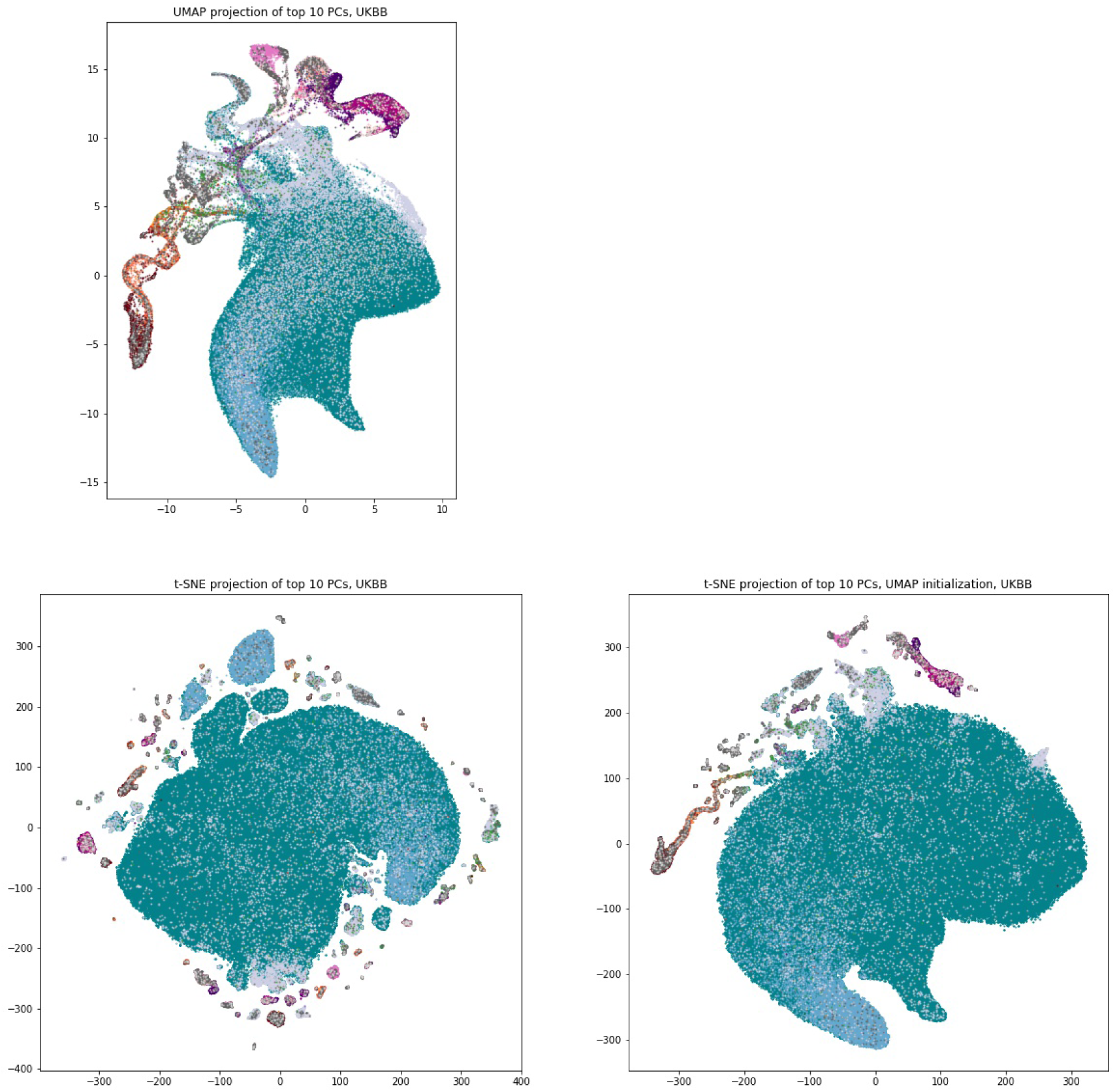
Comparing the visualizations of UMAP, standard t-SNE, and t-SNE initialized with a UMAP projection, on the top 10 principal components of the UKBB. t-SNE used 20000 iterations

**Fig. S44.**
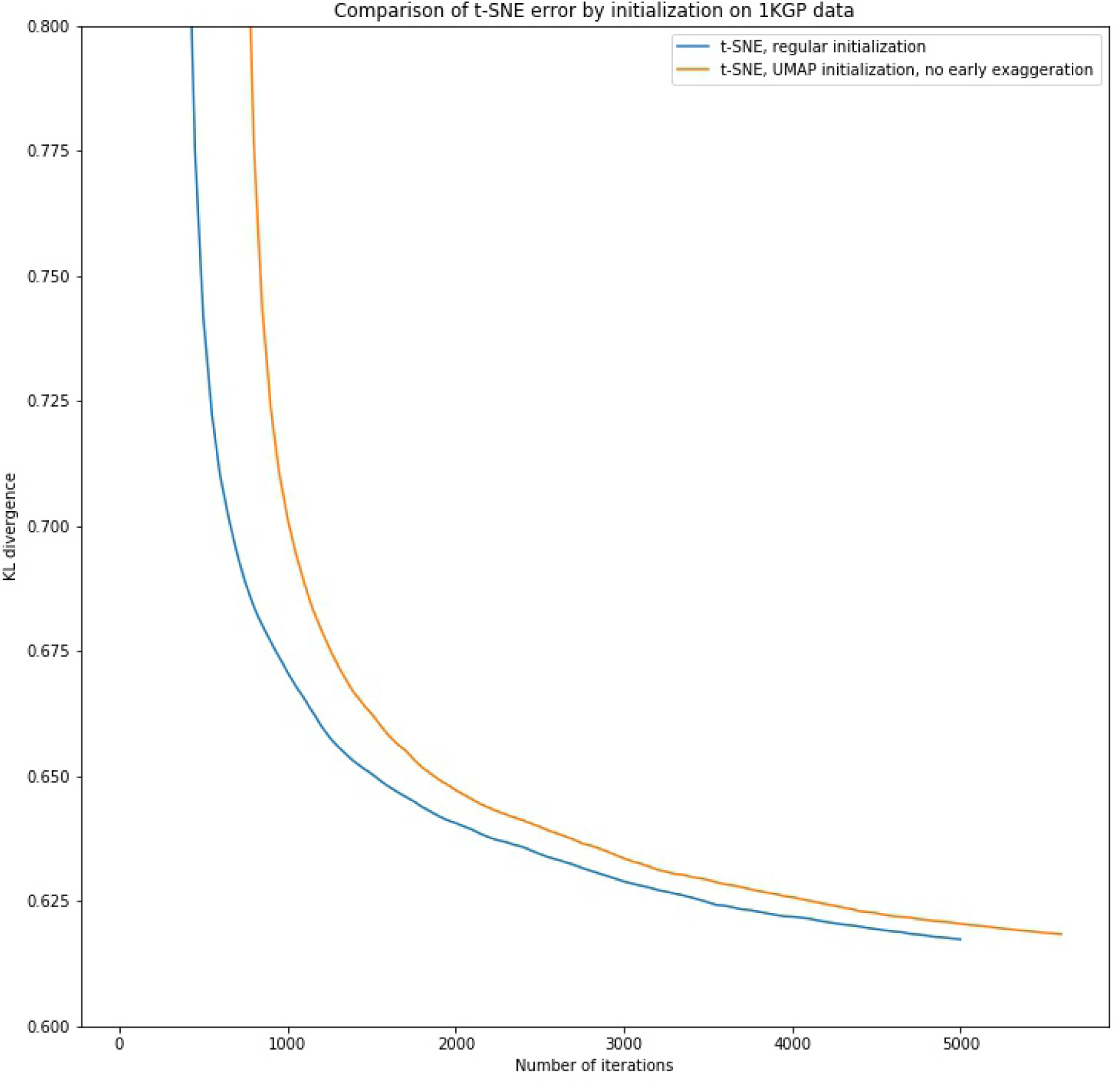
Comparing the error terms of standard t-SNE versus t-SNE initialized with a UMAP embedding and no early exaggeration. Done on the 1KGP dataset with 5000 iterations. The UMAP-initialized graph has been shifted by 600 iterations to approximate the 600 epochs UMAP uses for small datasets (*n* ≤ 10, 000).

**Fig. S45.**
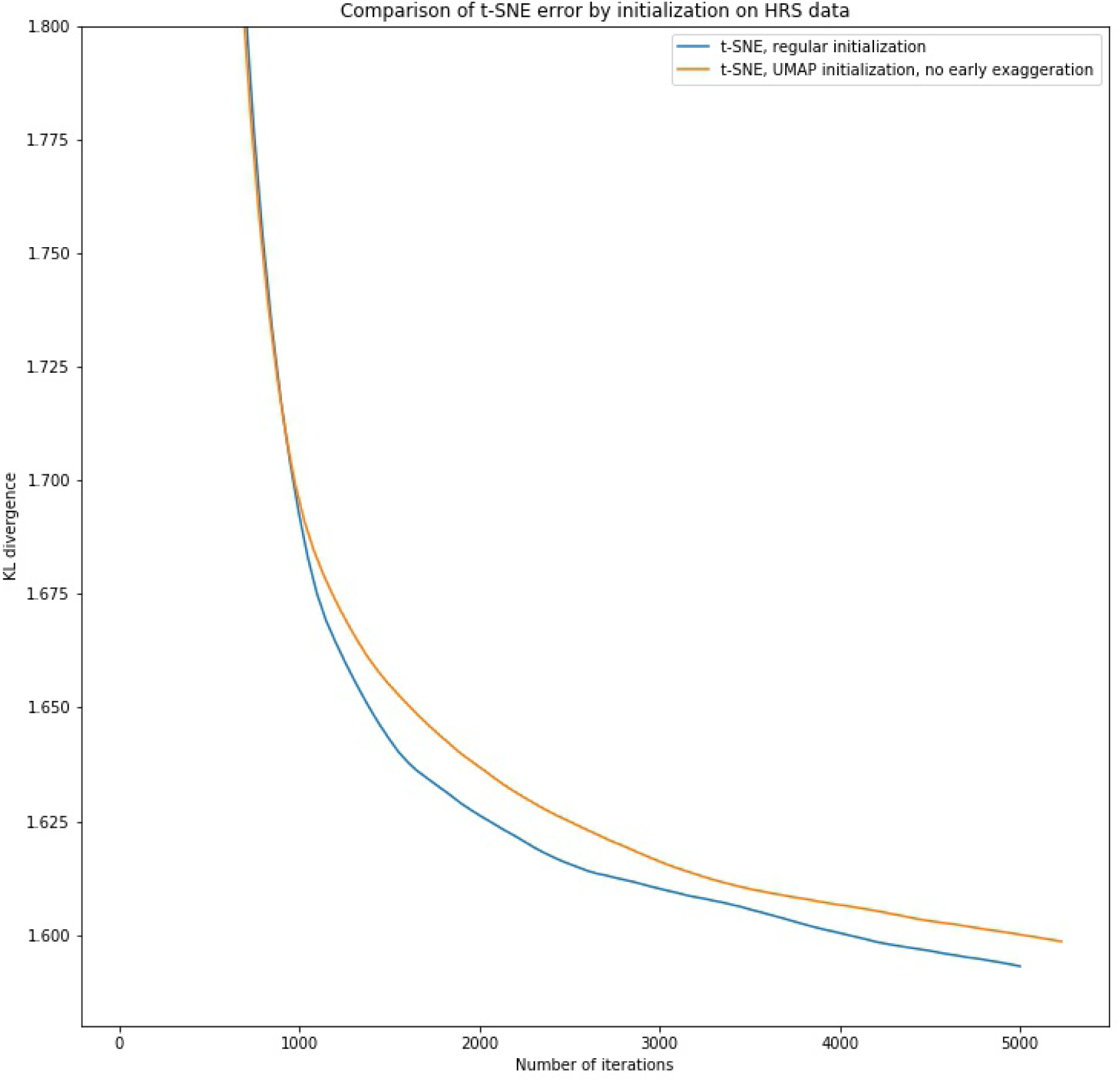
Comparing the error terms of standard t-SNE versus t-SNE initialized with a UMAP embedding and no early exaggeration. Done on the HRS dataset with 5000 iterations. The UMAP-initialized graph has been shifted by 230 iterations to approximate the 230 epochs UMAP uses for large datasets (*n* > 10, 000).

**Fig. S46.**
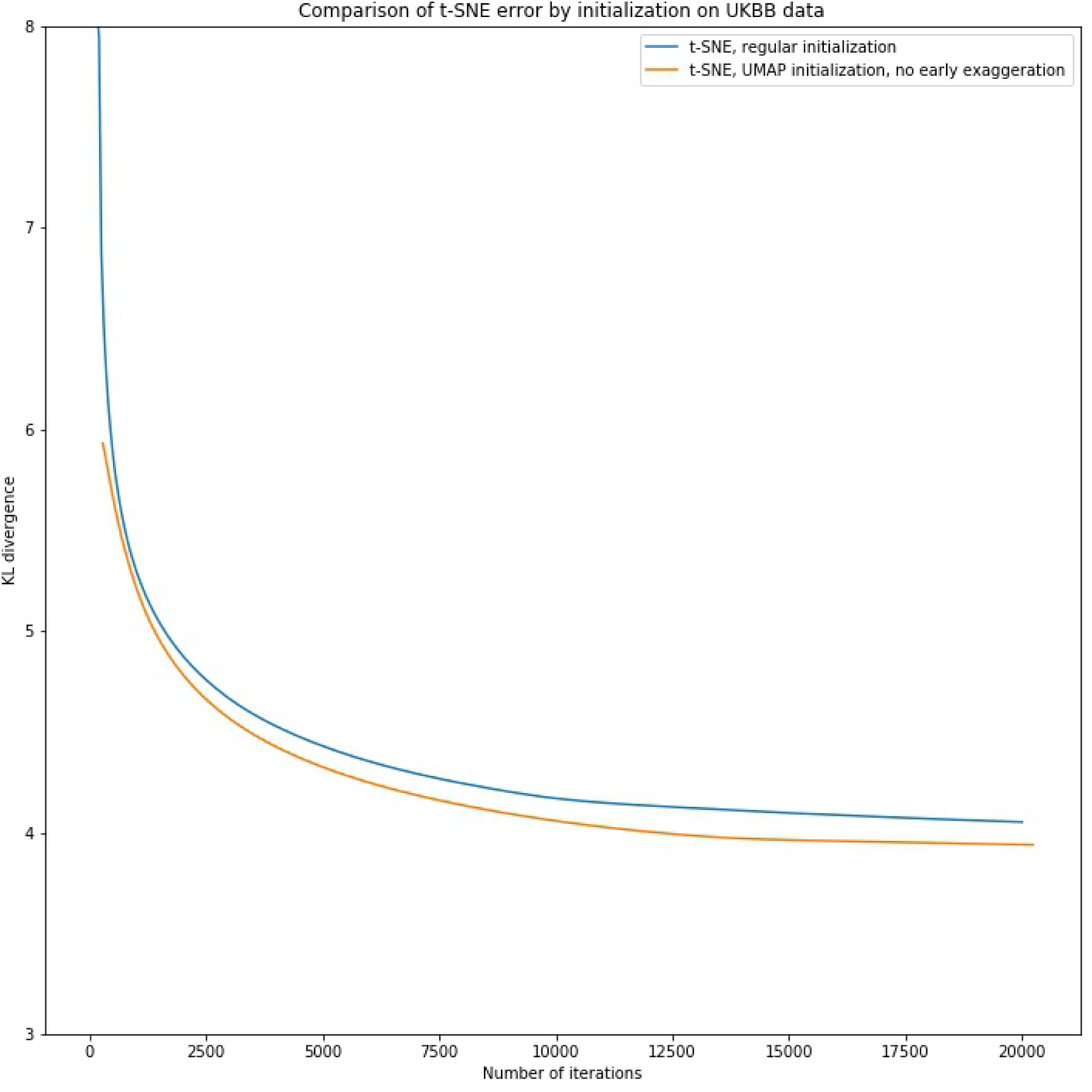
Comparing the error terms of standard t-SNE versus t-SNE initialized with a UMAP embedding and no early exaggeration. Done on the UKBB dataset with 20000 iterations. The UMAP-initialized graph has been shifted by 230 iterations to approximate the 230 epochs UMAP uses for large datasets (*n* > 10, 000).

